# Incorporating family history of disease improves polygenic risk scores in diverse populations

**DOI:** 10.1101/2021.04.15.439975

**Authors:** Margaux L.A. Hujoel, Po-Ru Loh, Benjamin M. Neale, Alkes L. Price

**Affiliations:** Department of Biostatistics, Harvard T.H. Chan School of Public Health, Boston, MA; Division of Genetics, Department of Medicine, Brigham and Women’s Hospital and Harvard Medical School; Program in Medical and Population Genetics, Broad Institute of MIT and Harvard; Department of Epidemiology, Harvard T.H. Chan School of Public Health, Boston, MA

**Author notes:** Corresponding authors: MLAH and ALP.

## Abstract

Polygenic risk scores derived from genotype data (PRS) and family history of disease (FH) both provide valuable information for predicting disease risk, enhancing prospects for clinical utility. PRS perform poorly when applied to diverse populations, but FH does not suffer this limitation. Here, we explore methods for combining both types of information (PRS-FH). We analyzed 10 complex diseases from the UK Biobank for which family history (parental and sibling history) was available for most target samples. PRS were trained using all British individuals (*N*=409K), and target samples consisted of unrelated non-British Europeans (*N*=42K), South Asians (*N*=7K), or Africans (*N*=7K). We evaluated PRS, FH, and PRS-FH using liability-scale *R*^2^, focusing on three well-powered diseases (type 2 diabetes, hypertension, depression) with *R*^2^ > 0.05 for PRS and/or FH in each target population. Averaging across these three diseases, PRS attained average prediction *R*^2^ of 5.8%, 4.0%, and 0.53% in non-British Europeans, South Asians, and Africans, confirming poor cross-population transferability. In contrast, PRS-FH attained average prediction *R*^2^ of 13%, 12%, and 10%, respectively, representing a large improvement in Europeans and an extremely large improvement in Africans; for each disease and each target population, the improvement was highly statistically significant. PRS-FH methods based on a logistic model and a liability threshold model performed similarly when covariates were not included in predictions (consistent with simulations), but the logistic model outperformed the liability threshold model when covariates were included. In conclusion, including family history greatly improves the accuracy of polygenic risk scores, particularly in diverse populations.

## Introduction

Polygenic risk scores derived from genetic data (PRS) can provide valuable information for predicting disease risk, enhancing prospects for clinical utility^1,2^. However, a well-recognized limitation of PRS methods is their poor cross-population transferability^3–7^. Family history of disease (FH) can provide complementary information about disease risk^8–11^, consistent with the rich history of leveraging data from ungenotyped but phenotyped relatives in analyses of quantitative traits in livestock^11–14^. In particular, FH has the potential to alleviate the poor cross-population transferability suffered by PRS. Combining PRS and FH information is an appealing paradigm for predicting disease risk, but it is currently unclear how to optimally combine these two sources of information. Previous studies that combined PRS and FH information restricted the PRS component to genome-wide significant loci^9,15,16^, instead of leveraging genome-wide polygenic signals; did not differentially incorporate family history for each type of relative^9,15–19^, to allow for differential environmental effects^20^ (in particular, ref. ^9^ relies on external narrow-sense heritability estimates); and did not model contributions of PRS and FH that vary as a function of the target population^9,15–17^, to optimize cross-population transferability. In addition, ref. ^9^ did not incorporate covariates; is not applicable to UK Biobank data, in which sibling history is reported as a binary variable (at least one sibling has the disease), rather than the number of affected siblings; and relies on external data to estimate model parameters. Other studies only considered PRS and FH information separately^10,21^.

Here, we develop a framework for predicting an individual’s risk of disease conditional on both their PRS and their family history of disease (PRS-FH), using either a logistic model^22^ or a liability threshold model^23^. We show via simulations and application to complex diseases from the UK Biobank^24^ that incorporating family history using PRS-FH greatly improves the accuracy of polygenic risk scores, with a particularly large improvement in diverse populations. The logistic model outperforms the liability threshold model in analyses with covariates, and we thus recommend the use of the logistic model.

## Results

### Overview of methods

We considered two PRS-FH methods based on a logistic model (PRS-FH_log_) and a liability threshold model (PRS-FH_liab_), respectively (see Methods). Both methods require a large training sample to estimate SNP effect sizes for the PRS (in this study, we use European training data), and a small additional training sample (e.g. *N*_eff_≥500; see Methods) from the target population to fit PRS-FH model parameters, which are specific to the target population. Both PRS-FH_log_ and PRS-FH_liab_ allow for sibling history to be reported as the presence or absence of at least one affected sibling (together with the total number of siblings), as in the UK Biobank. Both PRS-FH methods can be extended to incorporate covariates. We have publicly released open-source software implementing both methods as well as model parameters (specific to each target population) for both methods (see URLs).

The PRS-FH_log_ method relies on a logistic model and consists of 3 main steps (Figure 1): (1) compute PRS in all target population individuals by applying standard methods to training data; (2) use the training individuals from the target population to estimate logistic model coefficients, and (3) for each target individual, compute their predicted risk of disease, conditional on their PRS and the disease status of their first-degree relatives. In step 1, we apply BOLT-LMM^25,26^ to training data to jointly fit SNP effect sizes under a non-infinitesimal model, and compute PRS in target population individuals using these SNP effect sizes. In step 2, we estimate the contributions of the PRS and the disease status of first-degree relatives (mother, father, siblings; we allow different coefficients for each type of relative, to allow for differential environmental effects^20^) to the log-odds of disease, making a strong assumption that the log-odds of disease depends linearly on the PRS and disease status of first-degree relatives (see Discussion). These parameters are specific to the target population, requiring an extra layer of training data from the target population; in this study, we use 10-fold cross-validation in the target population. In step 3, we predict the risk of disease for each target individual as the logodds of disease based on the PRS and disease status of first-degree relatives.

**Figure 1:**
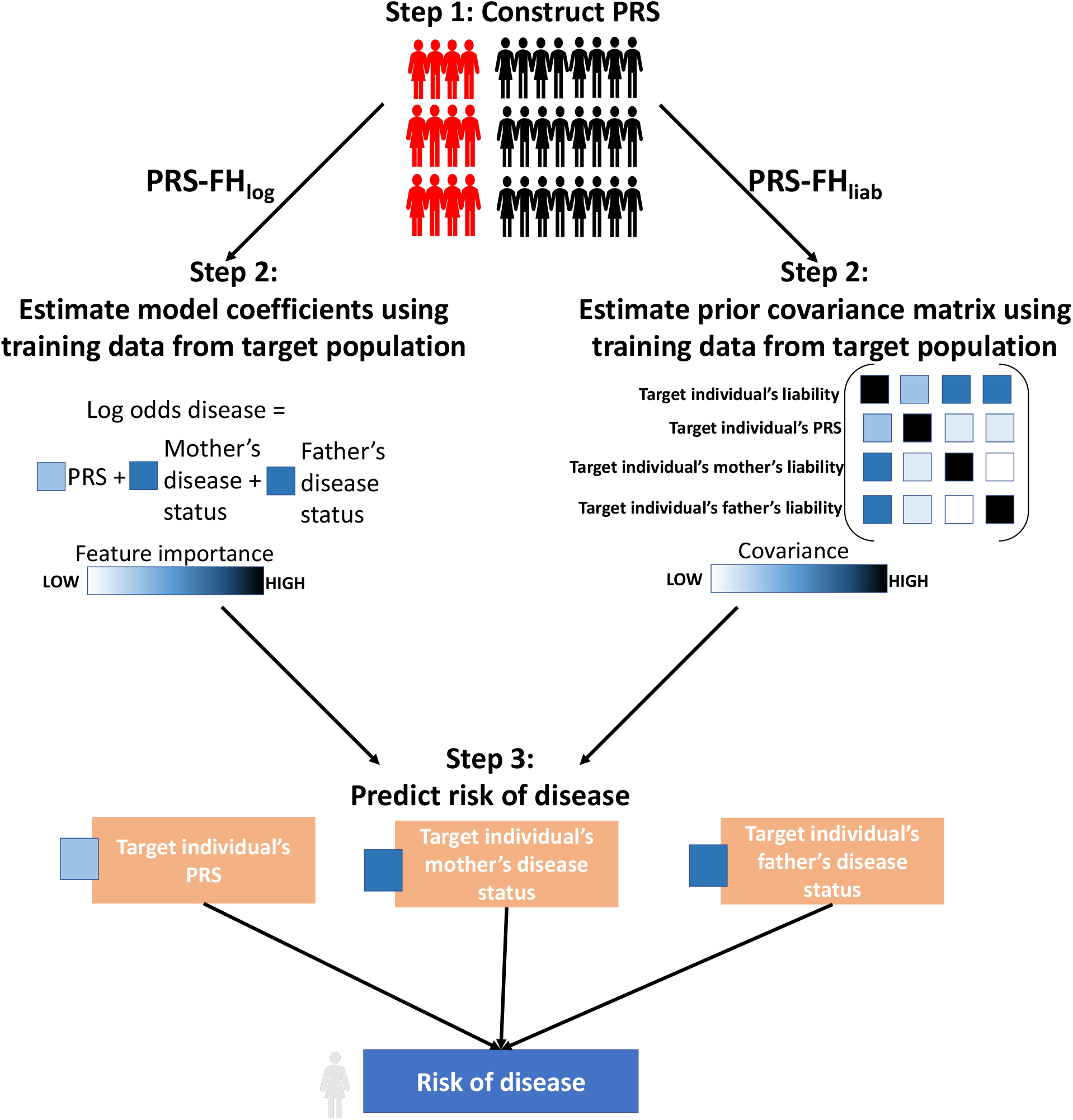
Overview of PRS-FH methods. We list the 3 steps of PRS-FH_log_ and PRS-FH_liab_. Although the PRS-FH_log_ model coefficients and PRS-FH_liab_ prior covariance shown here are the same for each parent, they may differ between mother and father. In addition, both methods can incorporate sibling history.

The PRS-FH_liab_ method relies on a liability threshold model^23^ and consists of 3 main steps (Figure 1): (1) compute PRS in all target population individuals by applying standard methods to training data; (2) use the training individuals from the target population to estimate liability threshold model parameters, and (3) for each target individual, compute their predicted risk of disease, conditional on their PRS and the disease status of their first-degree relatives. In step 1, we use BOLT-LMM^25,26^ (see above). In step 2, we estimate the variance/covariance matrix for a target individual’s total liability, their PRS, and the total liabilities of their first-degree relatives (we allow different covariances for each type of relative, analogous to above). As above, these target population-specific parameters require an extra layer of training data from the target population. In step 3, we predict the risk of disease for each target individual as the posterior probability of disease based on the PRS and disease status of first-degree relatives. The prevalence of disease (determined by the liability threshold) among target individuals may vary as a function of number of siblings, consistent with empirical data. The PRS-FH_liab_ method is conceptually related to the method of ref. ^9^, but key differences include the incorporation of genome-wide polygenic risk scores, the incorporation of different covariances for each type of relative, the way in which model parameters are estimated, the incorporation of target population-specific model parameters, and the way in which covariates are incorporated.

We compare PRS-FH_log_ and PRS-FH_liab_ to PRS alone as well as a predictor based on family history (FH) alone. The PRS method (and the PRS used within both PRS-FH methods) can employ any PRS algorithm; in this study, we use BOLT-LMM, which has been shown to attain high polygenic prediction accuracy in the UK Biobank^25,26^. The FH predictor can be constructed using a logistic model (FH_log_) or a liability threshold model (FH_liab_). Under a logistic model, the disease status of relatives linearly impacts the log-odds of disease for an individual. Under a liability threshold model, the posterior risk of disease is computed conditional on family history alone. We evaluate all methods using liability-scale *R*^2^ (ref. ^27^). We compute the standard error of liability-scale *R*^2^, and associated p-values, via a jackknife across individuals. Further details of all methods are provided in the Methods section.

### Simulations

We simulated genotypes at 100,000 unlinked SNPs for 400K unrelated PRS training samples and 40K unrelated target samples from the same population. We simulated case-control status for the PRS training samples, and case-control status plus family history (parental history for both parents) for the target samples (we did not include sibling history in these simulations); we simulated genotypes for both parents, used these to simulate genotypes for target samples (offspring), and simulated case-control status for both parents and target samples using a liability threshold model. PRS training samples and target samples were not ascertained for case-control status. Our default parameter settings involved 10,000 causal SNPs, total liability-scale heritability (*h^2^*) equal to 50%, liability-scale SNP-heritability 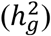 equal to 25%, and disease prevalence (*K*) equal to 1% (very low prevalence), 5% (low prevalence) or 25% (high prevalence) (implying liability threshold (*T*) equal to 2.33, 1.64, or 0.67 and total observed-scale heritability equal to 4%, 11% or 27%, respectively), with the same prevalence for parents and target samples; other parameter settings were also explored. Further details of the simulation framework are provided in the Methods section and Supplementary Table 1. We note that simulations using real LD patterns are essential for methods impacted by LD between SNPs, and that LD can impact the performance of PRS methods^1^; however, PRS-FH_log_ and PRS-FH_liab_ are not otherwise impacted by LD between SNPs, as no genotype data is used except for computing the PRS. We further note that simulations with LD using a subset of individuals from UK Biobank would not be feasible, as simulations of family history require genotypes of both target samples and relatives (in order to simulate the case-control status of both target samples and relatives), but genotypes of relatives are not available for (nearly all) UK Biobank samples.

We assessed the prediction accuracy of PRS, FH_log_, FH_liab_, PRS-FH_log_, and PRS-FH_liab_ by computing liability-scale *R*^2^ (ref. ^27^). Results are reported in Figure 2 and Supplementary Table 2. PRS attained much higher accuracy at higher prevalence, as expected due to higher observed-scale SNP-heritability (as PRS training samples were not ascertained for case-control status). FH_log_ and FH_liab_ performed similarly, and also attained higher accuracy at higher prevalence; at lower prevalence, most individuals have no affected parents, allowing little discrimination of risk based on family history. PRS and FH methods (FH_log_ and FH_liab_) performed similarly at low (5%) prevalence, but PRS outperformed FH at high (25%) prevalence-opposite to the results reported in ref. ^10^; this difference can be explained by the fact that we analyzed unascertained case-control data.

**Figure 2:**
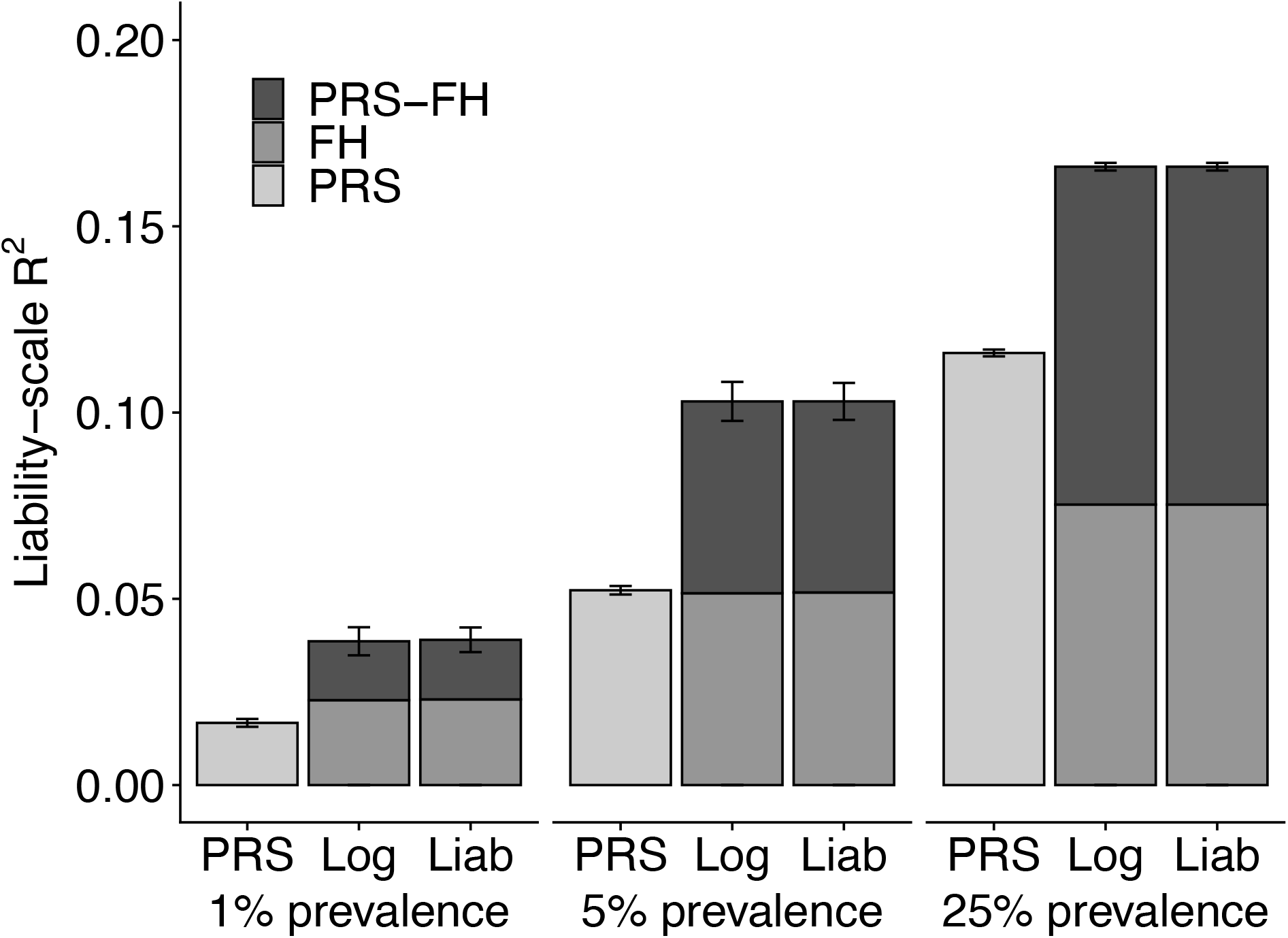
PRS-FH_log_ and PRS-FH_liab_ increase prediction accuracy in simulations. We report mean liability-scale *R*^2^ across 10 simulations for PRS alone, family history alone (FH_log_ and FH_liab_), and PRS-FH methods (PRS-FH_log_ and PRS-FH_liab_), for different values of disease prevalence. Error bars denote standard errors. Numerical results (including standard errors for FH_log_ and FH_liab_) are reported in Supplementary Table 2.

PRS-FH_log_ and PRS-FH_liab_ performed similarly, and substantially outperformed both PRS and FH methods at all prevalence values. Given that PRS-FH_liab_ makes assumptions that match the generative model used in these simulations, the strong performance of PRS-FH_log_ is supportive of the flexibility of the logistic model, even though it imposes a strong linearity assumption (on the log-odds scale). Differences in prediction *R*^2^ between PRS-FH_log_ (resp. PRS-FH_liab_) vs. PRS were smaller than the prediction *R*^2^ achieved by FH_log_ (resp. FH_liab_), due to positive correlations between PRS and FH predictions (average correlation = 0.03, 0.08, and 0.18 at very low, low, and high prevalence, respectively). Due to the poor performance (liability-scale *R*^2^ < 0.05) of all methods at very low prevalence, we restricted all further analyses to low or high prevalence.

We performed five secondary analyses. First, we assessed the calibration of each method (PRS, FH_log_, FH_liab_, PRS-FH_log_, and PRS-FH_liab_) by regressing observed disease status on the predictor (a slope of 1 implies correct calibration^28^). All methods were well-calibrated in both the low and high prevalence scenarios (Supplementary Table 3). Second, we increased the parental prevalence to twice the offspring prevalence. In these simulations, the predictive accuracy for all methods that incorporate family history (FH_log_, FH_liab_, PRS-FH_log_, and PRS-FH_liab_) increased (Supplementary Table 4). Third, we introduced environmental correlation, considering two scenarios in which the offspring had either the same or different environmental correlations with the mother and father. In both scenarios, the predictive accuracy for methods that incorporate family history (FH_log_, FH_liab_, PRS-FH_log_, and PRS-FH_liab_) increased (Supplementary Table 5). Fourth, we decreased or increased the heritability. Prediction accuracies increased with increasing heritability, but PRS-FH attained similar improvements (Supplementary Table 6). Fifth, we decreased or increased the polygenicity (number of causal SNPs) while keeping heritability constant. Prediction accuracies decreased with increasing polygenicity for methods incorporating a PRS predictor, but again PRS-FH attained similar improvements (Supplementary Table 7).

We conclude that, in these simulations, incorporating family history of disease (PRS-FH_log_ and PRS-FH_liab_) greatly increases prediction accuracy as compared to polygenic risk scores alone (PRS). We further conclude that PRS-FH_log_ and PRS-FH_liab_ generally perform similarly in these simulations. We note that the generative model in all of our simulations was the same as the liability threshold model that FH_liab_ and PRS-FH_liab_ use for prediction, and thus these simulations should be viewed as a best-case scenario for FH_liab_ and PRS-FH_liab_.

### Analysis of complex diseases from the UK Biobank

We analyzed data for 10 complex diseases from the UK Biobank^24^, consisting of genotype data, case-control status, and family history information for parents and siblings (Table 1). PRS were trained using all British individuals (*N*=409K), applying BOLT-LMM to autosomal genotyped SNPs with missingness <10% and minor allele frequency (MAF) >0.1% (672,288 SNPs). Target samples consisted of unrelated non-British Europeans (*N*=42K), South Asians (*N*=7K), or Africans (*N*=7K); target samples were unrelated to training samples and to each other (see Methods). Our primary focus was on three well-powered diseases (type 2 diabetes, depression, and hypertension) with (liability-scale) prediction *R*^2^ > 0.05 for PRS and/or FH methods in each target population; two of these diseases (type 2 diabetes, hypertension) have higher prevalence in South Asians and Africans (Table 1). We report averages across the three well-powered diseases. We also report results for each of the 10 diseases, defined as the set of diseases in the UK Biobank for which (i) family history (parental and sibling history) was available for most target samples and (ii) prediction *R*^2^ was statistically significant (after Bonferroni correction) for PRS and/or FH methods in the largest target population (non-British Europeans; Supplementary Table 8).

**Table 1:**
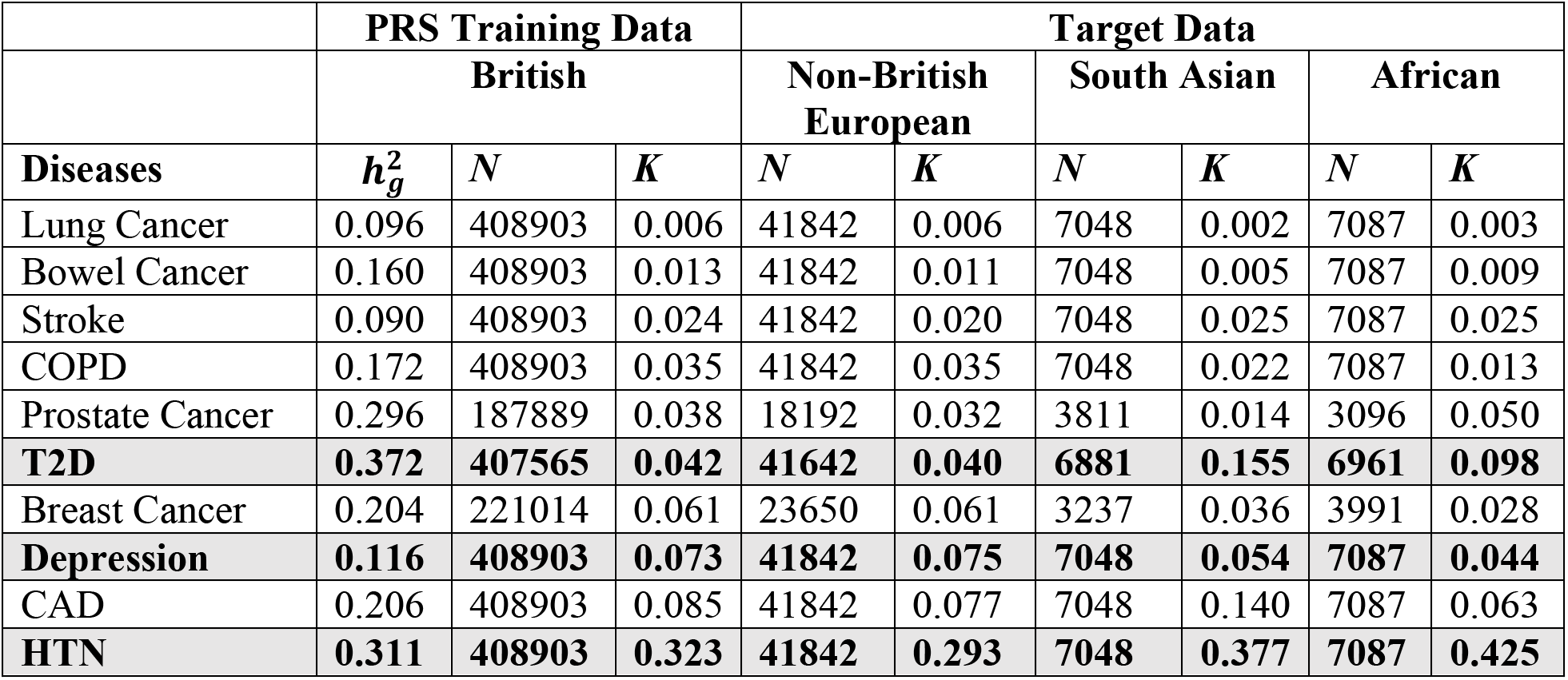
List of 10 UK Biobank diseases analyzed. For each disease, we report the SNP-heritability 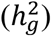 in UK Biobank British training data and the number of samples (*N*) and disease prevalence (*K*) in each UK Biobank training and target population. We note that the sample size and prevalence in British training data includes information from related individuals, but SNP-heritability was estimated using unrelated British individuals. Diseases are listed in order of disease prevalence in British training data. Our primary focus was on three well-powered diseases (type 2 diabetes, depression, and hypertension; denoted in bold) with (liability-scale) prediction *R*^2^ > 0.05 for PRS and/or FH in each target population. COPD, chronic obstructive pulmonary disease, defined as chronic bronchitis/emphysema; T2D, type 2 diabetes; CAD, coronary artery disease; HTN, hypertension.

We assessed the prediction accuracy of PRS, FH_log_, FH_liab_, PRS-FH_log_ and PRS-FH_liab_. Results are reported in Figure 3a and Supplementary Table 9. Across the three well-powered diseases, PRS attained average prediction *R*^2^ of 5.8%, 4.0%, and 0.53% in non-British Europeans, South Asians, and Africans, confirming poor cross-population transferability^3–7^. In contrast, FH_log_ attained similar prediction *R*^2^ across populations: 8.0%, 8.6% and 9.6%, with similar results for FH_liab_. Notably, PRS-FH_log_ attained average prediction *R*^2^ of 13%, 12%, and 10%, with similar results for PRS-FH_liab_. Thus, PRS-FH_log_ and PRS-FH_liab_ attained a large relative improvement vs. PRS in Europeans (consistent with simulations) and an extremely large relative improvement vs. PRS in Africans. For each disease and each target population, the difference between PRS-FH_log_ (or PRS-FH_liab_) and PRS was highly statistically significant (*p* < 2 × 10^-6^). Differences in prediction *R*^2^ between PRS-FH_log_ (or PRS-FH_liab_) and PRS were generally slightly smaller than the prediction *R*^2^ attained by FH, due to slight correlations between PRS and FH predictions (average = 0.07 across the three well-powered diseases and 0.05 across all 10 diseases; Supplementary Figure 1, Supplementary Table 10). Parameters estimated by PRS-FH_log_ and PRS-FH_liab_ are reported in Supplementary Table 11. Across the three well-powered diseases, sibling history was assigned higher weight than parental history regardless of target population (likely due to differential shared environmental effects^20^), whereas the weight assigned to PRS depended on the target population.

**Figure 3:**
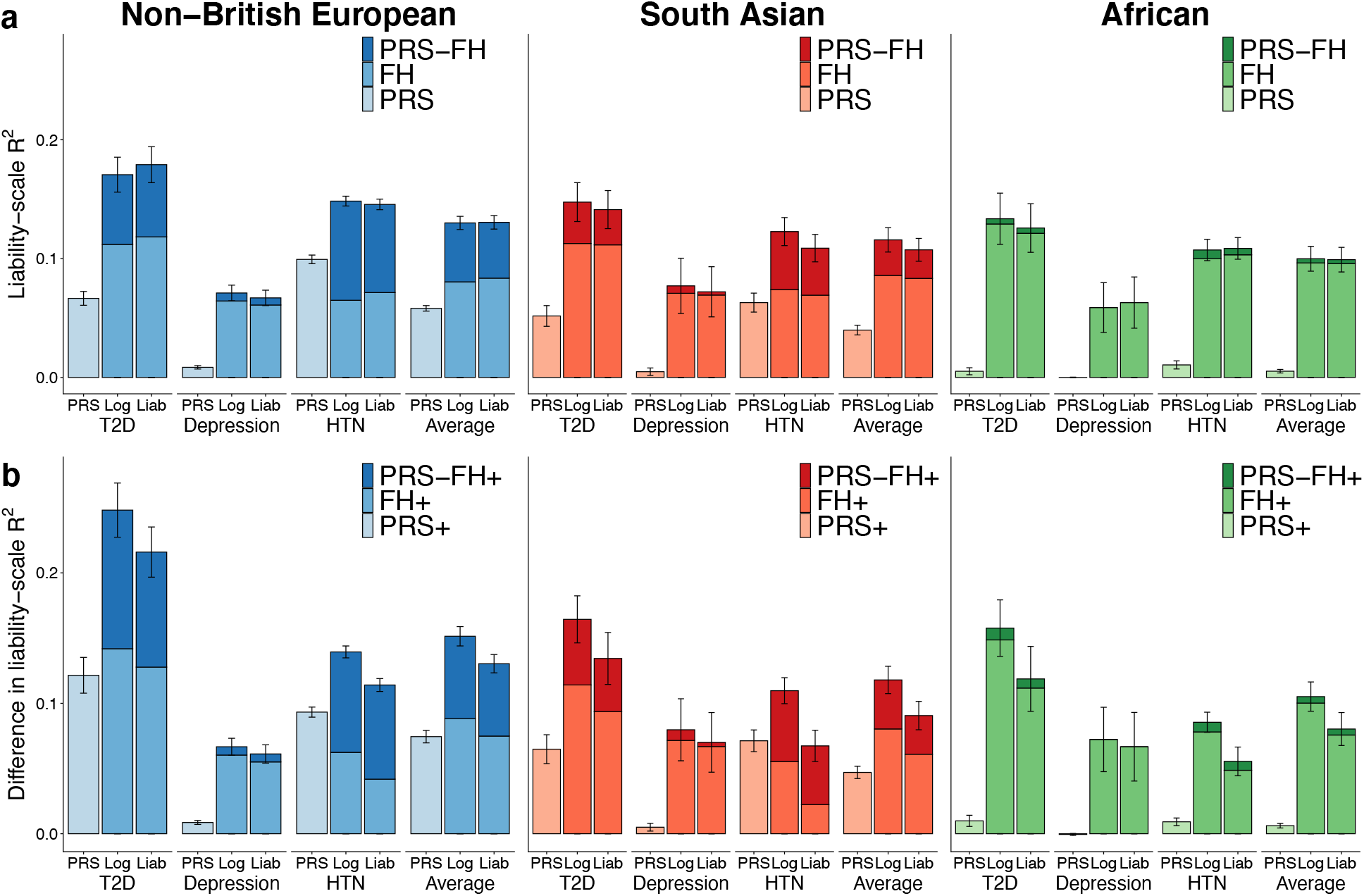
PRS-FH increases prediction accuracy in analyses of UK Biobank diseases. (a) Analyses without covariates. We report liability-scale *R*^2^ for PRS alone, family history alone (FH_log_ and FH_liab_), and PRS-FH methods (PRS-FH_log_ and PRS-FH_liab_), for different diseases and target populations. (b) Analyses with covariates. We report difference in liability-scale *R*^2^ (see text) for the corresponding methods incorporating covariates (PRS^+^, FH^+^, PRS-FH^+^), for different diseases and target populations. We focus on three well-powered diseases with *R*^2^ > 0.05 for PRS and/or FH in each target population. Numerical results (including standard errors for FH_log_ and FH_liab_) are reported in Supplementary Table 9 and Supplementary Table 20.

More broadly, PRS-FH_log_ (and PRS-FH_liab_) consistently attained higher prediction *R*^2^ than PRS across all 10 diseases (Supplementary Table 9). The prediction accuracy of PRS increased as a function of observed-scale SNP-heritability (which is partly determined by prevalence) (Supplementary Figure 2), and the prediction accuracy of FH increased as a function of both the covariance between liabilities of target samples and first-degree relatives (which is largely determined by total narrow-sense heritability) and the prevalence in first-degree relatives (Supplementary Figure 2, Supplementary Tables 4 and 6). The correlations between PRS and FH predictions were low for all diseases (−0.02 to 0.13), but increased as a function of prevalence and SNP-heritability (Supplementary Figure 1, Supplementary Table 10).

We performed eight secondary analyses. First, we assessed the calibration of each method. We determined that PRS-FH_log_ attained better calibration than PRS-FH_liab_ (average regression slope of 0.93 vs. 0.60 across 3 well-powered diseases; Supplementary Table 12). Second, we decreased the number of training samples from the target population used to fit PRS-FH model parameters below its default level (which is based on 10-fold cross-validation; see Overview of methods). For both PRS-FH_log_ and PRS-FH_liab_, the number of training samples from the target population had little impact on predictive accuracy for values of *N*_eff_≥500 (Supplementary Figure 3). Third, we compared the performance of both FH and PRS-FH methods when incorporating parental history only vs. both parental and sibling history. Incorporating both parental and sibling history attained moderately higher predictive accuracy (Supplementary Table 13). Fourth, we assessed the performance of a simplified logistic regression-based method that used a single binary independent variable for overall (parental and sibling) family history. We determined that PRS-FH_log_ attained significantly higher prediction accuracy than this method (Supplementary Table 14). Fifth, we assessed the potential benefit to FH_log_ and PRS-FH_log_ of including in the logistic model an interaction term between number of siblings and sibling history. We determined that there was no significant benefit (Supplementary Table 15). Sixth, we assessed whether FH_log_ and PRS-FH_log_ would benefit from incorporating the total number of siblings of each target individual using indicator variables in addition to a continuous variable. We determined that disease prevalence empirically varied non-linearly as a function of the number of siblings (which is known to correlate with socioeconomic factors) (Supplementary Table 16), and that accounting for this generally produced non-significant improvements (Supplementary Table 17). Seventh, we assessed whether FH_liab_ and PRS-FH_liab_ benefit from allowing the prevalence of disease (determined by the liability threshold) among target individuals to vary as a function of the number of siblings. We determined that FH_liab_ and PRS-FH_liab_ attained slightly higher prediction accuracy than corresponding methods that do not allow the prevalence of disease to vary as a function of the number of siblings (Supplementary Table 18); we elected to allow the primary FH_liab_ and PRS-FH_liab_ methods to benefit from this information as a conservative choice, as they were not ultimately the methods of choice (see below). Eighth, for each of the 5 methods, we evaluated the prevalence of disease in each percentile of predicted disease risk^2^. We confirmed that PRS-FH_log_ and PRS-FH_liab_ also performed best under this metric (Supplementary Figure 4).

We conclude that incorporating family history of disease (PRS-FH_log_ and PRS-FH_liab_) greatly increases prediction accuracy as compared to polygenic risk scores alone (PRS), particularly in Africans. We further conclude that PRS-FH_log_ and PRS-FH_liab_ generally perform similarly in analyses without covariates.

### Incorporation of covariates in UK Biobank analyses

We repeated the analyses of 10 complex diseases from the UK Biobank by incorporating covariates into each method: PRS^+^, FH^+^_log_, FH^+^_liab_, PRS-FH^+^_log_ and PRS-FH^+^_liab_; the covariates included age, sex, BMI and 20 principal components (see Methods). PRS^+^ incorporates covariates by training a logistic model with PRS and all covariates. FH^+^_log_ and PRS-FH^+^_log_ incorporate covariates by including them as independent variables in the logistic model. FH^+^_liab_ and PRS-FH^+^_liab_ incorporate covariates by estimating a disease threshold for the liability (exclusive of covariates) that varies based on the covariates (see Methods). We evaluated the predictive accuracy of each method using difference in liability-scale *R*^2^ (defined as liability-scale *R*^2^ minus the liability-scale *R*^2^ attained using covariates alone). As above, our primary focus was on the three well-powered diseases (type 2 diabetes, depression, and hypertension); the impact of covariates on these diseases was substantial, as covariates alone attained average prediction *R*^2^ of 20%, 17%, and 15% in non-British Europeans, South Asians, and Africans, with most of the prediction *R*^2^ contributed by age and BMI (Supplementary Table 19).

We assessed the prediction accuracy of PRS^+^, FH^+^_log_, FH^+^_liab_, PRS-FH^+^_log_ and PRS-FH^+^_liab_. Results are reported in Figure 3b and Supplementary Table 20. Across the three well-powered diseases, PRS^+^ attained average prediction accuracy (difference in liability-scale *R*^2^) of 7.4%, 4.7%, and 0.62% in non-British Europeans, South Asians, and Africans, again reflecting poor cross-population transferability^3–7^. In contrast, FH^+^_log_ attained similar prediction accuracy across populations: 8.8%, 8.0% and 10%; results were also similar across populations for FH^+^_liab_. Notably, PRS-FH^+^_log_ outperformed PRS-FH^+^_liab_, with prediction accuracies of 15%, 12%, and 11% for PRS-FH^+^_log_ in the three populations vs. 13%, 9.1%, and 8.0% for PRS-FH^+^_liab_ (most differences were statistically significant: *p*=0.0001-0.0007 for T2D, *p*=0.05-0.6 for depression, *p*=6×10^-28^-5×10^-9^ for HTN); similarly, FH^+^_log_ outperformed FH^+^_liab_. We note that PRS-FH^+^_log_ and FH^+^_log_ model the effects of family history and covariates jointly, whereas PRS-FH^+^_liab_ and FH^+^_liab_ model the effects of covariates marginally (see Methods); as both family history and PRS is correlated with covariates (Supplementary Table 21), this may explain the better performance of PRS-FH^+^_log_ and FH^+^_log_. The differences in prediction *R*^2^ attained by PRS^+^, FH^+^_log_, FH^+^_liab_, PRS-FH^+^_log_ and PRS-FH^+^_liab_ vs. a prediction model based on covariates alone were generally similar to the absolute predictive *R*^2^ attained by PRS, FH_log_, FH_liab_, PRS-FH_log_, and PRS-FH_liab_, with limited exceptions (Supplementary Table 9 and 20). Surprisingly, the relative prediction accuracy of PRS^+^ was sometimes larger than the prediction accuracy of PRS alone, which is mathematically possible under a logistic model. The pairwise correlations between PRS, FH_log_, FH_liab_, and a prediction based on covariates alone ranged from −0.06 to 0.16 (Supplementary Table 21).

We performed three secondary analyses. First, we assessed the calibration of each method. We determined that PRS-FH^+^_log_ attained better calibration than PRS-FH^+^_liab_ (average regression slope of 0.92 vs 0.68 across 3 well-powered diseases; Supplementary Table 22). Second, we compared the performance of both FH^+^ and PRS-FH^+^ methods when incorporating parental history only vs. both parental and sibling history. Incorporating both parental and sibling disease history attained moderately higher predictive accuracy for FH^+^_log_ and PRS-FH^+^_log_, but results were mixed for FH^+^_liab_ and PRS-FH^+^_liab_ (Supplementary Table 23). Third, we assessed the performance of a simplified logistic regression-based method (incorporating covariates) that used a single binary independent variable for overall (parental and sibling) family history. We determined that PRS-FH^+^_log_ attained significantly higher prediction accuracy than this method (Supplementary Table 24; analogous to Supplementary Table 14).

We conclude that when covariates are included in the predictions, incorporating family history of disease (PRS-FH^+^_log_ and PRS-FH^+^_liab_) continues to greatly increase prediction accuracy as compared to polygenic risk scores alone (PRS^+^), particularly in Africans. We further conclude that PRS-FH^+^_log_ outperforms PRS-FH^+^_liab_ in analyses with covariates.

## Discussion

We have explored methods for combining polygenic risk scores and family history (PRS-FH) to predict risk of disease, using a logistic model or a liability threshold model. We determined that PRS-FH greatly increases prediction accuracy as compared to polygenic risk scores alone across a broad set of simulations and empirical analyses, including analyses incorporating covariates. We recommend the use of the logistic model, which outperforms the liability threshold model in analyses with covariates (however, we note that the liability threshold model has proven valuable in other settings^23,29–33^). The increase in prediction accuracy attained by PRS-FH is particularly large in diverse populations (e.g. Africans), suggesting that PRS-FH will be a method of choice for closing the well-documented gap in disease risk prediction accuracy in diverse populations^3–7^. Our findings emphasize the value of collecting and incorporating family history data, as well as data on clinical covariates, whenever it is practical to do so.

PRS-FH differs from previous approaches for combining PRS and FH information^9,15–19^ in several key ways. First, previous methods restricted the PRS component to genome-wide significant loci, but PRS-FH leverages genome-wide polygenic signals, which have higher predictive value^1^. Second, previous methods incorporate each type of relative equally, but PRS-FH incorporates each type of relative separately, to allow for differential environmental effects^20^. In particular, a recent study that used a single binary independent variable for overall (parental and sibling) family history reported no significant improvement from incorporating family history in prostate cancer analyses of UK Biobank Europeans^34^ (AUC = 0.836 vs. 0.833; analogous to Supplementary Table 14), whereas PRS-FH^+^_log_ attained a significant improvement from incorporating family history in prostate cancer analyses of UK Biobank Europeans (*R*^2^ = 0.100 vs. 0.069, *p*=0.029, Supplementary Table 9; *p*=0.0035 in analyses with covariates, Supplementary Table 20). Third, previous methods do not allow the contributions of PRS and FH to vary as a function of the target population, but PRS-FH optimizes these contributions as a function of the target population, increasing prediction accuracy in diverse populations. Fourth, ref.^9^ did not incorporate covariates; furthermore, in our work all effects of family history and covariates are modeled jointly, in contrast to ref. ^16^ (and the approach for incorporating covariates discussed as a future direction in ref.^9^). Fifth, ref. ^9^ is not applicable to UK Biobank data, in which sibling history is reported as a binary variable (at least one sibling has the disease). Sixth, PRS-FH differs from ref. ^9^ in that model parameters are estimated within the target population of interest, rather than relying on external data sources.

Although PRS-FH greatly increases prediction accuracy, it has several limitations. First, PRS-FH requires an additional layer of training data from the target population in order to optimize the contributions of PRS and FH to the target population. However, this requires only a small number of training samples from the target population (e.g. *N*_eff_≥500; see Supplementary Figure 3), and the additional training step can be omitted for the diseases and target populations that we have analyzed here (for which these model parameters are reported in Supplementary Table 11). Second, the logistic model makes a strong assumption that the log-odds of disease depends linearly on the PRS and disease status of first-degree relatives—an assumption that lacks a strong theoretical justification. However, simulations and empirical results are strongly supportive of the practical ramifications of this assumption. Third, family history may reflect a different underlying genetic architecture than case-control status, for example, due to differences in the etiology of early-onset versus late-onset disease or differences in diagnostic criteria over time; however, we previously reported very high genetic correlations between case-control and family history phenotypes in the UK Biobank^33^. Fourth, self-reported family history information may be inaccurate. However, we previously determined that self-reported family history is reasonably accurate in the UK Biobank (~80% correlation between true and self-reported family history, based on sibling concordance^33^); the imperfect accuracy is explicitly accounted for by PRS-FH model parameters, and incorporating self-reported family history clearly improves prediction accuracy in our study. Fifth, we did not perform analyses in which we trained and validated in different cohorts. We anticipate that this will become possible in the future with the emergence of large biobanks collecting a rich set of phenotypes including family history^35,36^. Finally, incorporating training data from auxiliary traits has substantial potential to improve polygenic prediction accuracy^37,38^, but was not implemented in our study; incorporating auxiliary traits into PRS-FH is straightforward under the PRS-FH framework, and remains as a future research direction. Despite these limitations, we anticipate that PRS-FH will attain large increases in prediction accuracy in future studies, particularly in diverse populations.

## Acknowledgements

We are grateful to Omer Weissbrod for helpful discussions. This research was funded by NIH grants R01 HG006399 (A.L.P.), R01 MH101244 (A.L.P.), R37 MH107649 (B.N. and A.L.P.) and 5T32CA009337-32 (M.L.A.H.). P.-R.L. was supported by the Next Generation Fund at the Broad Institute of MIT and Harvard and a Sloan Research Fellowship. This research was conducted using the UK Biobank resource under application no. 16549.

## URLs

PRS-FH software: https://data.broadinstitute.org/alkesgroup/UKBB/PRSFH/.

## Methods

### PRS-FH_log_ method

The PRS-FH_log_ method models the PRS and the disease status of relatives as linearly impacting the log-odds of disease for an individual, as detailed below.

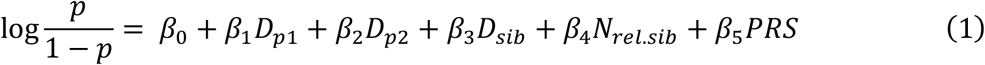

where *D*_*p*1_, *D*_*p*2_, and *D_sib_* are the binary disease status variables for an individual’s parents and siblings, respectively, *N_rel.sib_* is the number of relevant siblings of an individual (number of total siblings for non-sex-specific diseases, number of sisters for breast cancer, and number of brothers for prostate cancer), and *PRS* is the individual’s PRS. An individual’s PRS is constructed as a weighted sum of their genotypes:

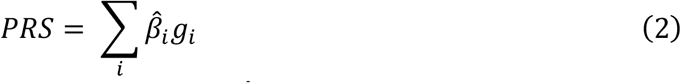

where *g_i_* are an individual’s genotypes at SNP *i* (0,1,2) and 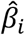 are the per-allele effect sizes of SNP *i* estimated using training data. We note that there are multiple algorithms for constructing PRS, however the construction of PRS is not the primary focus of this work.

We considered a logistic model incorporating the PRS (as a continuous covariate), the 3 binary indicators for the disease status of mother, father, and siblings, and a continuous covariate for the number of relevant siblings (see equation (1)). We elected not to use indicator variables for the number of total siblings an individual has (e.g. 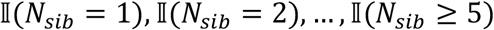) in the primary method as this generally produced non-significant improvements (Supplementary Table 17) and increased model complexity.

### PRS-FH_liab_

The PRS-FH_liab_ method models the family history of disease and PRS using a liability threshold model^23^. The liability threshold model assumes an individual has an underlying liability, *ϵ*, which is normally distribution with a mean of 0 and variance of 1. An individual is a case (*z* = 1) if and only if *ϵ* ≥ *T* otherwise the individual is a control (*z* = 0). *T* determines the disease prevalence *K*; *K* = 1 — Φ(*T*) where Φ(*T*) is the normal cumulative distribution function, i.e. Φ(*T*) = Pr (*N*(0,1) ≤ *T*).

We assume a multivariate normal distribution for the individual’s liability, the individual’s PRS, and the target individual’s relatives’ liabilities. For example, to incorporate the individual’s PRS as well as the parental and a sibling’s disease history we assume,

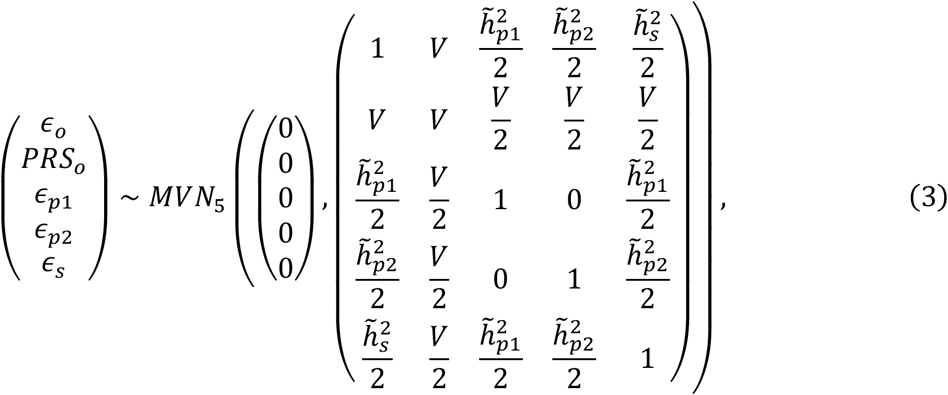

where *ϵ_o_* is the total liability of the target individual, *PRS_o_* is the PRS of the target individual, and *ϵ*_*p*1_, *ϵ*_*p*2_, and *ϵ*_*s*_ are the liabilities of the parents and the sibling, respectively, 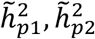, and 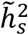 are the pseudo-heritabilities of the disease on the liability scale of the parents and the sibling, respectively, and *V* is the amount of variance the PRS can explain on the liability scale. The pseudo-heritabilities of the disease reflect a combination of heritability and shared environmental effects (which may vary across classes of relatives), and can be estimated using maximumlikelihood methods (see Supplementary Note and Supplementary Table 25 for justification of pseudo-heritability and details on its estimation). We can estimate the variance explained by the PRS on the liability scale as

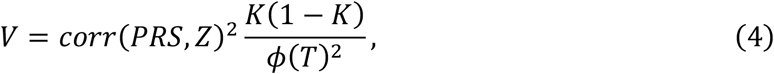

where *K* is the disease prevalence, Z is the disease status, T = Φ^−1^(*K*), and *ϕ* is the normal probability density function. This estimate of *V* is similar to previous derivations converting between the observed-scale and the liability-scale (see Supplementary Note)^29^. After estimating the liability-scale variance explained by the PRS (*V*), the raw PRS is scaled to have mean zero and the desired variance prior to being utilized by PRS-FH_liab_. Setting 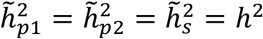 models PRS and family history of disease assuming no environmental correlation.

Using the distribution shown in equation (1), we can compute the posterior mean and variance of *ϵ_o_*, conditional on the individual’s PRS and the disease status of family members (e.g. if parent 1 is a case we can condition on *ϵ*_*p*1_ ≥ *T*_*p*1_). Given the mean and variance of the posterior distribution, denoted *μ*_*ϵ_o_*|_· and 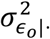 respectively, we assume normality and compute the posterior risk of disease for an individual to be:

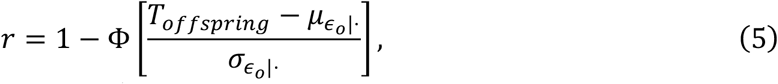

where *T_offspring_* is either Φ^−1^(*K_offspring_*) or a function of covariates, depending on the model being implemented (see below). We note that the posterior risk of disease is distinct from the posterior mean (and variance) of liability. We elected to use the posterior risk of disease, rather than simply the posterior mean or variance, as this appropriately weights the mean and variance of the liability for disease. Conditioning on family history will result in a non-normal distribution, however, this deviation from normality is generally small^9,23^. We elected to have *T_offspring_* depend on the number of total siblings an individual has through the use of indicator variables (e.g. 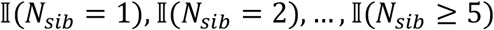) as a conservative choice as it optimized the performance of PRS-FH_liab_ (Supplementary Table 18), which ultimately was not the recommended method.

We use the Pearson-Aitken formula, as well as properties about truncated normal distributions, to compute posterior distributions^9,39^. Sibling history is reported as a binary condition in UK Biobank; individuals report whether at least one or none of their siblings are affected. For individuals who report at least one of their siblings is affected, the posterior mean and variance is estimated analytically (see Supplementary Note).

Missing family history for some relatives is not an issue as the posterior distribution is computed conditional on known information, and therefore the number of relatives being modeled is reduced when missing family history exists. We use estimates of disease prevalence which differ for mother, father, and offspring as well as estimates of pseudo-heritability which differ for mother and father.

### Simulations

We simulated genotypes at 100,000 unlinked SNPs and case-control status for 400,000 unrelated training samples. For computational simplicity we generated 10 genotype matrices for the training data and given these genotype matrices, can then generate 10 different case-control vectors represent different scenarios of ranging prevalence, 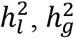, and polygenicity (number of causal SNPs). To obtain PRS, we computed prediction *β* for all 100,000 unlinked SNPs using BOLT-LMM^25,26^.

We simulated genotypes at 100,000 unlinked SNPs and case-control status plus family history (parental history for both parents) for 40,000 unrelated target samples. We simulated genotypes for both parents using the same minor allele frequency (MAF) values as the training data, used these to simulate genotypes for target samples (offspring), and simulated case-control status for both parents and target samples using a liability threshold model; target samples were not ascertained for case-control status. Again, for computational simplicity we generated 10 genotype matrices for the testing data and given these genotype matrices, then generated casecontrol and family history information under 16 different scenarios; the same 10 scenarios as the training data as well as 4 additional scenarios in which family members shared environmental correlation and 2 additional scenarios in which the parental disease prevalence was double that of the offspring (Supplementary Table 1). We note that environmental correlation and differing parental prevalence does not impact our training data as we are using case-control data only (not family history data) to train.

Within the target samples we use 10-fold cross-validation: for each fold we use the remaining 9 folds to estimate relevant model parameters. Given these parameters, the predicted risk of disease can be estimated for each individual within the held-out fold.

### UK Biobank data set

We analyzed 10 complex diseases from the UK Biobank^24^. To construct PRS, we computed prediction *β* for genotyped SNPs using all British individuals using BOLT-LMM^25,26^. These individuals were individuals of European ancestry (based on self-reported white-ethnicity) and British-ancestry individuals passing principal component analysis filters^24^. Our PRS consisted of 672,288 SNPs with missingness <10% and minor allele frequency (MAF) >0.1%; we mean normalized PRS based on the allele frequency within the training population.

We considered three distinct testing sets; these consisted of non-British European, South Asian (Indian, Pakistani, Bangladeshi), and African individuals (Black or Black British, Caribbean, African, Any other Black background). These testing sets were constructed through self-reported ethnicity; non-British European were individuals of European ancestry (based on self-reported white-ethnicity; White, British, Irish, Any other white background) who did not pass British-ancestry principal component analysis filters. We restricted to unrelated individuals (both unrelated to other individuals within the testing sets as well as unrelated to the training set). We use 10-fold cross-validation within the three testing sets to estimate relevant model parameters for both PRS-FH_log_ and PRS-FH_liab_. In detail, for individuals in a given fold we estimate the relevant parameters using the remaining 9-folds and use these parameters to predict risk.

UK Biobank collects family history of disease information for 12 diseases, in this work we focused on the 10 diseases for which PRS or FH produces a positive liability-scale *R*^2^ with a p-value less than the nominal 0.05/36 within non-British Europeans (Table 1; Supplementary Table 8). We primarily focused on three well-powered diseases (type 2 diabetes, depression, and hypertension) with (liability-scale) prediction *R*^2^ > 0.05 for PRS and/or FH in each target population (Supplementary Table 9). We note that depression was included as a well-powered disease despite its low SNP-heritability, because the contribution of sibling disease history (Supplementary Table 11) led to prediction *R*^2^ > 0.05 for FH in each target population. On the other hand, CAD was not included as a well-powered disease, due to poor performance of both PRS and FH in the African target population. For any individuals who reported 0 relevant siblings, disease status of siblings was set to 0.

### Application of PRS-FH_log_ to UK Biobank data

We applied PRS-FH_log_ to 10 complex diseases from the UK Biobank. Prior to model training and fitting, individuals with missing parental disease status for a given parental class (mother or father) were assigned the mean parental disease status for the respective parental class across the 9 training folds. Individuals with missing sibling disease status (for whom the number of siblings must be at least one or unknown) were assigned the mean sibling disease status across all individuals in the 9 training folds if the number of siblings was unknown, or the mean sibling disease status across individuals in the 9 training folds with at least one sibling if the number of siblings was known. Individuals with missing number of siblings were assigned the mean number of siblings across the 9 training folds if the sibling disease status was unknown, or the mean number of siblings subject to the same sibling disease status if known.

### Application of PRS-FH_liab_ to UK Biobank data

We applied PRS-FH_liab_ to 10 complex diseases from the UK Biobank. We used different estimates of disease prevalence for mother, father, siblings, and offspring; in any fold, when the prevalence of disease was 0 (for mother, father, sibling, or offspring) we set it equal to 1/(number of individuals within that fold). For all analyses that included sibling history, except when otherwise specified, the liability threshold for disease used to predict disease risk accounted for number of siblings (indicator variables for 0, 1, 2, 3, 4, ≥5 siblings), as we generally observed a U-shaped relationship between disease prevalence and number of siblings (Supplementary Table 16).

We used estimates of pseudo-heritability that differ for mother, father, and siblings; pseudo-heritability was estimated using maximum-likelihood (see Supplementary Note). If there were pairs of individuals with concordant disease status (e.g. both offspring and relative have disease), pseudo-heritability was set to 0 and the liability was not conditioned on this relative. UK Biobank collects sibling disease history as a binary “at least one” affected indicator, and as such we could estimate pseudo-heritability either using individuals with exactly one sibling or using all individuals with at least one sibling (see Supplementary Note). We elected to estimate pseudo-heritability using all individuals with at least one sibling, as the number of individuals with exactly one sibling can be prohibitively low (and may include no concordant disease pairs for diseases of low prevalence). In some of our early experiments in this project, we observed computational problems when estimated pseudo-heritability > 1.8 (i.e. pseudo-heritability/2 ≥ 0.9). Our software thus caps estimates of pseudo-heritability at 1.8. However, this did not impact any of the analyses reported in the current manuscript.

We estimated the amount of variance explained by the PRS on the liability scale, *V*, that varied based on the target population (Equation (4)). For each fold, we ran a permutation test (1000 permutations) of *H*_o_: *V* = 0, if we failed to reject the null hypothesis with *p* > 0.05 we set *V* = 0 (i.e. we did not use PRS to inform posterior disease risk).

### Decreasing the number of training samples from the target population

We decreased the number of training samples from the target population to different values of expected effective training sample size (*N_eff_*, which can vary with the number of *cases* sampled; see Equation (6)). For a given value of expected *N_eff_*, we constructed multiple independent training sample sets of that size by down-sampling individuals from each of the 10 folds, and averaged the resulting prediction accuracies across the training sample sets of that size (Supplementary Table 26 and Supplementary Figure 3).

The effective sample size (*N_eff_*) is computed for a training sample as:

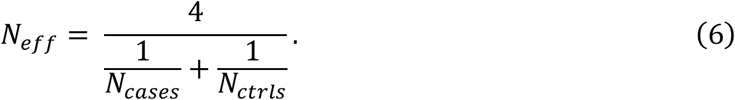

### Incorporation of covariates in UK Biobank analyses

We repeated the analyses of 10 complex diseases from the UK Biobank by incorporating covariates into each method. We included 23 covariates: age, sex, BMI, and 20 principal components. Prior to model training and fitting, individuals with missing BMI, age, or sex were assigned the mean age, sex, and BMI across the 9 training folds. The prediction method based solely on covariates modeled the 23 covariates linearly impacting the log-odds of disease for an individual; PRS+ (the prediction method based on covariates and PRS) models PRS and the 23 covariates (age, sex, BMI, and 20 principal components) linearly impacting the log-odds of disease for an individual. PRS-FH_log_ simply adds the 23 covariates into the logistic model incorporating family history of disease (FH_log_) or PRS and family history of disease (PRS-FH_log_). For FH_liab_ and PRS-FH_liab_, covariates were modeled as impacting the threshold for disease; a logistic model for disease as a function of covariates (23 covariates as well as indicators for number of siblings) was used to predict the risk of disease for an individual, thereby estimating the threshold for disease conditional on covariates.

### Jackknife standard errors of prediction accuracy and differences in prediction accuracy

We report estimates of liability-scale 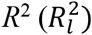 or the difference in 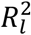 between two methods 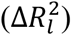. Given predicted disease risks (*r*) and observed phenotypes (*Z*), *R_o_* is estimated as 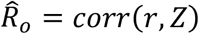 and 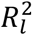 is estimated as 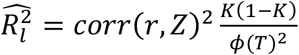 (ref. ^27^) for which *corr*(*r,Z*) = max (*corr*(*r,Z*), 0). (When using 10-fold cross-validation within a testing set to estimate relevant model parameters, 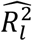 for a method is computed by concatenating across the 10 folds and computing a single 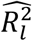, rather than computing the average of 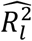 across the 10 folds.)

To test for significantly non-zero prediction accuracy or differences between methods we assess whether *R_o_* or Δ*R_o_* (where Δ denotes either the difference between 2 prediction methods, or the difference versus a covariates-only predictor in the setting with covariates) is significantly different from zero. We compute both jackknife standard errors as well as jackknife p-values (for *H_o_*: *R_o_* > 0 or *H_o_*:Δ*R_o_* = 0), employing a jackknife across individuals (we note that the alternative of employing a genomic block-jackknife is of interest for evaluation of PRS methods, but is not applicable to evaluation of FH and PRS-FH methods). We let all individuals in fold *i* be represented by *D_i_* and we construct *n* jackknife samples (*n*=100 in this study) by deleting each of the *n* folds as follows, *D*_[*i*]_ = {*D*_1_, *D*_2_,…, *D*_i–1_,…,*D*_i+1_,…,*D*_n_}. Each of these *D*_[*i*]_ are denoted blocks. We then compute 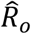 on each *D*_[*i*]_, denoting each such value as 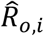. We then define the jackknife variance as

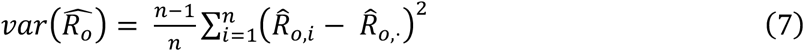

where 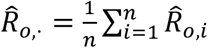. The jackknife variance for the difference in *R_o_* (or for 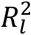) between two methods is computed in a similar manner. We computed a jackknife p-value by constructing pseudovalues as 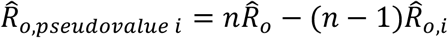 and test the hypothesis *H_o_*:*R_o_* = 0 by using the fact that

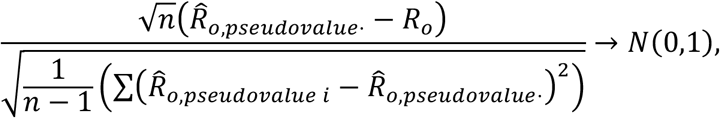

where 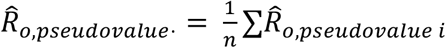. The jackknife p-value for *H_o_*: Δ*R_o_* 0 between two methods is computed in a similar manner. (We note the *n* folds used when computing jackknife standard errors and p-values are unrelated to the 10-folds used during cross-validation: individuals are concatenated across the 10 cross-validation-folds and then randomly assigned to the *n* jackknife folds.)

Jackknife assumes independence between blocks, while individuals are independent (by construction), individual predictions within a fold could use information from other folds, thus potentially inducing a correlation. To determine the potential effect of this we assessed the calibration of jackknife standard errors in simulations. For every simulation scenario, we computed an estimated variance of *R_o_* across the 10 simulation replicates (denoted empirical variance), as well as the average jackknife variance across the 10 simulation replicates (denoted mean jackknife variance). Across all simulation scenarios, the sum of the empirical variance was 0.00063 and 0.00059 while the sum of the mean jackknife variance was 0.00060 and 0.00057 for PRS-FH_log_ and PRS-FH_liab_, respectively. This suggests the standard errors are well-calibrated.

