## Supplementary for "Incorporating family history of disease improves polygenic risk scores in diverse populations"

**Supplementary Note**

**Pseudo-heritability: definition, estimation, and justification**

PRS-FH$_{liab}$ models the family history of disease and PRS using a liability threshold model in which the covariance between an individual's and a relative's liability is a function of the pseudo-heritability of the disease. In more detail, consider,

$$\begin{pmatrix}\epsilon_p\\ \epsilon_o\end{pmatrix} \sim N\left(\begin{pmatrix}0\\0\end{pmatrix},\begin{pmatrix}1 & h^2/2+\sigma\\ h^2/2+\sigma & 1\end{pmatrix}\right) = N\left(\begin{pmatrix}0\\0\end{pmatrix},\begin{pmatrix}1 & \tilde{h}^2/2\\ \tilde{h}^2/2 & 1\end{pmatrix}\right), \quad (1)$$

where $h^2$ denotes heritability, $\sigma$ some other shared effects, say environmental, and $\tilde{h}^2$ denotes pseudo-heritability.

We elected to use maximum-likelihood estimates of pseudo-heritability. In detail, considering equation (1) and using properties of truncated normals as well as selection theory:

$$E(\epsilon_p|truncation) = \lambda(T_p) = \begin{cases}\frac{\phi(T_p)}{1-\Phi(T_p)} & \epsilon_p \geq T_p\\ \frac{-\phi(T_p)}{\Phi(T_p)} & \epsilon_p < T_p\end{cases}; Var(\epsilon_p|truncation) = 1-\lambda(T_p)(\lambda(T_p)-T_p) \quad (2)$$

$$\epsilon_o|(\epsilon_p \geq T_p) \sim N(0.5\tilde{h}^2\lambda(T_p), 1-(0.5\tilde{h}^2)^2\lambda(T_p)(\lambda(T_p)-T_p)). \quad (3)$$

We know $D_p = 1$ if and only if $\epsilon_p \geq T_p$ and $D_o = 1$ if and only if $\epsilon_o \geq T_o$. We can consider the data as coming from a multinomial distribution with $p_{ij} = Pr(D_o = i \& D_p = j)$. Note that $p_{ij}$ is a function of $\tilde{h}^2$ (treated as unknown) and $K_o, K_p$ (treated as known).
Therefore the log-likelihood is:

$$\ell \propto \sum_{i,j\in\{0,1\}} N_{ij}\log p_{ij}, \quad (4)$$

where $N_{ij}$ is the number of individuals with $D_o = i \& D_p = j$. We can maximize the above with respect to the one unknown, $\tilde{h}^2$, to obtain the maximum likelihood estimate.
Consider the setting with siblings, where we know *at least one* sibling is affected and we know the number of siblings ($n_s$) per individual. In this case we have 2 options:
**1** We can restrict attention to individuals with one sibling and compute pseudo-heritability as was done in the case of parental history. This however has the disadvantage that for certain testing sets and diseases, the number of individuals with exactly one sibling is prohibitively low.
**2** We can note the following relevant multinomial probabilities:

$$p_{11,n_s} = Pr(D_o = 1 \& D_{sib} \geq 1|n_s); p_{10,n_s} = Pr(D_o = 1 \& D_{sib} = 0|n_s) \quad (5)$$

$$p_{01,n_s} = Pr(D_o = 0 \& D_{sib} \geq 1|n_s); p_{00,n_s} = Pr(D_o = 0 \& D_{sib} = 0|n_s). \quad (6)$$

For a given pseudo-heritability we can compute the distribution of $\vec{\epsilon}_s|z_o$ and then use `pmvnorm` from the package `mvtnorm` to compute the probability $\epsilon_s < T_s \forall s \in \{1, \ldots, n_s\}$. Then compute the following:

$$p_{11,n_s} = (1 - Pr(\epsilon_s < T_s \forall s \in \{1, \ldots, n_s\} | \epsilon_o \geq T_o))K_o; \tag{7}$$
$$p_{10,n_s} = Pr(\epsilon_s < T_s \forall s \in \{1, \ldots, n_s\} | \epsilon_o \geq T_o)K_o; \tag{8}$$
$$p_{01,n_s} = (1 - Pr(\epsilon_s < T_s \forall s \in \{1, \ldots, n_s\} | \epsilon_o \geq T_o))(1 - K_o); \tag{9}$$
$$p_{00,n_s} = Pr(\epsilon_s < T_s \forall s \in \{1, \ldots, n_s\} | \epsilon_o \geq T_o)(1 - K_o). \tag{10}$$

Therefore the log-likelihood is:

$$\ell \propto \sum_{n_s \in \{1, \ldots, N_s\}} \sum_{i,j \in \{0,1\}} N_{ij} \log p_{ij,n_s}, \tag{11}$$

where $N_s$ is the maximum number of siblings an individual has in the dataset. Note that again $p_{ij,n_s}$ is a function of $\tilde{h}^2$ (treated as unknown) and $K_o, K_s, n_s$ (treated as known). We can again maximize the log-likelihood to obtain the maximum likelihood estimate for $\tilde{h}^2$.

We examined the impact of using heritability versus pseudo-heritability in PRS-FH$_{liab}$ and how this was affected by environmental correlation in simulations (Supplementary Table 25). We compared the performance of PRS-FH$_{liab}$ (which uses a different covariance between the target individual's total liability and the relative's liability for each type of relative) to two modified PRS-FH$_{liab}$ methods, one uses the true generative heritability and the other uses the average parental covariance. When there is no environmental correlation, all prediction methods perform similarly; when there is environmental correlation that is consistent across family members both methods estimating pseudo-heritability perform similarly and better than the method using the true generative heritability; when there is environmental correlation that is different across family members PRS-FH$_{liab}$ (estimating pseudo-heritability and modeling mother and father history distinctly) achieves the highest predictive accuracy.

**Estimating the amount of variance explained by the PRS on the liability-scale**

PRS-FH$_{liab}$ models the family history of disease and PRS using a liability threshold model in which the covariance between an individual's liability and the measured component of the liability (the PRS, $M$) is the amount of variance explained by the PRS on the liability-scale, $V$. In more detail, consider,

$$\begin{pmatrix} \epsilon \\ M \end{pmatrix} \sim N\left(\begin{pmatrix} 0 \\ 0 \end{pmatrix}, \begin{pmatrix} V & V \\ V & 1 \end{pmatrix}\right), \tag{12}$$

where $M$ is a PRS on the liability scale. Therefore,

$$\rho = corr(PRS, D) = corr(M, D) = \frac{E[MD]}{\sqrt{K(1-K)}\sqrt{V}} = \frac{E[M|D=1]Pr(D=1)}{\sqrt{K(1-K)}\sqrt{V}} \tag{13}$$

Moreover,

$$E[M|D=1] = E[M|\epsilon \geq T] = 0 + V * (1)^{-1} * (E(\epsilon|\epsilon \geq T) - 0) = V\frac{\phi(T)}{1-\Phi(T)} \tag{14}$$

Therefore,

$$\rho = \frac{V\frac{\phi(T)}{1-\Phi(T)}K}{\sqrt{K(1-K)}\sqrt{V}} \rightarrow V = \rho^2 \frac{1-K}{K}\left(\frac{1-\Phi(T)}{\phi(T)}\right)^2 \tag{15}$$

Note that $1 - \Phi(T) = K$ therefore,

$$V = \rho^2 \frac{(1-K)K}{\phi(T)^2} \tag{16}$$

which is a transformation that has been previously proposed to go from observed to liability scale (Lee et al. 2012 Genetic Epi; Lee et al. 2011 AJHG).

**Posterior mean and variance of liability for individuals with at least one sibling affected**

In order to determine the posterior distribution for individuals with at least one sibling affected, we computed a weighted average. That is, denoting $\mathcal{P}$ as given parental disease history,

$$E(\epsilon_o|\mathcal{P}, D_s \geq 1, n_s) = \sum_{i=1}^{n_s} E(\epsilon_o|\mathcal{P}, D_s = i, n_s) Pr(D_s = i|\mathcal{P}, D_s \geq 1, n_s) \tag{17}$$

and,

$$\begin{aligned} Var(\epsilon_o|\mathcal{P}, D_s \geq 1, n_s) &= \sum_{i=1}^{n_s} Var(\epsilon_o|\mathcal{P}, D_s = i, n_s) Pr(D_s = i|\mathcal{P}, D_s \geq 1, n_s) \\ &+ \sum_{i=1}^{n_s} E(\epsilon_o|\mathcal{P}, D_s = i, n_s)^2 (1 - Pr(D_s = i|\mathcal{P}, D_s \geq 1, n_s)) Pr(D_s = i|\mathcal{P}, D_s \geq 1, n_s) \\ &- 2\sum_{i=2}^{n_s}\sum_{j=1}^{i-1} E(\epsilon_o|\mathcal{P}, D_s = i, n_s) E(\epsilon_o|\mathcal{P}, D_s = j, n_s) Pr(D_s = j|\mathcal{P}, D_s \geq 1, n_s) Pr(D_s = i|\mathcal{P}, D_s \geq 1, n_s) \end{aligned} \tag{18}$$

For an individual with a given $\mathcal{P}$ and number of siblings $n_s$ we compute the mean and variance of $\epsilon_o$ conditional on $\mathcal{P}$ as well as $i$ affected siblings $\forall i \in \{0, 1, \ldots, n_s\}$. Moreover, we compute the probability of observing the number of affected siblings ($i$) given $\mathcal{P}$ (through the function `pmvnorm()` in `R`). That is, we have,

$$E(\epsilon_o|\mathcal{P}, D_s = i, n_s); Var(\epsilon_o|\mathcal{P}, D_s = i, n_s); Pr(D_s = i|\mathcal{P}, n_s) \quad \forall i \in \{0, 1, \ldots, n_s\}. \tag{19}$$

The three above parameters allow us to compute $E(\epsilon_o|\mathcal{P}, D_s \geq 1, n_s)$ and $Var(\epsilon_o|\mathcal{P}, D_s \geq 1, n_s)$ by noting, for $i > 0$,

$$Pr(D_s = i|\mathcal{P}, D_s \geq 1, n_s) = \frac{Pr(D_s = i, D_s \geq 1|\mathcal{P}, n_s)}{Pr(D_s \geq 1|\mathcal{P}, n_s)} = \frac{Pr(D_s = i|\mathcal{P}, n_s)}{\sum_{j=1}^{n_s} Pr(D_s = j|\mathcal{P}, n_s)}, \tag{20}$$

While $\mathcal{P}$ denoted parental disease history this derivation can be extended to include both parental disease history and the PRS of the individual, thereby allowing for the computation of posterior distribution for individuals with at least one sibling affected both when incorporating only family history ($FH_{liab}$) and PRS and family history ($PRS - FH_{liab}$).

| | K | $h_l^2$ | $h_g^2$ | M | C | Correlation |
|---|---|---|---|---|---|---|
| 1 | 0.05 | 0.5 | 0.25 | 100K | 10K | 0 |
| 2 | 0.25 | 0.5 | 0.25 | 100K | 10K | 0 |
| 3 | 0.05 ($K_p = 0.10$) | 0.5 | 0.25 | 100K | 10K | 0 |
| 4 | 0.25 ($K_p = 0.50$) | 0.5 | 0.25 | 100K | 10K | 0 |
| 5 | 0.05 | 0.5 | 0.25 | 100K | 10K | Cov=$0.5(0.5h_l^2)$ |
| 6 | 0.25 | 0.5 | 0.25 | 100K | 10K | Cov=$0.5(0.5h_l^2)$ |
| 7 | 0.05 | 0.5 | 0.25 | 100K | 10K | $\text{Cov}_1 = 0.25(0.5h_l^2)$; $\text{Cov}_2 = 0.75(0.5h_l^2)$ |
| 8 | 0.25 | 0.5 | 0.25 | 100K | 10K | $\text{Cov}_1 = 0.25(0.5h_l^2)$; $\text{Cov}_2 = 0.75(0.5h_l^2)$ |
| 9 | 0.05 | 0.25 | 0.125 | 100K | 10K | 0 |
| 10 | 0.25 | 0.25 | 0.125 | 100K | 10K | 0 |
| 11 | 0.05 | 0.75 | 0.375 | 100K | 10K | 0 |
| 12 | 0.25 | 0.75 | 0.375 | 100K | 10K | 0 |
| 13 | 0.05 | 0.5 | 0.25 | 100K | 5K | 0 |
| 14 | 0.25 | 0.5 | 0.25 | 100K | 5K | 0 |
| 15 | 0.05 | 0.5 | 0.25 | 100K | 15K | 0 |
| 16 | 0.25 | 0.5 | 0.25 | 100K | 15K | 0 |

Supplementary Table 1: **Simulation parameter settings.** We report the simulation framework for data generation for all scenarios considered. M is the total number of SNPs given to BOLT-LMM; C is the total number of causal SNPs given to BOLT-LMM; For scenarios 3-6, these changes impact testing data only and not training data

| | PRS | | $\text{FH}_{log}$ | | $\text{FH}_{liab}$ | | $\text{PRS-FH}_{log}$ | | $\text{PRS-FH}_{liab}$ | |
|---|---|---|---|---|---|---|---|---|---|---|
| K | mean | s.e. | mean | s.e. | mean | s.e. | mean | s.e. | mean | s.e. |
| 1% | 0.0167 | 0.00105 | 0.0228 | 0.00294 | 0.023 | 0.00276 | 0.0386 | 0.00377 | 0.039 | 0.00331 |
| 5% | 0.0523 | 0.00115 | 0.0515 | 0.00342 | 0.0517 | 0.00335 | 0.103 | 0.00524 | 0.103 | 0.00496 |
| 25% | 0.116 | 0.000912 | 0.0753 | 0.00135 | 0.0753 | 0.00134 | 0.166 | 0.00103 | 0.166 | 0.00103 |

Supplementary Table 2: **Numerical results of main simulations.** For each prediction method we report the mean $R_l^2$ across the 10 simulation replicates as well as the standard error of this mean for different values of prevalence ($K$).

| | PRS | | $\text{FH}_{log}$ | | $\text{FH}_{liab}$ | | $\text{PRS-FH}_{log}$ | | $\text{PRS-FH}_{liab}$ | |
|---|---|---|---|---|---|---|---|---|---|---|
| Prevalence | mean | s.e. | mean | s.e. | mean | s.e. | mean | s.e. | mean | s.e. |
| 5% | 0.99 | 0.013 | 0.97 | 0.0058 | 0.98 | 0.0052 | 0.96 | 0.0058 | 0.99 | 0.0049 |
| 25% | 1 | 0.004 | 1 | 0.00082 | 1 | 0.0021 | 0.99 | 0.00096 | 0.99 | 0.0017 |

Supplementary Table 3: **Calibration results of main simulations.** We report the mean slope from the regression of disease status on predictions across the 10 simulation replicates as well as the standard error of this mean for each prediction method.

| K | | PRS | | FH$_{log}$ | | FH$_{liab}$ | | PRS-FH$_{log}$ | | PRS-FH$_{liab}$ | |
|---|---|---|---|---|---|---|---|---|---|---|---|
| $K_o$ | $K_p$ | mean | s.e. | mean | s.e. | mean | s.e. | mean | s.e. | | |
| **5%** | **5%** | 0.0523 | 0.00115 | 0.0515 | 0.00342 | 0.0517 | 0.00335 | 0.103 | 0.00524 | 0.103 | 0.00496 |
| 5% | 10% | 0.0523 | 0.00115 | 0.0698 | 0.00471 | 0.07 | 0.00464 | 0.119 | 0.0063 | 0.12 | 0.00606 |
| **25%** | **25%** | 0.116 | 0.000912 | 0.0753 | 0.00135 | 0.0753 | 0.00134 | 0.166 | 0.00103 | 0.166 | 0.00103 |
| 25% | 50% | 0.116 | 0.000912 | 0.0819 | 0.00121 | 0.0819 | 0.0012 | 0.171 | 0.00115 | 0.171 | 0.00113 |

Supplementary Table 4: **Results of simulations at different values of parental prevalence.** For each prediction method we report the mean $R_l^2$ across the 10 simulation replicates as well as the standard error of this mean. Default simulation parameters are denoted in bold font.

| | PRS | | FH$_{log}$ | | FH$_{liab}$ | | PRS-FH$_{log}$ | | PRS-FH$_{liab}$ | |
|---|---|---|---|---|---|---|---|---|---|---|
| Scenario | mean | s.e. | mean | s.e. | mean | s.e. | mean | s.e. | mean | s.e. |
| K=5% | | | | | | | | | | |
| **No correlation** | 0.0523 | 0.00115 | 0.0515 | 0.00342 | 0.0517 | 0.00335 | 0.103 | 0.00524 | 0.103 | 0.00496 |
| Correlation | 0.053 | 0.0014 | 0.167 | 0.00465 | 0.167 | 0.00454 | 0.217 | 0.00424 | 0.218 | 0.00408 |
| Differential correlation | 0.0546 | 0.00118 | 0.183 | 0.00531 | 0.183 | 0.00524 | 0.234 | 0.00539 | 0.236 | 0.00534 |
| K=25% | | | | | | | | | | |
| **No correlation** | 0.116 | 0.000912 | 0.0753 | 0.00135 | 0.0753 | 0.00134 | 0.166 | 0.00103 | 0.166 | 0.00103 |
| Correlation | 0.115 | 0.0015 | 0.187 | 0.00216 | 0.187 | 0.00216 | 0.263 | 0.00275 | 0.263 | 0.00272 |
| Differential correlation | 0.117 | 0.00113 | 0.194 | 0.00137 | 0.195 | 0.00137 | 0.273 | 0.00158 | 0.273 | 0.00159 |

Supplementary Table 5: **Results of simulations at different values of environmental correlation.** For each prediction method we report the mean $R_l^2$ across the 10 simulation replicates as well as the standard error of this mean. Default simulation parameters are denoted in bold font.

| | PRS | | FH$_{log}$ | | FH$_{liab}$ | | PRS-FH$_{log}$ | | PRS-FH$_{liab}$ | |
|---|---|---|---|---|---|---|---|---|---|---|
| Scenario | mean | s.e. | mean | s.e. | mean | s.e. | mean | s.e. | mean | s.e. |
| K=5% | | | | | | | | | | |
| $h_l^2 = 0.25, h_g^2 = 0.125$ | 0.015 | 0.00065 | 0.0097 | 0.00068 | 0.0097 | 0.00068 | 0.024 | 0.0012 | 0.024 | 0.0012 |
| $\boldsymbol{h_l^2 = 0.5, h_g^2 = 0.25}$ | 0.052 | 0.0011 | 0.051 | 0.0034 | 0.052 | 0.0034 | 0.1 | 0.0052 | 0.1 | 0.005 |
| $h_l^2 = 0.75, h_g^2 = 0.375$ | 0.12 | 0.0021 | 0.17 | 0.0052 | 0.17 | 0.0052 | 0.28 | 0.0073 | 0.29 | 0.0069 |
| K=25% | | | | | | | | | | |
| $h_l^2 = 0.25, h_g^2 = 0.125$ | 0.031 | 0.00047 | 0.018 | 0.00049 | 0.018 | 0.00049 | 0.046 | 0.00084 | 0.046 | 0.00083 |
| $\boldsymbol{h_l^2 = 0.5, h_g^2 = 0.25}$ | 0.12 | 0.00091 | 0.075 | 0.0014 | 0.075 | 0.0013 | 0.17 | 0.001 | 0.17 | 0.001 |
| $h_l^2 = 0.75, h_g^2 = 0.375$ | 0.24 | 0.0022 | 0.19 | 0.0013 | 0.19 | 0.0013 | 0.36 | 0.0022 | 0.36 | 0.0022 |

Supplementary Table 6: **Results of simulations at different values of heritability and SNP-heritability.** For each prediction method we report the mean $R_l^2$ across the 10 simulation replicates as well as the standard error of this mean. Default simulation parameters are denoted in bold font.

| | PRS | | $FH_{log}$ | | $FH_{liab}$ | | PRS-$FH_{log}$ | | PRS-$FH_{liab}$ | |
|---|---|---|---|---|---|---|---|---|---|---|
| Scenario | mean | s.e. | mean | s.e. | mean | s.e. | mean | s.e. | mean | s.e. |
| K=5% | | | | | | | | | | |
| C=5,000 | 0.078 | 0.00152 | 0.0523 | 0.00277 | 0.0525 | 0.00277 | 0.129 | 0.00339 | 0.131 | 0.00333 |
| **C=10,000** | 0.0523 | 0.00115 | 0.0515 | 0.00342 | 0.0517 | 0.00335 | 0.103 | 0.00524 | 0.103 | 0.00496 |
| C=15,000 | 0.0472 | 0.000926 | 0.0552 | 0.00151 | 0.0553 | 0.00153 | 0.0999 | 0.00132 | 0.101 | 0.00122 |
| K=25% | | | | | | | | | | |
| C=5,000 | 0.172 | 0.00172 | 0.0759 | 0.00137 | 0.0759 | 0.00137 | 0.216 | 0.00185 | 0.216 | 0.00187 |
| **C=10,000** | 0.116 | 0.000912 | 0.0753 | 0.00135 | 0.0753 | 0.00134 | 0.166 | 0.00103 | 0.166 | 0.00103 |
| C=15,000 | 0.0985 | 0.00134 | 0.0739 | 0.00145 | 0.0739 | 0.00145 | 0.152 | 0.00211 | 0.152 | 0.0021 |

Supplementary Table 7: **Results of simulations at different values of polygenicity.** For each prediction method we report the mean $R_l^2$ across the 10 simulation replicates as well as the standard error of this mean. Default simulation parameters are denoted in bold font.

| | PRS | | $FH_{log}$ | | $FH_{liab}$ | |
|---|---|---|---|---|---|---|
| Trait | $R_l^2$ | p | $R_l^2$ | p | $R_l^2$ | p |
| AD | 0.038 | 0.0044 | 0.0091 | 0.045 | 0.034 | 0.02 |
| PD | 0.00062 | 0.15 | 0.019 | 0.016 | 0.033 | 0.022 |
| LungCancer | 0.0035 | 0.0024 | 0.016 | 0.00033 | 0.017 | 1.6e-05 |
| BowelCancer | 0.0057 | 1.6e-06 | 0.0078 | 1.7e-05 | 0.0062 | 0.0001 |
| Stroke | 0.0006 | 0.074 | 0.019 | 2.7e-11 | 0.018 | 8.4e-11 |
| COPD | 0.018 | 4.6e-33 | 0.089 | 6.5e-47 | 0.098 | 2.7e-63 |
| ProstateCancer | 0.069 | 3.7e-55 | 0.04 | 7.4e-09 | 0.04 | 7.1e-09 |
| T2D | 0.067 | 4.8e-114 | 0.11 | 1.9e-70 | 0.12 | 5.4e-76 |
| BreastCancer | 0.032 | 1.7e-38 | 0.011 | 2.9e-12 | 0.01 | 3e-11 |
| Depression | 0.0086 | 1.7e-32 | 0.064 | 7e-83 | 0.061 | 4.8e-82 |
| CAD | 0.032 | 7.3e-95 | 0.063 | 5.3e-91 | 0.068 | 1.5e-116 |
| HTN | 0.099 | 0 | 0.065 | 0 | 0.071 | 0 |

Supplementary Table 8: **Results of applying PRS and FH methods to 12 diseases in UK Biobank non-British Europeans to identify diseases with statistically significant prediction $R^2$.** We present $R_l^2$ values and jackknife p-values for the test $H_0 : R_o > 0$ for predictors based on PRS-alone (PRS), and family history alone (FH; both logistic and liability threshold model) within non-British Europeans. In this work we focused on the 10 diseases for which PRS or FH produces a positive liability-scale R2 with a p-value less than the nominal $0.05/36 = 0.0013\bar{8}$ within non-British Europeans; all diseases were retained except Alzheimer's disease/dementia (AD) and Parkinson's disease (PD). AD, Alzheimer's disease/dementia; PD, Parkinson's disease; COPD, chronic bronchitis/emphysema; T2D, type 2 diabetes; CAD, coronary artery disease; HTN, hypertension.

| Trait | PRS $R_l^2$ (s.e.) | PRS p | $FH_{log}$ $R_l^2$ (s.e.) | $FH_{log}$ p | $FH_{liab}$ $R_l^2$ (s.e.) | $FH_{liab}$ p | PRS-$FH_{log}$ $R_l^2$ (s.e.) | PRS-$FH_{log}$ p | PRS-$FH_{log}$ $\Delta_{PRS}$ p | PRS-$FH_{log}$ $\Delta_{FH}$ p | PRS-$FH_{liab}$ $R_l^2$ | PRS-$FH_{liab}$ p | PRS-$FH_{liab}$ $\Delta_{PRS}$ p | PRS-$FH_{liab}$ $\Delta_{FH}$ p | $\Delta$log,liab p |
|---|---|---|---|---|---|---|---|---|---|---|---|---|---|---|---|
| **Non-British European** | | | | | | | | | | | | | | | |
| LungCancer | 0.0035 (0.003) | 0.0024 | 0.016 (0.009) | 0.00033 | 0.017 (0.008) | 1.6e-05 | 0.02 (0.01) | 0.00016 | 0.021 | 0.11 | 0.021 (0.009) | 6.8e-06 | 0.003 | 0.08 | 0.87 |
| BowelCancer | 0.0057 (0.002) | 1.6e-06 | 0.0078 (0.004) | 1.7e-05 | 0.0062 (0.003) | 0.0001 | 0.013 (0.005) | 6e-09 | 0.046 | 0.015 | 0.012 (0.004) | 4.9e-08 | 0.098 | 0.0098 | 0.17 |
| Stroke | 6e-04 (8e-04) | 0.074 | 0.019 (0.006) | 2.7e-11 | 0.018 (0.006) | 8.4e-11 | 0.019 (0.006) | 7e-12 | 1.4e-05 | 0.64 | 0.018 (0.006) | 7.5e-11 | 0.0001 | 0.29 | 0.67 |
| COPD | 0.018 (0.003) | 4.6e-33 | 0.089 (0.01) | 6.5e-47 | 0.098 (0.01) | 2.7e-63 | 0.1 (0.01) | 3e-58 | 8.8e-19 | 0.0002 | 0.11 (0.01) | 8.7e-79 | 5.5e-28 | 1.4e-06 | 0.087 |
| ProstateCancer | 0.069 (0.009) | 3.7e-55 | 0.04 (0.01) | 7.4e-09 | 0.04 (0.01) | 7.1e-09 | 0.1 (0.02) | 3.9e-24 | 0.029 | 1.2e-06 | 0.1 (0.02) | 6e-27 | 0.013 | 1.9e-07 | 0.5 |
| T2D | 0.067 (0.006) | 4.8e-114 | 0.11 (0.01) | 1.9e-70 | 0.12 (0.01) | 5.4e-76 | 0.17 (0.01) | 5.8e-114 | 1.6e-20 | 4.6e-11 | 0.18 (0.02) | 9.6e-116 | 8.3e-22 | 1.8e-17 | 0.037 |
| BreastCancer | 0.032 (0.005) | 1.7e-38 | 0.011 (0.003) | 2.9e-12 | 0.01 (0.003) | 3e-11 | 0.027 (0.005) | 1.6e-23 | 0.18 | 4.7e-05 | 0.03 (0.006) | 4e-26 | 0.62 | 8.4e-08 | 0.0041 |
| Depression | 0.0086 (0.001) | 1.7e-32 | 0.064 (0.007) | 7e-83 | 0.061 (0.006) | 4.8e-82 | 0.071 (0.007) | 2.3e-100 | 2e-37 | 1.2e-05 | 0.067 (0.006) | 1.4e-93 | 1.4e-30 | 7.3e-11 | 0.12 |
| CAD | 0.032 (0.003) | 7.3e-95 | 0.063 (0.006) | 5.3e-91 | 0.068 (0.006) | 1.5e-116 | 0.092 (0.008) | 5.5e-129 | 4.8e-31 | 1.4e-18 | 0.097 (0.007) | 3.4e-155 | 1.6e-37 | 8.7e-25 | 0.004 |
| HTN | 0.099 (0.004) | 0 | 0.065 (0.003) | 0 | 0.071 (0.004) | 0 | 0.15 (0.004) | 0 | 5.7e-69 | 6.1e-128 | 0.15 (0.004) | 0 | 6.4e-30 | 6.3e-199 | 0.14 |
| Average | 0.058 | | 0.08 | | 0.084 | | 0.13 | | | | 0.13 | | | | |
| **South Asian** | | | | | | | | | | | | | | | |
| LungCancer | 8.6e-05 (0.002) | 0.94 | 0 (0) | | 0 (0) | | 0 (0) | | 0.12 | | 0 (0) | | 0.12 | | |
| BowelCancer | 0.0026 (0.005) | 0.17 | 0.0042 (0.006) | 0.18 | 0.019 (0.02) | 0.091 | 0.0049 (0.007) | 0.17 | 0.84 | 0.71 | 0.019 (0.02) | 0.091 | 0.39 | | 0.087 |
| Stroke | 0.00092 (0.002) | 0.18 | 0.014 (0.01) | 0.0021 | 0.013 (0.008) | 0.00065 | 0.013 (0.009) | 0.0028 | 0.079 | 0.45 | 0.013 (0.008) | 0.00065 | 0.075 | | 1 |
| COPD | 0.00097 (0.002) | 0.16 | 0.018 (0.02) | 0.014 | 0.015 (0.01) | 0.015 | 0.018 (0.01) | 0.014 | 0.12 | 0.7 | 0.015 (0.01) | 0.015 | 0.17 | | 0.72 |
| ProstateCancer | 0.042 (0.02) | 6.5e-05 | 0.0047 (0.007) | 0.14 | 0.023 (0.03) | 0.073 | 0.016 (0.01) | 0.013 | 0.15 | 0.18 | 0.035 (0.03) | 0.016 | 0.83 | 0.44 | 0.098 |
| T2D | 0.052 (0.009) | 5.9e-33 | 0.11 (0.01) | 3.8e-69 | 0.11 (0.01) | 2.3e-61 | 0.15 (0.02) | 2.7e-73 | 2e-19 | 1.1e-05 | 0.14 (0.02) | 1.6e-69 | 8.4e-13 | 2e-08 | 0.25 |
| BreastCancer | 0.017 (0.01) | 0.00055 | 0.011 (0.009) | 0.011 | 0.0094 (0.008) | 0.0095 | 0.012 (0.01) | 0.044 | 0.51 | 0.86 | 0.015 (0.01) | 0.0054 | 0.74 | 0.51 | 0.46 |
| Depression | 0.0048 (0.003) | 0.0011 | 0.071 (0.02) | 2.6e-11 | 0.069 (0.02) | 8.2e-12 | 0.077 (0.02) | 2.2e-11 | 3.8e-07 | 0.075 | 0.072 (0.02) | 6.6e-12 | 2.3e-06 | 0.04 | 0.65 |
| CAD | 0.018 (0.005) | 9.7e-14 | 0.082 (0.01) | 1.6e-30 | 0.073 (0.01) | 2.6e-30 | 0.096 (0.02) | 3.5e-34 | 2.3e-14 | 0.0019 | 0.087 (0.01) | 6e-33 | 1.7e-12 | 0.00011 | 0.056 |
| HTN | 0.063 (0.008) | 1.2e-56 | 0.074 (0.009) | 1e-56 | 0.069 (0.009) | 2e-48 | 0.12 (0.01) | 4.9e-97 | 1.9e-16 | 3.4e-14 | 0.11 (0.01) | 9.4e-80 | 2.3e-06 | 5.9e-24 | 0.0012 |
| Average | 0.04 | | 0.086 | | 0.083 | | 0.12 | | | | 0.11 | | | | |
| **African** | | | | | | | | | | | | | | | |
| LungCancer | 0 (0) | | 0.012 (0.01) | 0.2 | 0.062 (0.08) | 0.13 | 0.012 (0.02) | 0.23 | 0.46 | 0.34 | 0.062 (0.08) | 0.13 | 0.26 | | 0.22 |
| BowelCancer | 0.003 (0.005) | 0.13 | 0.0022 (0.004) | 0.22 | 0.0013 (0.003) | 0.25 | 0.0042 (0.007) | 0.19 | 0.89 | 0.38 | 0.0013 (0.003) | 0.25 | 0.8 | | 0.41 |
| Stroke | 0.0028 (0.003) | 0.05 | 0.0032 (0.004) | 0.047 | 0.0091 (0.008) | 0.01 | 0.0042 (0.004) | 0.027 | 0.75 | 0.53 | 0.0085 (0.007) | 0.012 | 0.46 | 0.56 | 0.26 |
| COPD | 0 (0) | | 0.0017 (0.003) | 0.17 | 0.0044 (0.006) | 0.11 | 0.0016 (0.003) | 0.18 | 0.37 | 0.94 | 0.0044 (0.006) | 0.11 | 0.23 | | 0.36 |
| ProstateCancer | 0.011 (0.007) | 0.00097 | 0.11 (0.05) | 3.4e-05 | 0.12 (0.05) | 7.3e-06 | 0.11 (0.05) | 1.6e-05 | 0.0065 | 0.75 | 0.12 (0.05) | 2.6e-06 | 0.0023 | 0.38 | 0.19 |
| T2D | 0.0052 (0.003) | 0.00021 | 0.13 (0.02) | 1.6e-33 | 0.12 (0.02) | 3.8e-33 | 0.13 (0.02) | 2.1e-35 | 2.4e-17 | 0.21 | 0.13 (0.02) | 6.5e-35 | 2.3e-15 | 0.025 | 0.21 |
| BreastCancer | 0.0016 (0.003) | 0.17 | 0.00017 (7e-04) | 0.38 | 0.0014 (0.003) | 0.22 | 1.1e-05 (2e-04) | 0.99 | 0.011 | 0.0067 | 0.0014 (0.003) | 0.22 | 0.96 | | 0.021 |
| Depression | 1.8e-05 (3e-04) | 0.51 | 0.06 (0.02) | 1.6e-08 | 0.063 (0.02) | 4.4e-09 | 0.059 (0.02) | 1.9e-08 | 1.1e-06 | 0.0026 | 0.063 (0.02) | 4.4e-09 | 3.5e-07 | | 0.61 |
| CAD | 0.0033 (0.003) | 0.0056 | 0.024 (0.009) | 1.3e-08 | 0.017 (0.008) | 9.2e-06 | 0.029 (0.01) | 3.2e-09 | 0.00011 | 0.063 | 0.019 (0.008) | 3.2e-06 | 0.018 | 0.069 | 0.088 |
| HTN | 0.011 (0.003) | 3.3e-10 | 0.1 (0.009) | 1.2e-114 | 0.1 (0.009) | 8e-118 | 0.11 (0.009) | 2.1e-126 | 2e-33 | 0.01 | 0.11 (0.009) | 4.3e-126 | 4.3e-28 | 0.00013 | 0.61 |
| Average | 0.0053 | | 0.096 | | 0.096 | | 0.1 | | | | 0.099 | | | | |

Supplementary Table 9: **Numerical results for 10 diseases from the UK Biobank.** For each disease, ancestry, and model the $R_l^2$ value (and jackknife standard error) is shown along with jackknife p-values for the test $H_0 : R_o > 0$. The jackknife p-value is shown for the difference between the prediction $R_l^2$ of PRS-FH with PRS and the respective FH predictor (e.g. PRS-$FH_{liab}^+$ vs $FH_{liab}^+$ and similarly for log; $H_0 : \Delta R_o = 0$). The average is shown across the three well-powered traits (grey shading).

| Trait | Non-British European | | South Asian | | African | |
|---|---|---|---|---|---|---|
| | $FH_{log}$ | $FH_{liab}$ | $FH_{log}$ | $FH_{liab}$ | $FH_{log}$ | $FH_{liab}$ |
| LungCancer | 0.03 | 0.03 | -0.01 | | -0.00 | |
| BowelCancer | 0.02 | 0.01 | 0.01 | | -0.00 | |
| Stroke | 0.02 | 0.03 | 0.02 | | 0.00 | 0.00 |
| COPD | 0.06 | 0.06 | 0.02 | | -0.02 | |
| ProstateCancer | 0.04 | 0.04 | 0.03 | 0.03 | -0.00 | -0.01 |
| T2D | 0.10 | 0.10 | 0.13 | 0.12 | 0.01 | 0.00 |
| BreastCancer | 0.06 | 0.06 | 0.03 | 0.03 | 0.01 | |
| Depression | 0.04 | 0.03 | 0.02 | 0.03 | 0.01 | |
| CAD | 0.09 | 0.09 | 0.07 | 0.06 | -0.05 | -0.02 |
| HTN | 0.14 | 0.13 | 0.12 | 0.12 | 0.04 | 0.04 |
| Average (all) | 0.06 | 0.06 | 0.04 | 0.07 | 0 | 0 |
| Average (3 well-powered) | 0.09 | 0.09 | 0.09 | 0.09 | 0.02 | 0.02 |

Supplementary Table 10: **Correlations between PRS vs. FH predictions for 10 diseases from the UK Biobank.** The average across all ancestries and all 10 diseases is 0.034 for $FH_{log}$ and 0.047 for $FH_{liab}$; across 3 well-powered it is 0.068 for $FH_{log}$ and 0.072 for $FH_{liab}$.

**(a)**

| Trait | Intercept | PRS | P1 | P2 | Sibs | NumSibs |
|---|---|---|---|---|---|---|
| | | | Non-British European | | | |
| LungCancer | -5.3 | 2.2e+02 | 0.53 | 0.52 | 1.3 | 0.033 |
| BowelCancer | -4.7 | 69 | 0.65 | 0.41 | 0.59 | 0.046 |
| Stroke | -4.2 | 17 | 0.17 | 0.36 | 0.96 | 0.063 |
| COPD | -3.8 | 29 | 0.84 | 0.75 | 1.1 | 0.078 |
| ProstateCancer | -3.6 | 27 | 0.54 | | 1.7 | -0.015 |
| T2D | -3.6 | 21 | 0.56 | 0.74 | 1.2 | -0.015 |
| BreastCancer | -2.9 | 13 | 0.5 | | 0.55 | -0.022 |
| Depression | -2.8 | 13 | 0.83 | 0.85 | 0.72 | 0.0078 |
| CAD | -2.9 | 12 | 0.33 | 0.45 | 0.89 | 0.019 |
| HTN | -1.2 | 4.7 | 0.18 | 0.24 | 0.81 | -0.0062 |
| | | | South Asian | | | |
| LungCancer | -6.7 | 66 | 0.76 | 1.6 | 1.5 | 0.061 |
| BowelCancer | -5.1 | 52 | 1.1 | 1.2 | 1.6 | -0.077 |
| Stroke | -4.2 | 21 | 0.18 | 0.34 | 0.82 | 0.079 |
| COPD | -4.1 | 8 | 0.35 | 0.96 | 0.85 | 0.05 |
| ProstateCancer | -4.6 | 25 | 1.2 | | 1.6 | -0.067 |
| T2D | -2.8 | 15 | 0.34 | 0.3 | 1 | 0.031 |
| BreastCancer | -3.2 | 11 | 0.66 | | 0.96 | -0.11 |
| Depression | -3.3 | 11 | 0.7 | 0.96 | 1.1 | 0.05 |
| CAD | -2.6 | 8.8 | 0.37 | 0.45 | 0.77 | 0.077 |
| HTN | -1.1 | 3.6 | 0.11 | -0.015 | 0.86 | 0.044 |
| | | | African | | | |
| LungCancer | -4.8 | -1.2e+02 | 0.55 | -2.6 | 2.8 | -0.16 |
| BowelCancer | -4.9 | 32 | -0.55 | 0.37 | 1.6 | -0.034 |
| Stroke | -4.1 | 20 | -0.041 | 0.3 | 0.96 | 0.017 |
| COPD | -3.9 | -5.7 | 0.79 | 0.58 | 0.98 | -0.054 |
| ProstateCancer | -3.7 | 3.7 | 0.78 | | 2.2 | 0.02 |
| T2D | -3.2 | 3.3 | 0.52 | 0.59 | 1.1 | 0.0085 |
| BreastCancer | -3.6 | 0.57 | 0.59 | | 0.82 | -0.034 |
| Depression | -3.2 | 0.23 | 0.33 | 1 | 1.2 | -0.017 |
| CAD | -3.4 | 4 | 0.52 | 0.61 | 0.43 | 0.055 |
| HTN | -1 | 1.1 | 0.3 | 0.26 | 1 | -0.001 |

**(b)**

| | | | | | | Thresholding by number of siblings ($n_s$) | | | | |
|---|---|---|---|---|---|---|---|---|---|---|
| Trait | V | $\tilde{h}^2_{p1}$ | $\tilde{h}^2_{p2}$ | $\tilde{h}^2_{sib}$ | Int. | $n_s = 1$ | $n_s = 2$ | $n_s = 3$ | $n_s = 4$ | $n_s \geq 5$ |
| | | | | | Non-British European | | | | | |
| LungCancer | 0.0036 | 0.24 | 0.21 | 0.28 | -4.9 | -0.34 | -0.48 | -0.15 | -0.5 | 0.094 |
| BowelCancer | 0.0057 | 0.27 | 0.17 | 0.14 | -4.6 | 0.095 | 0.011 | 0.23 | 0.26 | 0.35 |
| Stroke | 9e-05 | 0.096 | 0.2 | 0.49 | -3.9 | -0.13 | 0.032 | -0.05 | 0.094 | 0.4 |
| COPD | 0.018 | 0.52 | 0.44 | 0.66 | -3.1 | -0.56 | -0.51 | -0.4 | -0.1 | 0.34 |
| ProstateCancer | 0.069 | 0.32 | 0.32 | 0.72 | -3.4 | -0.22 | 0.013 | 0.032 | -0.1 | 0.11 |
| T2D | 0.067 | 0.41 | 0.54 | 0.87 | -2.9 | -0.39 | -0.31 | -0.25 | -0.21 | -0.068 |
| BreastCancer | 0.032 | 0.28 | 0.28 | 0.38 | -2.7 | -0.12 | -0.042 | -0.0023 | -0.099 | -0.17 |
| Depression | 0.0086 | 0.53 | 0.56 | 0.81 | -2.4 | -0.14 | -0.12 | -0.15 | -0.17 | -0.022 |
| CAD | 0.032 | 0.31 | 0.37 | 0.6 | -2.3 | -0.3 | -0.34 | -0.24 | -0.17 | 0.18 |
| HTN | 0.099 | 0.27 | 0.33 | 0.92 | -0.69 | -0.28 | -0.25 | -0.29 | -0.21 | -0.018 |
| | | | | | South Asian | | | | | |
| LungCancer | 0 | | | | -22 | 14 | 8.1e-10 | 16 | 16 | 15 |
| BowelCancer | 0 | 0.41 | 0.42 | 0.49 | -20 | 15 | 15 | 14 | 14 | 14 |
| Stroke | 0 | 0.12 | 0.21 | 0.29 | -4.1 | 0.55 | -0.12 | 0.4 | 0.21 | 0.69 |
| COPD | 0 | 0.23 | 0.47 | 1.3 | -3.2 | -1.1 | -0.59 | -0.87 | -0.8 | -0.5 |
| ProstateCancer | 0.043 | 0.61 | 0.61 | 0.81 | -4 | -0.26 | 0.11 | -0.41 | -0.98 | -0.23 |
| T2D | 0.052 | 0.4 | 0.41 | 0.98 | -1.4 | -0.46 | -0.56 | -0.46 | -0.32 | -0.091 |
| BreastCancer | 0.018 | 0.32 | 0.32 | 0.42 | -5.7 | 2.8 | 2.1 | 2.4 | 2.5 | 2.3 |
| Depression | 0.0049 | 0.58 | 0.67 | 1.3 | -2.7 | -0.35 | -0.82 | -0.11 | -0.28 | -0.038 |
| CAD | 0.018 | 0.35 | 0.43 | 0.57 | -1.7 | -0.64 | -0.25 | -0.34 | -0.12 | 0.13 |
| HTN | 0.063 | 0.24 | 0.16 | 0.93 | -0.39 | -0.52 | -0.37 | -0.17 | -0.13 | 0.037 |
| | | | | | African | | | | | |
| LungCancer | 0 | | | 0.88 | -8 | 2.4 | 3.1 | 2.7 | 1.9 | 2 |
| BowelCancer | 0 | | 0.16 | 0.5 | -5.2 | 0.9 | 0.36 | 0.29 | 0.88 | 0.45 |
| Stroke | 0.0008 | 0.023 | 0.16 | 0.56 | -3.5 | -0.49 | -0.26 | -0.079 | -0.63 | -0.088 |
| COPD | 0 | 0.34 | 0.28 | 0.39 | -3.9 | -0.39 | -0.58 | -0.12 | -0.61 | -0.53 |
| ProstateCancer | 0.011 | 0.54 | 0.54 | 0.93 | -2.8 | -0.22 | -0.34 | -0.51 | -0.27 | 0.021 |
| T2D | 0.0053 | 0.46 | 0.51 | 1.1 | -2 | -0.25 | -0.3 | -0.46 | -0.3 | -0.099 |
| BreastCancer | 0 | 0.26 | 0.26 | 0.43 | -3.5 | -0.44 | 0.33 | -0.34 | -0.014 | 0.0036 |
| Depression | 0 | 0.41 | 0.67 | 1.4 | -2.7 | -0.63 | -0.26 | -0.36 | -0.4 | -0.39 |
| CAD | 0.0034 | 0.31 | 0.36 | 0.52 | -2.5 | -0.68 | -0.48 | -0.48 | -0.27 | -0.12 |
| HTN | 0.011 | 0.37 | 0.38 | 1.1 | -0.22 | -0.19 | -0.29 | -0.2 | -0.088 | -0.014 |

**(c)**

| Trait | Intercept | PRS | P1 | P2 | Sibs | NumSibs |
|---|---|---|---|---|---|---|
| | | | Non-British European | | | |
| LungCancer | -5.3 | 2.2e+02 | 0.54 | 0.53 | 1.3 | 0.033 |
| BowelCancer | -4.7 | 69 | 0.65 | 0.41 | 0.59 | 0.047 |
| Stroke | -4.2 | 17 | 0.17 | 0.36 | 0.96 | 0.064 |
| COPD | -3.8 | 29 | 0.84 | 0.75 | 1.1 | 0.078 |
| ProstateCancer | -3.6 | 27 | 0.54 | | 1.7 | -0.015 |
| T2D | -3.6 | 21 | 0.57 | 0.74 | 1.2 | -0.015 |
| BreastCancer | -2.9 | 13 | 0.5 | | 0.55 | -0.022 |
| Depression | -2.8 | 13 | 0.83 | 0.85 | 0.72 | 0.0079 |
| CAD | -2.9 | 12 | 0.33 | 0.45 | 0.89 | 0.019 |
| HTN | -1.2 | 4.7 | 0.18 | 0.24 | 0.81 | -0.0062 |
| | | | South Asian | | | |
| LungCancer | -6.7 | 67 | 0.97 | 1.9 | 1.8 | 0.062 |
| BowelCancer | -5.1 | 52 | 1.2 | 1.2 | 1.7 | -0.077 |
| Stroke | -4.2 | 21 | 0.18 | 0.34 | 0.82 | 0.08 |
| COPD | -4.2 | 8.1 | 0.35 | 0.96 | 0.86 | 0.05 |
| ProstateCancer | -4.6 | 25 | 1.3 | | 1.7 | -0.067 |
| T2D | -2.8 | 15 | 0.34 | 0.3 | 1 | 0.031 |
| BreastCancer | -3.2 | 10 | 0.66 | | 0.97 | -0.11 |
| Depression | -3.3 | 11 | 0.7 | 0.96 | 1.1 | 0.05 |
| CAD | -2.6 | 8.8 | 0.37 | 0.45 | 0.77 | 0.077 |
| HTN | -1.1 | 3.6 | 0.11 | -0.015 | 0.86 | 0.044 |
| | | | African | | | |
| LungCancer | -4.8 | -1.2e+02 | 0.71 | -2.6 | 2.8 | -0.16 |
| BowelCancer | -4.9 | 32 | -0.3 | 0.4 | 1.6 | -0.034 |
| Stroke | -4.1 | 20 | -0.037 | 0.3 | 0.97 | 0.017 |
| COPD | -3.9 | -5.7 | 0.8 | 0.59 | 1 | -0.054 |
| ProstateCancer | -3.7 | 3.7 | 0.78 | | 2.2 | 0.02 |
| T2D | -3.2 | 3.3 | 0.52 | 0.59 | 1.1 | 0.0085 |
| BreastCancer | -3.6 | 0.58 | 0.6 | | 0.83 | -0.034 |
| Depression | -3.2 | 0.23 | 0.34 | 1 | 1.2 | -0.017 |
| CAD | -3.4 | 4 | 0.52 | 0.61 | 0.43 | 0.055 |
| HTN | -1 | 1.1 | 0.3 | 0.26 | 1 | -0.001 |

**(d)**

| Trait | V | $\tilde{h}^2_{p1}$ | $\tilde{h}^2_{p2}$ | $\tilde{h}^2_{sib}$ | Int. | Thresholding by number of siblings ($n_s$) $n_s = 1$ | $n_s = 2$ | $n_s = 3$ | $n_s = 4$ | $n_s \geq 5$ |
|---|---|---|---|---|---|---|---|---|---|---|
| **Non-British European** | | | | | | | | | | |
| LungCancer | 0.0035 | 0.24 | 0.21 | 0.26 | -4.9 | -0.34 | -0.48 | -0.15 | -0.49 | 0.093 |
| BowelCancer | 0.0057 | 0.27 | 0.17 | 0.14 | -4.6 | 0.095 | 0.01 | 0.23 | 0.26 | 0.35 |
| Stroke | 0 | 0.096 | 0.2 | 0.47 | -3.9 | -0.13 | 0.032 | -0.05 | 0.094 | 0.4 |
| COPD | 0.018 | 0.52 | 0.44 | 0.7 | -3.1 | -0.56 | -0.51 | -0.4 | -0.1 | 0.34 |
| ProstateCancer | 0.069 | 0.32 | 0.32 | 0.76 | -3.4 | -0.22 | 0.013 | 0.032 | -0.1 | 0.11 |
| T2D | 0.067 | 0.41 | 0.54 | 0.87 | -2.9 | -0.39 | -0.31 | -0.25 | -0.21 | -0.068 |
| BreastCancer | 0.032 | 0.28 | 0.28 | 0.37 | -2.7 | -0.12 | -0.042 | -0.0022 | -0.099 | -0.17 |
| Depression | 0.0086 | 0.53 | 0.56 | 0.82 | -2.4 | -0.14 | -0.12 | -0.15 | -0.17 | -0.022 |
| CAD | 0.032 | 0.31 | 0.37 | 0.6 | -2.3 | -0.3 | -0.34 | -0.24 | -0.17 | 0.18 |
| HTN | 0.099 | 0.27 | 0.33 | 0.92 | -0.69 | -0.28 | -0.25 | -0.29 | -0.21 | -0.018 |
| **South Asian** | | | | | | | | | | |
| LungCancer | 0 | 0.38 | 0.68 | 0.38 | -22 | 15 | -1e-09 | 16 | 16 | 15 |
| BowelCancer | 0 | 0.42 | 0.43 | 0.54 | -20 | 15 | 15 | 14 | 14 | 14 |
| Stroke | 0 | 0.12 | 0.21 | 0.27 | -4.1 | 0.53 | -0.13 | 0.38 | 0.19 | 0.67 |
| COPD | 0 | 0.23 | 0.47 | 1.3 | -3.2 | -1.1 | -0.59 | -0.87 | -0.8 | -0.5 |
| ProstateCancer | 0.042 | 0.62 | 0.62 | 0.78 | -4 | -0.29 | 0.074 | -0.45 | -1 | -0.27 |
| T2D | 0.052 | 0.4 | 0.41 | 0.96 | -1.4 | -0.46 | -0.56 | -0.46 | -0.31 | -0.091 |
| BreastCancer | 0.017 | 0.32 | 0.32 | 0.47 | -4.4 | 1.5 | 0.92 | 1.2 | 1.3 | 1.1 |
| Depression | 0.0048 | 0.59 | 0.67 | 1.3 | -2.7 | -0.35 | -0.82 | -0.11 | -0.28 | -0.04 |
| CAD | 0.018 | 0.35 | 0.43 | 0.57 | -1.7 | -0.64 | -0.26 | -0.34 | -0.12 | 0.13 |
| HTN | 0.063 | 0.24 | 0.16 | 0.93 | -0.39 | -0.52 | -0.37 | -0.17 | -0.14 | 0.037 |
| **African** | | | | | | | | | | |
| LungCancer | 0 | 0.34 | | 0.76 | -6.6 | 1 | 1.7 | 1.3 | 0.54 | 0.65 |
| BowelCancer | 0 | 3.6e-05 | 0.16 | 0.47 | -5.2 | 0.88 | 0.35 | 0.29 | 0.86 | 0.43 |
| Stroke | 0 | 0.01 | 0.16 | 0.58 | -3.5 | -0.49 | -0.26 | -0.077 | -0.62 | -0.089 |
| COPD | 0 | 0.34 | 0.28 | 0.32 | -3.9 | -0.39 | -0.57 | -0.12 | -0.61 | -0.53 |
| ProstateCancer | 0.011 | 0.54 | 0.54 | 0.93 | -2.8 | -0.22 | -0.33 | -0.51 | -0.27 | 0.018 |
| T2D | 0.0052 | 0.46 | 0.51 | 1.1 | -2 | -0.25 | -0.3 | -0.46 | -0.3 | -0.1 |
| BreastCancer | 0 | 0.27 | 0.27 | 0.47 | -3.5 | -0.44 | 0.32 | -0.34 | -0.017 | -0.0016 |
| Depression | 0 | 0.42 | 0.67 | 1.4 | -2.7 | -0.63 | -0.26 | -0.36 | -0.4 | -0.39 |
| CAD | 0.0033 | 0.31 | 0.36 | 0.52 | -2.5 | -0.68 | -0.48 | -0.48 | -0.27 | -0.12 |
| HTN | 0.011 | 0.37 | 0.38 | 1.1 | -0.22 | -0.19 | -0.29 | -0.2 | -0.088 | -0.014 |

Supplementary Table 11: **Model parameters estimated by PRS-FH$_{log}$ and PRS-FH$_{liab}$.** We report (a) model parameters estimated by PRS-FH$_{log}$ (using 10-fold cross-validation training data; average across 10 folds), (b) model parameters estimated by PRS-FH$_{liab}$ (using 10-fold cross-validation training data; average across 10 folds), (c) model parameters estimated by PRS-FH$_{log}$ (using all training data), (d) model parameters estimated by PRS-FH$_{liab}$ (using all training data).

| Trait | PRS | $FH_{log}$ | $FH_{liab}$ | PRS-$FH_{log}$ | PRS-$FH_{liab}$ |
|---|---|---|---|---|---|
| | | | Non-British European | | |
| LungCancer | 1.5 | 0.73 | 0.86 | 0.76 | 0.88 |
| BowelCancer | 0.79 | 0.74 | 0.69 | 0.82 | 0.79 |
| Stroke | 0.39 | 0.84 | 0.71 | 0.84 | 0.71 |
| COPD | 1.2 | 0.94 | 0.89 | 0.94 | 0.89 |
| ProstateCancer | 0.96 | 0.89 | 0.84 | 0.84 | 0.81 |
| T2D | 0.9 | 0.94 | 0.79 | 0.92 | 0.83 |
| BreastCancer | 0.92 | 0.9 | 0.74 | 0.72 | 0.69 |
| Depression | 1 | 0.94 | 0.61 | 0.94 | 0.63 |
| CAD | 0.93 | 0.96 | 0.87 | 0.97 | 0.91 |
| HTN | 0.97 | 0.99 | 0.66 | 1 | 0.8 |
| Average | 0.96 | 0.95 | 0.68 | 0.95 | 0.75 |
| | | | South Asian | | |
| LungCancer | 0.085 | -0.24 | -0.13 | -0.25 | -0.13 |
| BowelCancer | 0.28 | 0.26 | 0.5 | 0.27 | 0.5 |
| Stroke | 0.59 | 0.7 | 0.64 | 0.65 | 0.64 |
| COPD | 0.19 | 0.73 | 0.19 | 0.71 | 0.19 |
| ProstateCancer | 0.37 | 0.22 | 0.42 | 0.33 | 0.44 |
| T2D | 2.3 | 0.95 | 0.59 | 0.95 | 0.63 |
| BreastCancer | 0.46 | 0.67 | 0.55 | 0.44 | 0.49 |
| Depression | 0.62 | 0.83 | 0.37 | 0.85 | 0.37 |
| CAD | 1.2 | 0.95 | 0.82 | 0.96 | 0.85 |
| HTN | 0.9 | 0.98 | 0.57 | 0.99 | 0.66 |
| Average | 1.3 | 0.92 | 0.51 | 0.93 | 0.55 |
| | | | African | | |
| LungCancer | -0.39 | 0.29 | 0.56 | 0.28 | 0.56 |
| BowelCancer | 0.29 | 0.3 | 0.18 | 0.38 | 0.18 |
| Stroke | 0.5 | 0.42 | 0.45 | 0.44 | 0.43 |
| COPD | -0.072 | 0.26 | 0.34 | 0.25 | 0.34 |
| ProstateCancer | 0.18 | 0.86 | 0.88 | 0.83 | 0.89 |
| T2D | 0.31 | 0.96 | 0.61 | 0.95 | 0.62 |
| BreastCancer | 0.017 | 0.11 | 0.21 | 0.026 | 0.21 |
| Depression | 0.022 | 0.8 | 0.31 | 0.79 | 0.31 |
| CAD | 0.21 | 0.83 | 0.55 | 0.86 | 0.58 |
| HTN | 0.3 | 0.99 | 0.56 | 0.99 | 0.57 |
| Average | 0.21 | 0.91 | 0.5 | 0.91 | 0.5 |
| Average | 0.81 | 0.93 | 0.56 | 0.93 | 0.6 |

Supplementary Table 12: **Calibration results for 10 diseases from the UK Biobank.** We report the slope from the regression of disease status on predictions for each prediction method. The average is shown across the three well-powered traits (grey shading).

| Trait | PRS-FH$_{log}$ All | PRS-FH$_{log}$ No-sib | PRS-FH$_{log}$ Δ | PRS-FH$_{log}$ Δp | PRS-FH$_{liab}$ All | PRS-FH$_{liab}$ No-sib | PRS-FH$_{liab}$ Δ | PRS-FH$_{liab}$ Δp |
|---|---|---|---|---|---|---|---|---|
| **Non-British European** | | | | | | | | |
| LungCancer | 0.02 | 0.0053 | 0.015 | 0.063 | 0.021 | 0.0057 | 0.015 | 0.012 |
| BowelCancer | 0.013 | 0.011 | 0.0022 | 0.43 | 0.012 | 0.011 | 0.00048 | 0.82 |
| Stroke | 0.019 | 0.0036 | 0.016 | 5.5e-05 | 0.018 | 0.0032 | 0.015 | 0.00011 |
| COPD | 0.1 | 0.061 | 0.043 | 5.4e-07 | 0.11 | 0.063 | 0.051 | 8.2e-16 |
| ProstateCancer | 0.1 | 0.059 | 0.041 | 0.0019 | 0.1 | 0.065 | 0.038 | 0.0037 |
| T2D | 0.17 | 0.12 | 0.052 | 5.4e-08 | 0.18 | 0.12 | 0.057 | 1.5e-07 |
| BreastCancer | 0.027 | 0.023 | 0.0037 | 0.031 | 0.03 | 0.027 | 0.0036 | 0.12 |
| Depression | 0.071 | 0.057 | 0.014 | 1.9e-05 | 0.067 | 0.059 | 0.0083 | 0.08 |
| CAD | 0.092 | 0.059 | 0.033 | 1.9e-14 | 0.097 | 0.06 | 0.038 | 4.2e-20 |
| HTN | 0.15 | 0.11 | 0.034 | 1.4e-48 | 0.15 | 0.11 | 0.031 | 1.1e-17 |
| **South Asian** | | | | | | | | |
| LungCancer | 0 | 0 | 0 | | 0 | 0 | 0 | |
| BowelCancer | 0.0049 | 0.00067 | 0.0042 | 0.41 | 0.019 | 0.00035 | 0.019 | 0.14 |
| Stroke | 0.013 | 0.00068 | 0.013 | 0.015 | 0.013 | 0.00097 | 0.012 | 0.012 |
| COPD | 0.018 | 0.011 | 0.0073 | 0.39 | 0.015 | 0.012 | 0.0022 | 0.86 |
| ProstateCancer | 0.016 | 0.018 | -0.0021 | 0.8 | 0.035 | 0.021 | 0.014 | 0.49 |
| T2D | 0.15 | 0.078 | 0.069 | 1.3e-15 | 0.14 | 0.081 | 0.06 | 3.2e-08 |
| BreastCancer | 0.012 | 0.0048 | 0.0077 | 0.075 | 0.015 | 0.0076 | 0.0074 | 0.21 |
| Depression | 0.077 | 0.05 | 0.027 | 0.033 | 0.072 | 0.051 | 0.021 | 0.29 |
| CAD | 0.096 | 0.049 | 0.048 | 9.1e-08 | 0.087 | 0.049 | 0.038 | 2.9e-06 |
| HTN | 0.12 | 0.067 | 0.056 | 7.1e-16 | 0.11 | 0.067 | 0.042 | 3.4e-06 |
| **African** | | | | | | | | |
| LungCancer | 0.012 | 0 | 0.012 | 0.46 | 0.062 | 0 | 0.062 | 0.26 |
| BowelCancer | 0.0042 | 0 | 0.0042 | 0.38 | 0.0013 | 0 | 0.0013 | 0.51 |
| Stroke | 0.0042 | 0.00019 | 0.004 | 0.074 | 0.0085 | 0 | 0.0085 | 0.024 |
| COPD | 0.0016 | 0.00024 | 0.0014 | 0.4 | 0.0044 | 0.00047 | 0.0039 | 0.29 |
| ProstateCancer | 0.11 | 0.026 | 0.083 | 0.035 | 0.12 | 0.029 | 0.094 | 0.02 |
| T2D | 0.13 | 0.063 | 0.071 | 1e-07 | 0.13 | 0.063 | 0.063 | 1.7e-05 |
| BreastCancer | 1.1e-05 | 0 | 1.1e-05 | 0.023 | 0.0014 | 0 | 0.0014 | 0.45 |
| Depression | 0.059 | 0.033 | 0.026 | 0.047 | 0.063 | 0.033 | 0.03 | 0.095 |
| CAD | 0.029 | 0.018 | 0.011 | 0.082 | 0.019 | 0.018 | 0.0016 | 0.79 |
| HTN | 0.11 | 0.041 | 0.067 | 5.4e-18 | 0.11 | 0.041 | 0.068 | 7e-16 |

Supplementary Table 13: **Results of PRS-FH methods using parental history only for 10 diseases from the UK Biobank.** We report the prediction $R_l^2$ for both PRS-FH prediction methods including parental and sibling history (All) and parental history only (No-sib), as well as the difference in prediction $R_l^2$ (Δ) and the jackknife for the difference ($H_0 : \Delta R_o = 0$).

| Trait | PRS-FH$_{log}$ | PRS+$I(FH \geq 1)_{log}$ | Δ | Δp |
|---|---|---|---|---|
| | | Non-British European | | |
| LungCancer | 0.02 | 0.02 | -3.2e-05 | 1 |
| BowelCancer | 0.013 | 0.013 | 0.00054 | 0.84 |
| Stroke | 0.019 | 0.0077 | 0.011 | 0.0025 |
| COPD | 0.1 | 0.069 | 0.034 | 8.4e-05 |
| ProstateCancer | 0.1 | 0.075 | 0.026 | 0.018 |
| T2D | 0.17 | 0.14 | 0.029 | 0.00061 |
| BreastCancer | 0.027 | 0.026 | 0.00073 | 0.45 |
| Depression | 0.071 | 0.066 | 0.005 | 0.18 |
| CAD | 0.092 | 0.061 | 0.031 | 8.7e-12 |
| HTN | 0.15 | 0.12 | 0.025 | 1.9e-30 |
| | | South Asian | | |
| LungCancer | 0 | 0.015 | -0.015 | 0.27 |
| BowelCancer | 0.0049 | 0.0053 | -0.00039 | 0.89 |
| Stroke | 0.013 | 0.0069 | 0.0064 | 0.32 |
| COPD | 0.018 | 0.0065 | 0.011 | 0.25 |
| ProstateCancer | 0.016 | 0.019 | -0.0035 | 0.65 |
| T2D | 0.15 | 0.1 | 0.044 | 3.1e-06 |
| BreastCancer | 0.012 | 0.012 | -1.3e-05 | 0.94 |
| Depression | 0.077 | 0.08 | -0.0025 | 0.85 |
| CAD | 0.096 | 0.059 | 0.037 | 6.2e-05 |
| HTN | 0.12 | 0.075 | 0.048 | 1.5e-16 |
| | | African | | |
| LungCancer | 0.012 | 0 | 0.012 | 0.46 |
| BowelCancer | 0.0042 | 6.6e-05 | 0.0041 | 0.12 |
| Stroke | 0.0042 | 0.0014 | 0.0027 | 0.44 |
| COPD | 0.0016 | 0.0017 | -8.3e-05 | 0.96 |
| ProstateCancer | 0.11 | 0.068 | 0.041 | 0.25 |
| T2D | 0.13 | 0.076 | 0.057 | 1.2e-05 |
| BreastCancer | 1.1e-05 | 0.0021 | -0.0021 | 0.0011 |
| Depression | 0.059 | 0.069 | -0.011 | 0.49 |
| CAD | 0.029 | 0.022 | 0.0077 | 0.29 |
| HTN | 0.11 | 0.044 | 0.064 | 3.1e-20 |

Supplementary Table 14: **Results of PRS-FH$_{log}$ versus a simplified logistic regression-based method which used an indicator variable for presence of family history for 10 diseases from the UK Biobank.** We report the prediction $R_l^2$ for the PRS-FH$_{log}$ prediction method (distinct independent predictors for maternal, paternal, and sibling history) and the simplified logistic regression-based method (one predictor for a family history of disease; PRS+$I(FH \geq 1)_{log}$ , as well as the difference in prediction $R_l^2$ (Δ) and the jackknife p-value for the difference ($H_0 : \Delta R_o = 0$). $I(FH \geq 1) = 1$ if an individual reported either their mother, father, or sibling was affected; $I(FH \geq 1) = 0$ if an individual reported both mother and father were unaffected and either reported 0 relevant siblings or that 0 of their siblings were affected; otherwise $I(FH \geq 1) = NA$. Individuals with $I(FH \geq 1) = NA$ were assigned the mean $I(FH \geq 1)$ across all individuals in the 9 training folds.

| Trait | $FH_{log}$ None | $FH_{log}$ Interaction | $FH_{log}$ Δ | $FH_{log}$ Δp | PRS-$FH_{log}$ None | PRS-$FH_{log}$ Interaction | PRS-$FH_{log}$ Δ | PRS-$FH_{log}$ Δp |
|---|---|---|---|---|---|---|---|---|
| **Non-British European** | | | | | | | | |
| LungCancer | 0.016 | 0.013 | -0.003 | 0.076 | 0.02 | 0.017 | -0.0032 | 0.11 |
| BowelCancer | 0.0078 | 0.0068 | -0.00096 | 0.18 | 0.013 | 0.012 | -0.0013 | 0.15 |
| Stroke | 0.019 | 0.019 | 0.00044 | 0.69 | 0.019 | 0.02 | 0.0004 | 0.72 |
| COPD | 0.089 | 0.087 | -0.0015 | 0.092 | 0.1 | 0.1 | -0.0016 | 0.06 |
| ProstateCancer | 0.04 | 0.039 | -0.0017 | 0.054 | 0.1 | 0.099 | -0.0016 | 0.2 |
| T2D | 0.11 | 0.11 | -0.0007 | 0.0051 | 0.17 | 0.17 | -0.00079 | 0.012 |
| BreastCancer | 0.011 | 0.01 | -0.00056 | 0.17 | 0.027 | 0.026 | -0.00054 | 0.12 |
| Depression | 0.064 | 0.064 | 4.5e-05 | 0.91 | 0.071 | 0.071 | 6.1e-05 | 0.87 |
| CAD | 0.063 | 0.063 | -3.9e-07 | 1 | 0.092 | 0.092 | -3.8e-05 | 0.94 |
| HTN | 0.065 | 0.065 | 9.1e-05 | 0.65 | 0.15 | 0.15 | -2.5e-05 | 0.83 |
| **South Asian** | | | | | | | | |
| LungCancer | 0 | 0 | 0 | | 0 | 0 | 0 | |
| BowelCancer | 0.0042 | 0 | -0.0042 | 0.35 | 0.0049 | 0 | -0.0049 | 0.34 |
| Stroke | 0.014 | 0.011 | -0.003 | 0.11 | 0.013 | 0.01 | -0.0028 | 0.093 |
| COPD | 0.018 | 0.0085 | -0.0098 | 0.18 | 0.018 | 0.0081 | -0.0097 | 0.17 |
| ProstateCancer | 0.0047 | 0.0027 | -0.0021 | 0.12 | 0.016 | 0.012 | -0.0044 | 0.075 |
| T2D | 0.11 | 0.11 | -0.00028 | 0.69 | 0.15 | 0.15 | -0.00012 | 0.88 |
| BreastCancer | 0.011 | 0.012 | 0.0011 | 0.85 | 0.012 | 0.014 | 0.0015 | 0.56 |
| Depression | 0.071 | 0.069 | -0.0015 | 0.39 | 0.077 | 0.076 | -0.0015 | 0.36 |
| CAD | 0.082 | 0.081 | -0.00094 | 0.29 | 0.096 | 0.096 | -0.0008 | 0.3 |
| HTN | 0.074 | 0.074 | -0.0001 | 0.85 | 0.12 | 0.12 | -4.6e-05 | 0.94 |
| **African** | | | | | | | | |
| LungCancer | 0.012 | 0.025 | 0.013 | 0.52 | 0.012 | 0.03 | 0.018 | 0.48 |
| BowelCancer | 0.0022 | 0.00019 | -0.002 | 0.12 | 0.0042 | 0.00052 | -0.0037 | 0.14 |
| Stroke | 0.0032 | 0.0031 | -0.00015 | 0.91 | 0.0042 | 0.0042 | 5.2e-05 | 0.98 |
| COPD | 0.0017 | 0 | -0.0017 | 0.35 | 0.0016 | 0 | -0.0016 | 0.37 |
| ProstateCancer | 0.11 | 0.11 | 0.0019 | 0.92 | 0.11 | 0.11 | 0.0027 | 0.87 |
| T2D | 0.13 | 0.13 | -0.0017 | 0.078 | 0.13 | 0.13 | -0.0019 | 0.04 |
| BreastCancer | 0.00017 | 0 | -0.00017 | 0.75 | 1.1e-05 | 0 | -1.1e-05 | 0.023 |
| Depression | 0.06 | 0.056 | -0.0045 | 0.16 | 0.059 | 0.054 | -0.0043 | 0.17 |
| CAD | 0.024 | 0.023 | -0.0016 | 0.36 | 0.029 | 0.028 | -0.0015 | 0.44 |
| HTN | 0.1 | 0.1 | -0.00016 | 0.81 | 0.11 | 0.11 | -0.00018 | 0.77 |

Supplementary Table 15: **Results of $FH_{log}$ and PRS-$FH_{log}$ methods incorporating an interaction term between number of siblings and sibling disease status for 10 diseases from the UK Biobank.** We report the prediction $R_l^2$ for both $FH_{log}$ and PRS-$FH_{log}$ prediction methods not including and including an interaction term between number of siblings and disease status of siblings (None and Interaction, respectively), as well as the difference in prediction $R_l^2$ (Δ) and the jackknife for the difference ($H_0 : \Delta R_o = 0$).

| Trait | Disease prevalence when $n_s =$ 0 | 1 | 2 | 3 | 4 | 5+ |
|---|---|---|---|---|---|---|
| **Non-British European** | | | | | | |
| LungCancer | 0.0073 | 0.0055 | 0.0048 | 0.0066 | 0.0047 | 0.0084 |
| BowelCancer | 0.0098 | 0.011 | 0.0099 | 0.012 | 0.013 | 0.014 |
| Stroke | 0.017 | 0.017 | 0.02 | 0.019 | 0.021 | 0.029 |
| COPD | 0.039 | 0.025 | 0.027 | 0.03 | 0.04 | 0.061 |
| ProstateCancer | 0.036 | 0.027 | 0.034 | 0.034 | 0.03 | 0.037 |
| T2D | 0.046 | 0.034 | 0.037 | 0.039 | 0.041 | 0.047 |
| BreastCancer | 0.061 | 0.059 | 0.063 | 0.065 | 0.06 | 0.055 |
| Depression | 0.077 | 0.072 | 0.074 | 0.072 | 0.07 | 0.081 |
| CAD | 0.087 | 0.068 | 0.065 | 0.072 | 0.077 | 0.11 |
| HTN | 0.33 | 0.27 | 0.28 | 0.27 | 0.29 | 0.33 |
| **South Asian** | | | | | | |
| LungCancer | 0 | 0.0019 | 0 | 0.0035 | 0.0027 | 0.0019 |
| BowelCancer | 0 | 0.0077 | 0.0087 | 0.0061 | 0.0027 | 0.0045 |
| Stroke | 0.017 | 0.027 | 0.014 | 0.023 | 0.02 | 0.031 |
| COPD | 0.041 | 0.014 | 0.023 | 0.017 | 0.019 | 0.025 |
| ProstateCancer | 0.025 | 0.014 | 0.02 | 0.012 | 0.0069 | 0.014 |
| T2D | 0.16 | 0.13 | 0.12 | 0.13 | 0.15 | 0.18 |
| BreastCancer | 0 | 0.052 | 0.029 | 0.039 | 0.041 | 0.034 |
| Depression | 0.033 | 0.046 | 0.029 | 0.058 | 0.05 | 0.062 |
| CAD | 0.12 | 0.085 | 0.12 | 0.11 | 0.14 | 0.17 |
| HTN | 0.35 | 0.29 | 0.32 | 0.36 | 0.37 | 0.41 |
| **African** | | | | | | |
| LungCancer | 0.0021 | 0.0037 | 0.0077 | 0.0048 | 0.0023 | 0.0026 |
| BowelCancer | 0.0042 | 0.013 | 0.0077 | 0.0072 | 0.013 | 0.0084 |
| Stroke | 0.023 | 0.019 | 0.023 | 0.028 | 0.016 | 0.027 |
| COPD | 0.021 | 0.013 | 0.011 | 0.017 | 0.01 | 0.011 |
| ProstateCancer | 0.062 | 0.045 | 0.041 | 0.034 | 0.043 | 0.057 |
| T2D | 0.12 | 0.092 | 0.087 | 0.076 | 0.088 | 0.1 |
| BreastCancer | 0.039 | 0.019 | 0.039 | 0.021 | 0.028 | 0.029 |
| Depression | 0.067 | 0.033 | 0.048 | 0.043 | 0.042 | 0.042 |
| CAD | 0.069 | 0.041 | 0.049 | 0.049 | 0.06 | 0.069 |
| HTN | 0.47 | 0.4 | 0.38 | 0.4 | 0.42 | 0.44 |

Supplementary Table 16: **Values of disease prevalence as a function of number of siblings for 10 diseases from the UK Biobank.** Across 10 diseases we report the disease prevalence among individuals with varying number of siblings (0,1,. . .,4,5+).

| | FH$_{log}$ | | | | PRS-FH$_{log}$ | | | |
|---|---|---|---|---|---|---|---|---|
| Trait | N$_{sib}$ | $N_{sib}^{+}$ | Δ | Δp | N$_{sib}$ | $N_{sib}^{+}$ | Δ | Δ p |
| **Non-British European** | | | | | | | | |
| LungCancer | 0.016 | 0.017 | -0.00044 | 0.87 | 0.02 | 0.021 | -0.00055 | 0.86 |
| BowelCancer | 0.0078 | 0.0057 | 0.0021 | 0.0029 | 0.013 | 0.011 | 0.0024 | 0.0047 |
| Stroke | 0.019 | 0.018 | 0.0011 | 0.26 | 0.019 | 0.018 | 0.0011 | 0.25 |
| COPD | 0.089 | 0.1 | -0.011 | 0.00014 | 0.1 | 0.11 | -0.012 | 0.00027 |
| ProstateCancer | 0.04 | 0.042 | -0.0015 | 0.44 | 0.1 | 0.1 | -0.00052 | 0.83 |
| T2D | 0.11 | 0.11 | -0.002 | 0.34 | 0.17 | 0.17 | -0.0033 | 0.21 |
| BreastCancer | 0.011 | 0.0091 | 0.0015 | 0.042 | 0.027 | 0.026 | 0.0011 | 0.15 |
| Depression | 0.064 | 0.066 | -0.0018 | 0.095 | 0.071 | 0.073 | -0.0018 | 0.095 |
| CAD | 0.063 | 0.067 | -0.0048 | 0.0009 | 0.092 | 0.096 | -0.0044 | 0.0053 |
| HTN | 0.065 | 0.073 | -0.0078 | 4.5e-11 | 0.15 | 0.15 | -0.0065 | 1.5e-08 |
| **South Asian** | | | | | | | | |
| LungCancer | 0 | 0 | 0 | | 0 | 0 | 0 | |
| BowelCancer | 0.0042 | 0.012 | -0.0082 | 0.17 | 0.0049 | 0.014 | -0.0095 | 0.12 |
| Stroke | 0.014 | 0.013 | 0.0018 | 0.51 | 0.013 | 0.012 | 0.0014 | 0.6 |
| COPD | 0.018 | 0.019 | -0.00042 | 0.93 | 0.018 | 0.018 | -0.00034 | 0.94 |
| ProstateCancer | 0.0047 | 0.0041 | 0.0006 | 0.85 | 0.016 | 0.017 | -0.00069 | 0.88 |
| T2D | 0.11 | 0.11 | -0.0022 | 0.42 | 0.15 | 0.15 | -0.0024 | 0.42 |
| BreastCancer | 0.011 | 0.013 | -0.0021 | 0.69 | 0.012 | 0.015 | -0.0021 | 0.5 |
| Depression | 0.071 | 0.074 | -0.0033 | 0.39 | 0.077 | 0.08 | -0.0024 | 0.54 |
| CAD | 0.082 | 0.081 | 0.0011 | 0.47 | 0.096 | 0.095 | 0.0013 | 0.43 |
| HTN | 0.074 | 0.075 | -0.00071 | 0.65 | 0.12 | 0.12 | -0.00017 | 0.9 |
| **African** | | | | | | | | |
| LungCancer | 0.012 | 0.014 | -0.002 | 0.91 | 0.012 | 0.015 | -0.0027 | 0.92 |
| BowelCancer | 0.0022 | 0 | 0.0022 | 0.024 | 0.0042 | 7.7e-05 | 0.0041 | 0.066 |
| Stroke | 0.0032 | 0.0023 | 0.00099 | 0.55 | 0.0042 | 0.0031 | 0.0011 | 0.55 |
| COPD | 0.0017 | 0.0011 | 0.00054 | 0.7 | 0.0016 | 0.0013 | 0.00035 | 0.8 |
| ProstateCancer | 0.11 | 0.099 | 0.0065 | 0.36 | 0.11 | 0.1 | 0.0048 | 0.5 |
| T2D | 0.13 | 0.13 | -0.001 | 0.82 | 0.13 | 0.13 | -0.001 | 0.83 |
| BreastCancer | 0.00017 | 2.6e-10 | 0.00017 | 2.4e-05 | 1.1e-05 | 0 | 1.1e-05 | 0.038 |
| Depression | 0.06 | 0.062 | -0.002 | 0.62 | 0.059 | 0.061 | -0.002 | 0.61 |
| CAD | 0.024 | 0.023 | 0.0011 | 0.69 | 0.029 | 0.027 | 0.0019 | 0.54 |
| HTN | 0.1 | 0.1 | -0.003 | 0.13 | 0.11 | 0.11 | -0.0031 | 0.12 |

Supplementary Table 17: **Results of FH$_{log}$ and PRS-FH$_{log}$ methods that account for variation in disease prevalence as a function of number of siblings for 10 diseases from the UK Biobank.** We report the prediction $R_l^2$ for both FH$_{log}$ and PRS-FH$_{log}$ prediction methods including a continuous variable reflecting the total number of relevant siblings an individual has (N$_{sib}$) and a model that additionally includes 5 indicator variables reflecting 1, 2, 3, 4, or 5 or more total siblings ($N_{sib}^{+}$), as well as the difference in prediction $R_l^2$ (Δ) and the jackknife for the difference ($H_0 : \Delta R_o = 0$).

| Trait | FH$_{liab}$ T$_{sib}$ | FH$_{liab}$ T$_{nosib}$ | FH$_{liab}$ Δ | FH$_{liab}$ Δp | PRS-FH$_{liab}$ T$_{sib}$ | PRS-FH$_{liab}$ T$_{nosib}$ | PRS-FH$_{liab}$ Δ | PRS-FH$_{liab}$ Δ p |
|---|---|---|---|---|---|---|---|---|
| **Non-British European** | | | | | | | | |
| LungCancer | 0.017 | 0.014 | 0.0028 | 0.46 | 0.021 | 0.018 | 0.0033 | 0.42 |
| BowelCancer | 0.0062 | 0.0068 | -0.00059 | 0.69 | 0.012 | 0.012 | -0.00089 | 0.6 |
| Stroke | 0.018 | 0.014 | 0.0042 | 0.017 | 0.018 | 0.014 | 0.0043 | 0.016 |
| COPD | 0.098 | 0.066 | 0.032 | 1.3e-11 | 0.11 | 0.083 | 0.032 | 3.7e-10 |
| ProstateCancer | 0.04 | 0.039 | 0.00058 | 0.72 | 0.1 | 0.1 | 0.0011 | 0.63 |
| T2D | 0.12 | 0.11 | 0.0038 | 0.019 | 0.18 | 0.17 | 0.0053 | 0.011 |
| BreastCancer | 0.01 | 0.012 | -0.0013 | 0.05 | 0.03 | 0.032 | -0.0012 | 0.1 |
| Depression | 0.061 | 0.06 | 0.00048 | 0.23 | 0.067 | 0.066 | 0.00056 | 0.19 |
| CAD | 0.068 | 0.057 | 0.011 | 1.3e-09 | 0.097 | 0.086 | 0.011 | 2.4e-07 |
| HTN | 0.071 | 0.066 | 0.0053 | 4.1e-18 | 0.15 | 0.14 | 0.0057 | 5.7e-14 |
| **South Asian** | | | | | | | | |
| LungCancer | 0 | 0 | 0 | | 0 | 0 | 0 | |
| BowelCancer | 0.019 | 0.013 | 0.006 | 0.52 | 0.019 | 0.013 | 0.006 | 0.52 |
| Stroke | 0.013 | 0.0066 | 0.0066 | 0.098 | 0.013 | 0.0066 | 0.0066 | 0.098 |
| COPD | 0.015 | 0.014 | 0.00061 | 0.74 | 0.015 | 0.014 | 0.00061 | 0.74 |
| ProstateCancer | 0.023 | 0.021 | 0.0016 | 0.81 | 0.035 | 0.032 | 0.0033 | 0.67 |
| T2D | 0.11 | 0.11 | 0.0037 | 0.15 | 0.14 | 0.14 | 0.0039 | 0.17 |
| BreastCancer | 0.0094 | 0.0094 | 5.6e-05 | 0.98 | 0.015 | 0.016 | -0.00069 | 0.91 |
| Depression | 0.069 | 0.065 | 0.0045 | 0.1 | 0.072 | 0.068 | 0.0044 | 0.11 |
| CAD | 0.073 | 0.06 | 0.013 | 0.003 | 0.087 | 0.075 | 0.012 | 0.0063 |
| HTN | 0.069 | 0.06 | 0.0097 | 1.2e-08 | 0.11 | 0.098 | 0.01 | 2.4e-07 |
| **African** | | | | | | | | |
| LungCancer | 0.062 | 0.041 | 0.021 | 0.62 | 0.062 | 0.041 | 0.021 | 0.62 |
| BowelCancer | 0.0013 | 0.0073 | -0.006 | 0.12 | 0.0013 | 0.0073 | -0.006 | 0.12 |
| Stroke | 0.0091 | 0.0096 | -0.00047 | 0.83 | 0.0085 | 0.0088 | -0.0003 | 0.89 |
| COPD | 0.0044 | 0.003 | 0.0014 | 0.57 | 0.0044 | 0.003 | 0.0014 | 0.57 |
| ProstateCancer | 0.12 | 0.12 | -0.0047 | 0.63 | 0.12 | 0.13 | -0.0038 | 0.71 |
| T2D | 0.12 | 0.12 | -0.00095 | 0.67 | 0.13 | 0.13 | -0.0013 | 0.58 |
| BreastCancer | 0.0014 | 0.0035 | -0.0022 | 0.32 | 0.0014 | 0.0035 | -0.0022 | 0.32 |
| Depression | 0.063 | 0.062 | 0.0013 | 0.48 | 0.063 | 0.062 | 0.0013 | 0.48 |
| CAD | 0.017 | 0.012 | 0.0044 | 0.042 | 0.019 | 0.015 | 0.0044 | 0.064 |
| HTN | 0.1 | 0.1 | 0.0024 | 0.018 | 0.11 | 0.11 | 0.0025 | 0.015 |

Supplementary Table 18: **Results of FH$_{liab}$ and PRS-FH$_{liab}$ methods that do not account for variation in disease prevalence as a function of number of siblings for 10 diseases from the UK Biobank.** We report the prediction $R_l^2$ for both FH$_{liab}$ and PRS-FH$_{liab}$ prediction methods including a disease threshold that varies as a function of the number of siblings an individual has (T$_{sib}$) and a model that does not (T$_{nosib}$), as well as the difference in prediction $R_l^2$ (Δ) and the jackknife for the difference ($H_0 : \Delta R_o = 0$).

| Trait | Covar $R_l^2$ | Covar p | PCs $R_l^2$ | PCs p | Age $R_l^2$ | Age p | BMI $R_l^2$ | BMI p | Sex $R_l^2$ | Sex p |
|---|---|---|---|---|---|---|---|---|---|---|
| **Non-British European** | | | | | | | | | | |
| LungCancer | 0.055 | 7.2e-18 | 0.0014 | 0.045 | 0.06 | 1.7e-24 | 0 | | 0.0014 | 0.027 |
| BowelCancer | 0.074 | 6.5e-32 | 0.0037 | 0.00077 | 0.068 | 5.8e-40 | 0.00059 | 0.081 | 0.0068 | 1.4e-06 |
| Stroke | 0.11 | 4e-70 | 0.0055 | 6.1e-08 | 0.07 | 3.6e-71 | 0.017 | 5.4e-21 | 0.012 | 2.7e-14 |
| COPD | 0.1 | 3.5e-117 | 0.016 | 1.7e-23 | 0.057 | 9.6e-112 | 0.026 | 1.2e-21 | 0.0038 | 1.4e-09 |
| ProstateCancer | 0.16 | 2.3e-85 | 0.0004 | 0.13 | 0.17 | 6e-115 | 0 | | | |
| T2D | 0.32 | 1.4e-170 | 0.0051 | 2.3e-10 | 0.057 | 7.4e-105 | 0.17 | 6.7e-87 | 0.036 | 1.7e-78 |
| BreastCancer | 0.041 | 1.6e-47 | 4.9e-05 | 0.29 | 0.046 | 2.5e-55 | 0.00019 | 0.13 | | |
| Depression | 0.033 | 3.4e-73 | 0.0007 | 0.0043 | 0.0012 | 4.2e-05 | 0.018 | 2.8e-34 | 0.0082 | 1.7e-28 |
| CAD | 0.23 | 0 | 0.0064 | 1.9e-19 | 0.11 | 0 | 0.044 | 5.5e-78 | 0.054 | 8.2e-168 |
| HTN | 0.26 | 0 | 0.0024 | 3e-14 | 0.13 | 0 | 0.13 | 0 | 0.019 | 3.5e-109 |
| Average | 0.2 | | 0.0028 | | 0.062 | | 0.11 | | 0.021 | |
| **South Asian** | | | | | | | | | | |
| LungCancer | 0 | | 0 | | 0.027 | 0.018 | 0 | | 0.0054 | 0.18 |
| BowelCancer | 0.0011 | 0.28 | 0 | | 0.035 | 0.0012 | 0 | | 0 | |
| Stroke | 0.082 | 4.6e-10 | 0.0073 | 0.063 | 0.066 | 6.2e-15 | 0.0065 | 0.0076 | 0.01 | 0.0012 |
| COPD | 0.054 | 3.2e-12 | 0 | | 0.067 | 1.9e-11 | 0.0041 | 0.086 | 0.0048 | 0.018 |
| ProstateCancer | 0.07 | 1e-06 | 0.0015 | 0.2 | 0.17 | 1.6e-10 | 0 | | | |
| T2D | 0.19 | 2.8e-111 | 0.0003 | 0.19 | 0.11 | 2.1e-87 | 0.045 | 4.7e-19 | 0.024 | 4.2e-23 |
| BreastCancer | 0 | | 0 | | 0.011 | 0.003 | 0 | | | |
| Depression | 0.02 | 2.1e-06 | 0 | | 0 | | 0.023 | 4.9e-05 | 0.0089 | 5.9e-06 |
| CAD | 0.23 | 2.8e-115 | 0.0025 | 0.0026 | 0.14 | 7.5e-81 | 0.019 | 4.9e-15 | 0.055 | 4.4e-48 |
| HTN | 0.29 | 0 | 0.0006 | 0.058 | 0.21 | 6.3e-223 | 0.067 | 3e-64 | 0.011 | 1.7e-12 |
| Average | 0.17 | | 0.0003 | | 0.11 | | 0.045 | | 0.015 | |
| **African** | | | | | | | | | | |
| LungCancer | 0.014 | 0.033 | 0.00015 | 0.59 | 0.048 | 0.0074 | 0.00061 | 0.39 | 0.00074 | 0.39 |
| BowelCancer | 0.017 | 0.0023 | 0.00039 | 0.27 | 0.053 | 2.8e-06 | 0 | | 0 | |
| Stroke | 0.066 | 3.8e-09 | 0 | 0.98 | 0.078 | 1.3e-10 | 0.0052 | 0.03 | 0 | 1 |
| COPD | 0.0052 | 0.037 | 0 | | 0.024 | 0.00063 | 0.0003 | 0.34 | 0 | |
| ProstateCancer | 0.3 | 1.3e-26 | 0.0023 | 0.055 | 0.3 | 4.3e-30 | 0.0005 | 0.25 | | |
| T2D | 0.17 | 6.1e-75 | 0.0024 | 0.01 | 0.12 | 8.6e-50 | 0.045 | 1e-20 | 0.0069 | 3.7e-05 |
| BreastCancer | 0 | | 0 | | 0.012 | 0.0022 | 0 | | | |
| Depression | 0.022 | 1e-06 | 0.0036 | 0.023 | 0.0014 | 0.051 | 0.0069 | 0.002 | 0.012 | 6.2e-06 |
| CAD | 0.13 | 8.3e-36 | 0.001 | 0.081 | 0.11 | 6.3e-28 | 0.018 | 1.7e-08 | 0.00043 | 0.19 |
| HTN | 0.26 | 0 | 0.014 | 1.3e-20 | 0.19 | 1.4e-209 | 0.076 | 3.4e-88 | 0.0013 | 0.0029 |
| Average | 0.15 | | 0.0065 | | 0.1 | | 0.043 | | 0.0069 | |

Supplementary Table 19: **Results of predictions using covariates alone for 10 diseases from the UK Biobank.** The prediction $R_l^2$ is shown for models based on covariates alone: 20 PCs (PCs), age, BMI, Sex, or 20 PCs, age, BMI, and sex (Covar), as well as the jackknife p-value for the test $H_o : R_o > 0$.The average is shown across the three well-powered traits (grey shading).

**(a)**

| | $PRS^{+}$ | | $FH^{+}_{log}$ | | $FH^{+}_{liab}$ | | $PRS\text{-}FH^{+}_{log}$ | | $PRS\text{-}FH^{+}_{liab}$ | |
|---|---|---|---|---|---|---|---|---|---|---|
| Trait | $R^2_l$ | p | $R^2_l$ | p | $R^2_l$ | p | $R^2_l$ | p | $R^2_l$ | p |
| Non-British European | | | | | | | | | | |
| LungCancer | 0.059 | 1e-18 | 0.064 | 1.9e-13 | 0.067 | 7.2e-14 | 0.07 | 2.6e-13 | 0.072 | 5.9e-14 |
| BowelCancer | 0.082 | 7.6e-33 | 0.077 | 1.4e-31 | 0.076 | 2.2e-30 | 0.085 | 5.2e-34 | 0.084 | 1.3e-32 |
| Stroke | 0.11 | 2.4e-71 | 0.12 | 2.2e-69 | 0.11 | 1.2e-55 | 0.12 | 7.9e-71 | 0.11 | 6.3e-56 |
| COPD | 0.12 | 3.7e-130 | 0.17 | 1.6e-105 | 0.18 | 7.1e-113 | 0.19 | 2.4e-117 | 0.2 | 1.2e-125 |
| ProstateCancer | 0.25 | 4.2e-74 | 0.2 | 3.2e-69 | 0.19 | 3.1e-53 | 0.29 | 5e-78 | 0.28 | 7.4e-67 |
| T2D | 0.44 | 6.6e-215 | 0.46 | 5.2e-238 | 0.45 | 1.3e-232 | 0.57 | 4.5e-291 | 0.53 | 4e-288 |
| BreastCancer | 0.068 | 5.4e-59 | 0.052 | 9.8e-53 | 0.05 | 4.2e-54 | 0.077 | 1.3e-59 | 0.077 | 3.4e-65 |
| Depression | 0.041 | 1.7e-101 | 0.093 | 5.1e-137 | 0.088 | 1.3e-123 | 0.1 | 1.1e-156 | 0.094 | 3.7e-136 |
| CAD | 0.28 | 0 | 0.29 | 0 | 0.28 | 0 | 0.34 | 0 | 0.32 | 0 |
| HTN | 0.35 | 0 | 0.32 | 0 | 0.3 | 0 | 0.4 | 0 | 0.37 | 0 |
| Average | 0.28 | | 0.29 | | 0.28 | | 0.35 | | 0.33 | |
| South Asian | | | | | | | | | | |
| LungCancer | 0 | | 0 | | 0 | | 0 | | 0 | |
| BowelCancer | 0.0015 | 0.24 | 0.0088 | 0.093 | 0.019 | 0.062 | 0.0094 | 0.084 | 0.019 | 0.062 |
| Stroke | 0.079 | 3.2e-10 | 0.085 | 3.1e-10 | 0.091 | 2e-10 | 0.082 | 2.6e-10 | 0.091 | 2e-10 |
| COPD | 0.056 | 4.7e-12 | 0.052 | 9.9e-09 | 0.024 | 0.00025 | 0.053 | 9.3e-09 | 0.024 | 0.00025 |
| ProstateCancer | 0.12 | 4.9e-06 | 0.067 | 1.5e-05 | 0.098 | 0.00025 | 0.13 | 1.4e-05 | 0.14 | 8.4e-05 |
| T2D | 0.25 | 1.1e-134 | 0.3 | 1.4e-166 | 0.28 | 1.1e-158 | 0.35 | 1.8e-185 | 0.32 | 1.1e-176 |
| BreastCancer | 3.4e-05 | 0.63 | 0.00045 | 0.29 | 0.00079 | 0.22 | 0.0025 | 0.098 | 0.0039 | 0.051 |
| Depression | 0.025 | 5.6e-07 | 0.091 | 1.2e-13 | 0.087 | 7.1e-14 | 0.099 | 1.5e-13 | 0.09 | 7.1e-14 |
| CAD | 0.26 | 2.8e-124 | 0.3 | 1.8e-155 | 0.29 | 6.6e-147 | 0.32 | 2.4e-162 | 0.31 | 3.3e-151 |
| HTN | 0.36 | 0 | 0.34 | 0 | 0.31 | 0 | 0.4 | 0 | 0.35 | 0 |
| Average | 0.21 | | 0.25 | | 0.23 | | 0.28 | | 0.26 | |
| African | | | | | | | | | | |
| LungCancer | 0.014 | 0.037 | 0.0056 | 0.059 | 0.034 | 0.067 | 0.0053 | 0.062 | 0.034 | 0.067 |
| BowelCancer | 0.02 | 0.0035 | 0.019 | 0.028 | 0.019 | 0.012 | 0.025 | 0.037 | 0.019 | 0.012 |
| Stroke | 0.065 | 5.6e-09 | 0.064 | 1.1e-08 | 0.064 | 5e-08 | 0.063 | 2e-08 | 0.063 | 8.8e-08 |
| COPD | 0.0045 | 0.046 | 0.008 | 0.024 | 0.0086 | 0.021 | 0.0074 | 0.026 | 0.0086 | 0.021 |
| ProstateCancer | 0.32 | 6.2e-26 | 0.36 | 2.3e-24 | 0.36 | 9.9e-23 | 0.37 | 8.9e-25 | 0.37 | 3.1e-23 |
| T2D | 0.18 | 8e-82 | 0.32 | 9e-134 | 0.28 | 2.5e-102 | 0.32 | 4.8e-138 | 0.29 | 8.4e-108 |
| BreastCancer | 0 | | 0 | | 0.00026 | 0.44 | 0 | | 0.00026 | 0.44 |
| Depression | 0.021 | 1.2e-06 | 0.096 | 1.6e-11 | 0.088 | 8e-11 | 0.094 | 1.7e-11 | 0.088 | 8e-11 |
| CAD | 0.13 | 3e-36 | 0.15 | 1.8e-39 | 0.13 | 7.8e-35 | 0.15 | 3e-40 | 0.13 | 2.9e-35 |
| HTN | 0.27 | 0 | 0.34 | 0 | 0.31 | 0 | 0.34 | 0 | 0.31 | 0 |
| Average | 0.15 | | 0.25 | | 0.22 | | 0.25 | | 0.23 | |

**(b)**

| Trait | PRS$^+$ $R^2_l$ | p | $FH^+_{log}$ $R^2_l$ | p | $FH^+_{liab}$ $R^2_l$ | p | PRS-$FH^+_{log}$ $R^2_l$ | p | PRS-$FH^+_{liab}$ $R^2_l$ | p |
|---|---|---|---|---|---|---|---|---|---|---|
| Non-British European | | | | | | | | | | |
| LungCancer | 0.0038 (0.003) | 0.26 | 0.0087 (0.01) | 0.35 | 0.012 (0.01) | 0.2 | 0.014 (0.01) | 0.2 | 0.017 (0.01) | 0.11 |
| BowelCancer | 0.0083 (0.005) | 0.077 | 0.0029 (0.005) | 0.53 | 0.0022 (0.006) | 0.7 | 0.011 (0.006) | 0.067 | 0.0098 (0.007) | 0.13 |
| Stroke | -3e-06 (0.001) | 0.99 | 0.008 (0.006) | 0.2 | 0.002 (0.009) | 0.83 | 0.0084 (0.006) | 0.17 | 0.0017 (0.009) | 0.85 |
| COPD | 0.017 (0.005) | 0.00016 | 0.072 (0.01) | 2.3e-10 | 0.076 (0.01) | 8.7e-11 | 0.088 (0.01) | 2.8e-13 | 0.093 (0.01) | 1.4e-14 |
| ProstateCancer | 0.086 (0.02) | 2e-07 | 0.035 (0.01) | 0.014 | 0.029 (0.02) | 0.12 | 0.13 (0.02) | 5.8e-10 | 0.12 (0.03) | 3.5e-08 |
| T2D | 0.12 (0.01) | 1.7e-21 | 0.14 (0.02) | 6.1e-19 | 0.13 (0.02) | 1.6e-13 | 0.25 (0.02) | 1.8e-37 | 0.22 (0.02) | 1.9e-31 |
| BreastCancer | 0.027 (0.006) | 4.6e-06 | 0.011 (0.003) | 0.00024 | 0.0097 (0.004) | 0.014 | 0.036 (0.007) | 7.4e-08 | 0.036 (0.007) | 2e-07 |
| Depression | 0.0086 (0.001) | 9e-09 | 0.06 (0.006) | 5e-27 | 0.055 (0.007) | 1.6e-19 | 0.067 (0.006) | 2.7e-32 | 0.061 (0.007) | 2.9e-23 |
| CAD | 0.053 (0.005) | 1.9e-26 | 0.062 (0.007) | 1.2e-20 | 0.056 (0.008) | 2.7e-13 | 0.11 (0.008) | 1.5e-45 | 0.096 (0.009) | 1.1e-31 |
| HTN | 0.093 (0.004) | 1e-123 | 0.062 (0.004) | 2e-69 | 0.042 (0.005) | 7.7e-20 | 0.14 (0.005) | 5.2e-191 | 0.11 (0.005) | 8.4e-113 |
| Average | 0.074 | | 0.088 | | 0.075 | | 0.15 | | 0.13 | |
| South Asian | | | | | | | | | | |
| LungCancer | 0 (0) | | 0 (0) | | 0 (0) | | 0 (0) | | 0 (0) | |
| BowelCancer | 0.00038 (9e-04) | 0.63 | 0.0077 (0.009) | 0.24 | 0.018 (0.02) | 0.2 | 0.0083 (0.01) | 0.23 | 0.018 (0.02) | 0.2 |
| Stroke | -0.0028 (0.003) | 0.29 | 0.0038 (0.01) | 0.71 | 0.0092 (0.01) | 0.4 | 0.00047 (0.01) | 0.94 | 0.0092 (0.01) | 0.4 |
| COPD | 0.0015 (0.002) | 0.5 | -0.0017 (0.01) | 0.86 | -0.03 (0.02) | 0.11 | -0.0011 (0.01) | 0.89 | -0.03 (0.02) | 0.11 |
| ProstateCancer | 0.052 (0.03) | 0.062 | -0.002 (0.02) | 0.92 | 0.029 (0.05) | 0.52 | 0.056 (0.04) | 0.14 | 0.069 (0.06) | 0.18 |
| T2D | 0.065 (0.01) | 7.2e-10 | 0.11 (0.02) | 1.5e-15 | 0.094 (0.02) | 2e-07 | 0.16 (0.02) | 1.7e-23 | 0.13 (0.02) | 1.7e-12 |
| BreastCancer | 3.4e-05 (3e-04) | 0.73 | 0.00045 (0.001) | 0.58 | 0.00079 (0.002) | 0.44 | 0.0025 (0.004) | 0.2 | 0.0039 (0.004) | 0.1 |
| Depression | 0.005 (0.003) | 0.057 | 0.072 (0.02) | 7.8e-06 | 0.067 (0.02) | 0.00024 | 0.08 (0.02) | 3e-06 | 0.07 (0.02) | 0.00017 |
| CAD | 0.027 (0.007) | 6e-05 | 0.066 (0.01) | 2.2e-07 | 0.056 (0.01) | 2.3e-05 | 0.09 (0.01) | 4.1e-10 | 0.077 (0.01) | 1.2e-07 |
| HTN | 0.071 (0.008) | 1.9e-18 | 0.055 (0.007) | 3.5e-14 | 0.022 (0.01) | 0.043 | 0.11 (0.01) | 6.2e-29 | 0.067 (0.01) | 9.9e-09 |
| Average | 0.047 | | 0.08 | | 0.061 | | 0.12 | | 0.091 | |
| African | | | | | | | | | | |
| LungCancer | -0.00023 (0.002) | 0.9 | -0.0085 (0.008) | 0.052 | 0.02 (0.03) | 0.53 | -0.0087 (0.008) | 0.049 | 0.02 (0.03) | 0.53 |
| BowelCancer | 0.003 (0.004) | 0.36 | 0.0019 (0.01) | 0.87 | 0.0025 (0.008) | 0.78 | 0.0082 (0.02) | 0.65 | 0.0025 (0.008) | 0.78 |
| Stroke | -0.00039 (0.004) | 0.91 | -0.0022 (0.007) | 0.76 | -0.0021 (0.01) | 0.88 | -0.0031 (0.008) | 0.7 | -0.003 (0.01) | 0.83 |
| COPD | -0.00072 (7e-04) | 0.24 | 0.0028 (0.004) | 0.54 | 0.0033 (0.005) | 0.52 | 0.0022 (0.004) | 0.62 | 0.0033 (0.005) | 0.52 |
| ProstateCancer | 0.016 (0.01) | 0.22 | 0.053 (0.04) | 0.22 | 0.056 (0.05) | 0.3 | 0.069 (0.05) | 0.13 | 0.071 (0.06) | 0.18 |
| T2D | 0.0099 (0.004) | 0.023 | 0.15 (0.02) | 7e-14 | 0.11 (0.02) | 2.6e-06 | 0.16 (0.02) | 1e-14 | 0.12 (0.02) | 6.5e-07 |
| BreastCancer | 0 (0) | | 0 (0) | | 0.00026 (0.001) | 0.89 | 0 (0) | | 0.00026 (0.001) | 0.89 |
| Depression | -0.00055 (8e-04) | 0.46 | 0.074 (0.03) | 9.1e-05 | 0.067 (0.03) | 0.0019 | 0.072 (0.02) | 0.00011 | 0.067 (0.03) | 0.0019 |
| CAD | 0.0032 (0.004) | 0.42 | 0.022 (0.01) | 0.049 | 0.0016 (0.01) | 0.91 | 0.027 (0.01) | 0.027 | 0.0054 (0.01) | 0.71 |
| HTN | 0.0091 (0.003) | 0.002 | 0.078 (0.007) | 9e-26 | 0.049 (0.01) | 7e-06 | 0.085 (0.008) | 2.3e-27 | 0.055 (0.01) | 4.1e-07 |
| Average | 0.0062 | | 0.1 | | 0.076 | | 0.11 | | 0.08 | |

**(c)**

| Trait | PRS-FH$^+_{log}$ $R^2_l$ | p | $\Delta_{PRS}$ p | $\Delta_{FH}$ p | PRS-FH$^+_{liab}$ $R^2_l$ | p | $\Delta_{PRS}$ p | $\Delta_{FH}$ p | $\Delta_{liab,log}$p |
|---|---|---|---|---|---|---|---|---|---|
| **Non-British European** | | | | | | | | | |
| LungCancer | 0.014 (0.01) | 0.2 | 0.29 | 0.14 | 0.017 (0.01) | 0.11 | 0.17 | 0.11 | 0.46 |
| BowelCancer | 0.011 (0.006) | 0.067 | 0.54 | 0.081 | 0.0098 (0.007) | 0.13 | 0.81 | 0.035 | 0.58 |
| Stroke | 0.0084 (0.006) | 0.17 | 0.17 | 0.75 | 0.0017 (0.009) | 0.85 | 0.86 | 0.38 | 0.16 |
| COPD | 0.088 (0.01) | 2.8e-13 | 1.3e-09 | 0.0015 | 0.093 (0.01) | 1.4e-14 | 6e-11 | 3.5e-05 | 0.35 |
| ProstateCancer | 0.13 (0.02) | 5.8e-10 | 0.0035 | 6e-08 | 0.12 (0.03) | 3.5e-08 | 0.088 | 4.3e-11 | 0.51 |
| T2D | 0.25 (0.02) | 1.8e-37 | 4.4e-13 | 8.3e-16 | 0.22 (0.02) | 1.9e-31 | 6.3e-07 | 1.3e-20 | 0.00012 |
| BreastCancer | 0.036 (0.007) | 7.4e-08 | 0.0011 | 2.8e-05 | 0.036 (0.007) | 2e-07 | 0.024 | 6.7e-07 | 0.99 |
| Depression | 0.067 (0.006) | 2.7e-32 | 8.7e-25 | 2.2e-05 | 0.061 (0.007) | 2.9e-23 | 7.1e-17 | 5.3e-10 | 0.054 |
| CAD | 0.11 (0.008) | 1.5e-45 | 6.5e-17 | 6.3e-21 | 0.096 (0.009) | 1.1e-31 | 4e-08 | 4.5e-28 | 2e-06 |
| HTN | 0.14 (0.005) | 5.2e-191 | 1.6e-55 | 4.4e-104 | 0.11 (0.005) | 8.4e-113 | 7.2e-07 | 8.7e-141 | 5.7e-28 |
| Average | 0.15 | | | | 0.13 | | | | |
| **South Asian** | | | | | | | | | |
| LungCancer | 0 (0) | | | | 0 (0) | | | | |
| BowelCancer | 0.0083 (0.01) | 0.23 | 0.24 | 0.81 | 0.018 (0.02) | 0.2 | 0.2 | | 0.23 |
| Stroke | 0.00047 (0.01) | 0.94 | 0.74 | 0.19 | 0.0092 (0.01) | 0.4 | 0.28 | | 0.2 |
| COPD | -0.0011 (0.01) | 0.89 | 0.8 | 0.79 | -0.03 (0.02) | 0.11 | 0.1 | | 0.052 |
| ProstateCancer | 0.056 (0.04) | 0.14 | 0.91 | 0.035 | 0.069 (0.06) | 0.18 | 0.74 | 0.084 | 0.65 |
| T2D | 0.16 (0.02) | 1.7e-23 | 6.7e-13 | 1.3e-07 | 0.13 (0.02) | 1.7e-12 | 0.00016 | 7.3e-11 | 0.00067 |
| BreastCancer | 0.0025 (0.004) | 0.2 | 0.019 | 0.2 | 0.0039 (0.004) | 0.1 | 0.0057 | 0.054 | 0.36 |
| Depression | 0.08 (0.02) | 3e-06 | 8.1e-06 | 0.033 | 0.07 (0.02) | 0.00017 | 0.00066 | 0.027 | 0.47 |
| CAD | 0.09 (0.01) | 4.1e-10 | 3e-07 | 0.00011 | 0.077 (0.01) | 1.2e-07 | 0.00014 | 1.4e-06 | 0.01 |
| HTN | 0.11 (0.01) | 6.2e-29 | 2.3e-10 | 5.2e-14 | 0.067 (0.01) | 9.9e-09 | 0.72 | 2e-20 | 4.2e-11 |
| Average | 0.12 | | | | 0.091 | | | | |
| **African** | | | | | | | | | |
| LungCancer | -0.0087 (0.008) | 0.049 | 0.063 | 0.7 | 0.02 (0.03) | 0.53 | 0.52 | | 0.31 |
| BowelCancer | 0.0082 (0.02) | 0.65 | 0.75 | 0.27 | 0.0025 (0.008) | 0.78 | 0.96 | | 0.61 |
| Stroke | -0.0031 (0.008) | 0.7 | 0.69 | 0.78 | -0.003 (0.01) | 0.83 | 0.85 | 0.52 | 1 |
| COPD | 0.0022 (0.004) | 0.62 | 0.49 | 0.46 | 0.0033 (0.005) | 0.52 | 0.43 | | 0.71 |
| ProstateCancer | 0.069 (0.05) | 0.13 | 0.23 | 0.27 | 0.071 (0.06) | 0.18 | 0.3 | 0.11 | 0.91 |
| T2D | 0.16 (0.02) | 1e-14 | 1.3e-13 | 0.039 | 0.12 (0.02) | 6.5e-07 | 6e-06 | 0.0088 | 0.00074 |
| BreastCancer | 0 (0) | | | | 0.00026 (0.001) | 0.89 | 0.89 | | 0.89 |
| Depression | 0.072 (0.02) | 0.00011 | 9.1e-05 | 0.13 | 0.067 (0.03) | 0.0019 | 0.0017 | | 0.65 |
| CAD | 0.027 (0.01) | 0.027 | 0.035 | 0.22 | 0.0054 (0.01) | 0.71 | 0.88 | 0.1 | 0.012 |
| HTN | 0.085 (0.008) | 2.3e-27 | 3.4e-24 | 0.0041 | 0.055 (0.01) | 4.1e-07 | 2.3e-05 | 3.5e-05 | 5.3e-09 |
| Average | 0.11 | | | | 0.08 | | | | |

Supplementary Table 20: **Numerical results of analyses with covariates for 10 diseases from the UK Biobank.** The average is shown across the three well-powered traits (grey shading). The **(a)** raw and **(b)** relative performance of all prediction methods are shown (relative is relative to a model incorporating covariates alone across all diseases); PRS-$FH^{+}_{log}$ and PRS-$FH^{+}_{liab}$ are compared in **(c)**. Within **(c)** the p-value for the difference of PRS-FH versus FH is shown within the model of interest (e.g. PRS-$FH^{+}_{liab}$ vs $FH^{+}_{liab}$ and similarly for log; $H_0 : \Delta R_o = 0$). The three well-powered traits are highlighted with grey shading. The increases in prediction $R^2$ attained by $PRS^{+}$, $FH^{+}_{log}$, $FH^{+}_{liab}$, PRS-$FH^{+}_{log}$ and PRS-$FH^{+}_{liab}$ vs. a prediction model based on covariates alone were generally similar to the absolute predictive $R^2$ attained by PRS, $FH_{log}$, $FH_{liab}$, PRS-$FH_{log}$, and PRS-$FH_{liab}$ with limited exceptions. The exception to this trend is the predictive accuracy of $FH^{+}_{liab}$ (and likewise PRS-$FH^{+}_{liab}$) for HTN in South Asians and Africans. This could be due to a large correlation between the FH and covariate predictions for HTN within South Asians and Africans (Supplementary Table 21) and the inability of the liability-based methods to jointly estimate the effects of FH and covariates. The overall correlation between the $FH_{liab}$ and covariate prediction is 0.11 and 0.12 within South Asians and Africans for HTN, respectively. This strong correlation is driven primarily by the correlation of $FH_{liab}$ with age within both within South Asians and Africans (0.095 and 0.10, respectively) and with sex within Africans (0.096).

| Trait | PRS Covar | PRS Age | PRS Sex | PRS BMI | PRS PCs | $FH_{log}$ Covar | $FH_{log}$ Age | $FH_{log}$ Sex | $FH_{log}$ BMI | $FH_{log}$ PCs | $FH_{liab}$ Covar | $FH_{liab}$ Age | $FH_{liab}$ Sex | $FH_{liab}$ BMI | $FH_{liab}$ PCs |
|---|---|---|---|---|---|---|---|---|---|---|---|---|---|---|---|
| Non-British European | | | | | | | | | | | | | | | |
| LungCancer | -0.0054 | 0.00012 | 0.00089 | 0.017 | -0.016 | 0.12 | 0.094 | 0.016 | 0.05 | 0.082 | 0.13 | 0.11 | 0.021 | 0.064 | 0.087 |
| BowelCancer | -0.018 | 0.0014 | -0.0038 | 0.0017 | -0.057 | 0.12 | 0.074 | 0.013 | 0.031 | 0.14 | 0.11 | 0.062 | 0.015 | 0.029 | 0.14 |
| Stroke | 0.0068 | -0.01 | 0.0044 | 0.029 | -0.0011 | 0.13 | 0.1 | -0.0066 | 0.028 | 0.14 | 0.13 | 0.12 | -0.0054 | 0.029 | 0.1 |
| COPD | 0.05 | -0.0097 | 0.014 | 0.059 | 0.073 | 0.14 | 0.063 | -0.02 | 0.048 | 0.17 | 0.16 | 0.094 | -0.016 | 0.057 | 0.16 |
| ProstateCancer | 0.019 | 0.0045 | | 0.002 | 0.07 | 0.069 | 0.071 | | 0.0079 | 0.0075 | 0.074 | 0.075 | | 0.013 | 0.011 |
| T2D | 0.054 | -0.0013 | 0.013 | 0.057 | 0.09 | 0.061 | 0.017 | -0.011 | 0.074 | 0.049 | 0.081 | 0.047 | -0.002 | 0.079 | 0.046 |
| BreastCancer | -0.0084 | -0.011 | | -0.0068 | -0.016 | 0.06 | 0.055 | | 0.013 | 0.049 | 0.067 | 0.059 | | 0.016 | 0.05 |
| Depression | 0.028 | 0.01 | 0.0057 | 0.031 | 0.0079 | 0.055 | 0.041 | 0.057 | 0.0042 | 0.011 | 0.045 | 0.032 | 0.055 | 0.0055 | -0.014 |
| CAD | 0.021 | -0.015 | 0.0036 | 0.046 | 0.048 | 0.095 | 0.14 | -0.038 | 0.047 | 0.079 | 0.12 | 0.16 | -0.032 | 0.055 | 0.071 |
| HTN | 0.064 | -0.016 | 0.009 | 0.097 | 0.058 | 0.051 | 0.029 | -0.056 | 0.068 | 0.033 | 0.087 | 0.072 | -0.046 | 0.074 | 0.02 |
| South Asian | | | | | | | | | | | | | | | |
| LungCancer | -0.0099 | -0.0061 | -0.0097 | 0.0096 | 0.0031 | 0.019 | 0.065 | 0.029 | 0.054 | -0.0069 | 0.0085 | 0.059 | 0.012 | 0.045 | -0.011 |
| BowelCancer | -0.0024 | -0.0078 | 0.013 | 0.0013 | -0.00093 | -0.0096 | -0.0073 | 0.0073 | 0.031 | 0.0035 | -0.015 | -0.017 | 0.0013 | 0.023 | 0.0021 |
| Stroke | 0.011 | -0.018 | 0.0029 | 0.028 | 0.02 | 0.1 | 0.14 | 0.0043 | 0.029 | 0.012 | 0.086 | 0.13 | 0.0027 | 0.024 | 0.00072 |
| COPD | -0.012 | -0.026 | -0.017 | 0.017 | -0.007 | 0.081 | 0.076 | 0.0084 | 0.073 | -0.0054 | 0.041 | 0.028 | -0.0091 | 0.036 | 0.0035 |
| ProstateCancer | -0.0044 | 0.024 | | 0.01 | -0.02 | 0.021 | 0.014 | | 0.0086 | 0.023 | 0.03 | 0.028 | | -0.0017 | 0.018 |
| T2D | 0.0067 | -0.012 | -0.023 | -0.0014 | 0.095 | 0.087 | 0.06 | -0.0027 | 0.07 | 0.064 | 0.085 | 0.06 | 0.0052 | 0.063 | 0.06 |
| BreastCancer | -0.028 | -0.012 | | -0.003 | -0.027 | 0.016 | -0.0011 | | -0.066 | 0.045 | 0.028 | 0.029 | | -0.034 | 0.025 |
| Depression | -0.023 | 0.011 | 0.0012 | 0.013 | -0.072 | 0.045 | 0.026 | 0.035 | 0.029 | 0.02 | 0.034 | 0.033 | 0.027 | 0.023 | 0.011 |
| CAD | -0.017 | -0.034 | 0.0023 | 0.029 | -0.012 | 0.11 | 0.15 | -0.024 | 0.054 | 0.044 | 0.1 | 0.14 | -0.017 | 0.039 | 0.032 |
| HTN | -0.00098 | -0.035 | -0.015 | 0.057 | -0.0026 | 0.15 | 0.13 | -0.023 | 0.082 | 0.041 | 0.11 | 0.095 | -0.02 | 0.067 | 0.036 |
| African | | | | | | | | | | | | | | | |
| LungCancer | -0.025 | 0.0058 | -0.0099 | -0.045 | -0.043 | 0.01 | 0.029 | -0.014 | 0.039 | 0.0059 | 0.011 | 0.032 | -0.01 | 0.034 | 0.00058 |
| BowelCancer | -0.0074 | 0.0068 | -0.0051 | 0.012 | -0.012 | 0.0076 | 0.01 | 0.0069 | 0.014 | 0.012 | 0.01 | 0.0063 | 0.012 | 0.0036 | 0.017 |
| Stroke | 0.034 | 0.0096 | -0.0035 | 0.022 | 0.055 | 0.087 | 0.12 | -0.0058 | 0.031 | 0.0098 | 0.082 | 0.11 | -0.0091 | 0.019 | 0.0074 |
| COPD | -0.057 | -0.0033 | -0.0092 | 0.035 | -0.075 | -0.011 | -0.016 | 0.021 | 0.00031 | 0.0087 | 0.0046 | -0.0013 | 0.023 | 0.013 | 0.0049 |
| ProstateCancer | 0.036 | -0.008 | | -0.034 | 0.13 | 0.13 | 0.12 | | 0.023 | 0.021 | 0.14 | 0.13 | | 0.022 | 0.028 |
| T2D | -0.0057 | -0.014 | -0.015 | 0.045 | -0.033 | 0.058 | 0.067 | -0.072 | 0.084 | -0.0046 | 0.073 | 0.096 | -0.065 | 0.071 | -0.0072 |
| BreastCancer | -0.011 | 0.014 | | -0.0032 | -0.022 | 0.0075 | 0.036 | | 0.017 | -0.0032 | 0.018 | 0.049 | | 0.014 | -0.0028 |
| Depression | -0.036 | -0.0068 | 0.013 | 0.0088 | -0.058 | 0.069 | 0.082 | 0.062 | 0.02 | 0.019 | 0.071 | 0.064 | 0.053 | 0.03 | 0.023 |
| CAD | -0.013 | -0.0087 | -0.013 | 0.021 | -0.0066 | 0.068 | 0.055 | -0.04 | 0.066 | 0.0073 | 0.056 | 0.067 | -0.052 | 0.057 | -0.011 |
| HTN | 0.014 | -0.02 | -0.003 | 0.031 | 0.021 | 0.11 | 0.078 | 0.095 | 0.086 | 0.04 | 0.12 | 0.1 | 0.096 | 0.079 | 0.039 |

Supplementary Table 21: **Correlations between PRS and FH predictions vs. predictions using covariates alone for 10 diseases from the UK Biobank.**

| Trait | Covariates | $PRS^+$ | $FH^+_{log}$ | $FH^+_{liab}$ | $PRS\text{-}FH^+_{log}$ | $PRS\text{-}FH^+_{liab}$ |
|---|---|---|---|---|---|---|
| Non-British European | | | | | | |
| LungCancer | 0.74 | 0.74 | 0.7 | 0.67 | 0.71 | 0.69 |
| BowelCancer | 0.82 | 0.83 | 0.8 | 0.76 | 0.81 | 0.78 |
| Stroke | 0.93 | 0.92 | 0.89 | 0.75 | 0.89 | 0.75 |
| COPD | 0.96 | 0.95 | 0.94 | 0.82 | 0.93 | 0.81 |
| ProstateCancer | 0.88 | 0.84 | 0.87 | 0.75 | 0.85 | 0.77 |
| T2D | 0.87 | 0.91 | 0.9 | 0.78 | 0.93 | 0.79 |
| BreastCancer | 0.88 | 0.84 | 0.9 | 0.79 | 0.85 | 0.78 |
| Depression | 0.95 | 0.96 | 0.93 | 0.64 | 0.93 | 0.65 |
| CAD | 0.96 | 0.98 | 0.98 | 0.87 | 0.99 | 0.89 |
| HTN | 0.99 | 1 | 1 | 0.87 | 1 | 0.89 |
| Average | 0.94 | 0.96 | 0.94 | 0.76 | 0.95 | 0.78 |
| South Asian | | | | | | |
| LungCancer | -0.027 | -0.041 | -0.047 | -0.025 | -0.051 | -0.025 |
| BowelCancer | 0.094 | 0.11 | 0.22 | 0.26 | 0.22 | 0.26 |
| Stroke | 0.7 | 0.69 | 0.68 | 0.67 | 0.67 | 0.67 |
| COPD | 0.63 | 0.64 | 0.54 | 0.2 | 0.53 | 0.2 |
| ProstateCancer | 0.37 | 0.41 | 0.32 | 0.34 | 0.4 | 0.39 |
| T2D | 0.91 | 0.93 | 0.93 | 0.74 | 0.94 | 0.77 |
| BreastCancer | -0.24 | 0.019 | 0.077 | 0.096 | 0.14 | 0.18 |
| Depression | 0.64 | 0.68 | 0.77 | 0.39 | 0.79 | 0.4 |
| CAD | 0.93 | 0.94 | 0.94 | 0.88 | 0.94 | 0.9 |
| HTN | 0.98 | 0.98 | 0.98 | 0.82 | 0.98 | 0.85 |
| Average | 0.84 | 0.86 | 0.9 | 0.65 | 0.91 | 0.67 |
| African | | | | | | |
| LungCancer | 0.14 | 0.13 | 0.062 | 0.16 | 0.057 | 0.16 |
| BowelCancer | 0.3 | 0.32 | 0.27 | 0.26 | 0.31 | 0.26 |
| Stroke | 0.71 | 0.7 | 0.66 | 0.55 | 0.65 | 0.54 |
| COPD | 0.24 | 0.22 | 0.24 | 0.25 | 0.23 | 0.25 |
| ProstateCancer | 0.79 | 0.79 | 0.76 | 0.65 | 0.77 | 0.66 |
| T2D | 0.86 | 0.86 | 0.91 | 0.7 | 0.91 | 0.71 |
| BreastCancer | -0.13 | -0.12 | -0.051 | 0.052 | -0.047 | 0.052 |
| Depression | 0.65 | 0.64 | 0.82 | 0.33 | 0.82 | 0.33 |
| CAD | 0.86 | 0.85 | 0.85 | 0.74 | 0.86 | 0.75 |
| HTN | 0.98 | 0.98 | 0.98 | 0.76 | 0.98 | 0.77 |
| Average | 0.83 | 0.83 | 0.91 | 0.6 | 0.9 | 0.6 |
| Average | 0.87 | 0.88 | 0.91 | 0.67 | 0.92 | 0.68 |

Supplementary Table 22: **Calibration results of analyses with covariates for 10 diseases from the UK Biobank.** We report the slope from the regression of disease status on predictions for each prediction method. The average is shown across the three well-powered traits (grey shading).

| | PRS-FH$^{+}_{log}$ | | | | PRS-FH$^{+}_{liab}$ | | | |
|---|---|---|---|---|---|---|---|---|
| Trait | All | No-sib | Δ | Δp | All | No-sib | Δ | Δp |
| **Non-British European** | | | | | | | | |
| LungCancer | 0.014 | 0.0046 | 0.0097 | 0.31 | 0.017 | 0.0055 | 0.012 | 0.19 |
| BowelCancer | 0.011 | 0.015 | -0.0035 | 0.0011 | 0.0098 | 0.014 | -0.0044 | 0.11 |
| Stroke | 0.0084 | 0.00077 | 0.0076 | 0.15 | 0.0017 | -0.00025 | 0.002 | 0.8 |
| COPD | 0.088 | 0.053 | 0.035 | 2.8e-05 | 0.093 | 0.055 | 0.038 | 7.9e-07 |
| ProstateCancer | 0.13 | 0.11 | 0.018 | 0.15 | 0.12 | 0.11 | 0.014 | 0.48 |
| T2D | 0.25 | 0.2 | 0.048 | 3.4e-05 | 0.22 | 0.18 | 0.038 | 0.019 |
| BreastCancer | 0.036 | 0.034 | 0.0024 | 0.14 | 0.036 | 0.034 | 0.0018 | 0.58 |
| Depression | 0.067 | 0.056 | 0.011 | 0.00044 | 0.061 | 0.057 | 0.0045 | 0.38 |
| CAD | 0.11 | 0.088 | 0.02 | 3.5e-06 | 0.096 | 0.083 | 0.013 | 0.02 |
| HTN | 0.14 | 0.12 | 0.016 | 6.3e-23 | 0.11 | 0.11 | -0.00034 | 0.93 |
| **South Asian** | | | | | | | | |
| LungCancer | 0 | 0 | 0 | | 0 | 0 | 0 | |
| BowelCancer | 0.0083 | 0.003 | 0.0053 | 0.39 | 0.018 | 0.0017 | 0.016 | 0.21 |
| Stroke | 0.00047 | -0.007 | 0.0075 | 0.39 | 0.0092 | -0.0036 | 0.013 | 0.17 |
| COPD | -0.0011 | 0.0071 | -0.0082 | 0.17 | -0.03 | 0.011 | -0.041 | 0.023 |
| ProstateCancer | 0.056 | 0.049 | 0.0064 | 0.65 | 0.069 | 0.041 | 0.028 | 0.48 |
| T2D | 0.16 | 0.12 | 0.042 | 1.5e-06 | 0.13 | 0.11 | 0.02 | 0.21 |
| BreastCancer | 0.0025 | 0.00033 | 0.0022 | 0.12 | 0.0039 | 0.00071 | 0.0032 | 0.11 |
| Depression | 0.08 | 0.053 | 0.027 | 0.023 | 0.07 | 0.055 | 0.015 | 0.47 |
| CAD | 0.09 | 0.069 | 0.021 | 0.0031 | 0.077 | 0.067 | 0.0096 | 0.31 |
| HTN | 0.11 | 0.095 | 0.015 | 0.00036 | 0.067 | 0.083 | -0.015 | 0.12 |
| **African** | | | | | | | | |
| LungCancer | -0.0087 | -0.0067 | -0.0021 | 0.53 | 0.02 | -0.0017 | 0.022 | 0.49 |
| BowelCancer | 0.0082 | -0.0026 | 0.011 | 0.47 | 0.0025 | -0.0016 | 0.0041 | 0.65 |
| Stroke | -0.0031 | -0.0041 | 0.001 | 0.89 | -0.003 | -0.0059 | 0.0029 | 0.85 |
| COPD | 0.0022 | -0.0011 | 0.0033 | 0.26 | 0.0033 | 0.00043 | 0.0029 | 0.49 |
| ProstateCancer | 0.069 | 0.037 | 0.032 | 0.4 | 0.071 | 0.043 | 0.028 | 0.59 |
| T2D | 0.16 | 0.11 | 0.045 | 0.00036 | 0.12 | 0.1 | 0.017 | 0.41 |
| BreastCancer | 0 | 0 | 0 | | 0.00026 | 0 | 0.00026 | 0.89 |
| Depression | 0.072 | 0.04 | 0.032 | 0.031 | 0.067 | 0.041 | 0.026 | 0.26 |
| CAD | 0.027 | 0.028 | -0.00027 | 0.95 | 0.0054 | 0.025 | -0.019 | 0.051 |
| HTN | 0.085 | 0.051 | 0.034 | 5.6e-11 | 0.055 | 0.05 | 0.0059 | 0.54 |

Supplementary Table 23: **Results of PRS-FH methods using parental history only in analyses with covariates for 10 diseases from the UK Biobank.** We report the relative prediction $R^2_l$ for both PRS-FH prediction methods (relative to a model with covariates alone) including parental and sibling history (All) and parental history only (No-sib), as well as the difference in prediction $R^2_l$ (Δ) and the jackknife for the difference ($H_0 : \Delta R_o = 0$). The effect of incorporating sibling history on the predictive accuracy of PRS-FH$^{+}_{liab}$ is less clear, this is in contrast to the clear benefit when covariates were not being modeled (Supplementary Table 13 vs. 23)

| Trait | PRS-FH$^{+}_{log}$ | PRS+$I(FH \geq 1)^{+}_{log}$ | $\Delta$ | $\Delta$p |
|---|---|---|---|---|
| Non-British European | | | | |
| LungCancer | 0.014 | 0.021 | -0.0067 | 0.22 |
| BowelCancer | 0.011 | 0.013 | -0.0022 | 0.45 |
| Stroke | 0.0084 | 0.0046 | 0.0038 | 0.45 |
| COPD | 0.088 | 0.063 | 0.025 | 0.0023 |
| ProstateCancer | 0.13 | 0.13 | 0.0017 | 0.81 |
| T2D | 0.25 | 0.22 | 0.03 | 0.0011 |
| BreastCancer | 0.036 | 0.035 | 0.00087 | 0.44 |
| Depression | 0.067 | 0.063 | 0.0035 | 0.34 |
| CAD | 0.11 | 0.085 | 0.024 | 2.1e-07 |
| HTN | 0.14 | 0.12 | 0.014 | 9.4e-15 |
| South Asian | | | | |
| LungCancer | 0 | 0 | 0 | |
| BowelCancer | 0.0083 | 0.0057 | 0.0026 | 0.72 |
| Stroke | 0.00047 | 0.0011 | -0.00063 | 0.93 |
| COPD | -0.0011 | 0.0036 | -0.0048 | 0.51 |
| ProstateCancer | 0.056 | 0.039 | 0.017 | 0.3 |
| T2D | 0.16 | 0.14 | 0.024 | 0.018 |
| BreastCancer | 0.0025 | 0.0018 | 0.00069 | 0.63 |
| Depression | 0.08 | 0.081 | -0.0016 | 0.91 |
| CAD | 0.09 | 0.073 | 0.017 | 0.034 |
| HTN | 0.11 | 0.093 | 0.016 | 6e-05 |
| African | | | | |
| LungCancer | -0.0087 | -0.0015 | -0.0073 | 0.16 |
| BowelCancer | 0.0082 | 0.0081 | 0.00018 | 0.98 |
| Stroke | -0.0031 | -0.004 | 0.00085 | 0.9 |
| COPD | 0.0022 | -0.00069 | 0.0029 | 0.33 |
| ProstateCancer | 0.069 | 0.075 | -0.0065 | 0.77 |
| T2D | 0.16 | 0.11 | 0.053 | 6.8e-05 |
| BreastCancer | 0 | 3.8e-06 | -3.8e-06 | 0.016 |
| Depression | 0.072 | 0.074 | -0.0016 | 0.92 |
| CAD | 0.027 | 0.024 | 0.0036 | 0.6 |
| HTN | 0.085 | 0.041 | 0.045 | 1.9e-15 |

Supplementary Table 24: **Results of PRS-FH$_{log}$ versus a simplified logistic regression-based method which used an indicator variable for presence of family history in analyses with covariates for 10 diseases from the UK Biobank.** We report the relative prediction $R^2_l$ (relative to a model with covariates alone) for the PRS-FH$^{+}_{log}$ prediction method (distinct independent predictors for maternal, paternal, and sibling history) and the simplified logistic regression-based method (one predictor for a family history of disease; PRS+$I(FH \geq 1)^{+}_{log}$ , as well as the difference in prediction $R^2_l$ ($\Delta$) and the jackknife p-value for the difference ($H_0 : \Delta R_o = 0$). $I(FH \geq 1) = 1$ if an individual reported either their mother, father, or sibling was affected; $I(FH \geq 1) = 0$ if an individual reported both mother and father were unaffected and either reported 0 relevant siblings or that 0 of their siblings were affected; otherwise $I(FH \geq 1) = NA$. Individuals with $I(FH \geq 1) = NA$ were assigned the mean $I(FH \geq 1)$ across all individuals in the 9 training folds.

| Scenario | $h_l^2$ mean | s.e. | Average pseudo mean | s.e. | Diff. pseudo mean | s.e. |
|---|---|---|---|---|---|---|
| K=5% | | | FH | | | |
| No correlation | 0.0526 | 0.00333 | 0.0521 | 0.00337 | 0.0517 | 0.00335 |
| Correlation | 0.167 | 0.00434 | 0.167 | 0.00464 | 0.167 | 0.00454 |
| Differential correlation | 0.173 | 0.00512 | 0.173 | 0.00518 | 0.183 | 0.00524 |
| | | | PRS-FH$_{liab}$ | | | |
| No correlation | 0.104 | 0.00494 | 0.104 | 0.00498 | 0.103 | 0.00496 |
| Correlation | 0.201 | 0.00295 | 0.219 | 0.00415 | 0.218 | 0.00408 |
| Differential correlation | 0.206 | 0.00428 | 0.222 | 0.00584 | 0.236 | 0.00534 |
| K=25% | | | FH | | | |
| No correlation | 0.0755 | 0.00135 | 0.0753 | 0.00136 | 0.0753 | 0.00134 |
| Correlation | 0.187 | 0.00217 | 0.188 | 0.00218 | 0.187 | 0.00216 |
| Differential correlation | 0.188 | 0.00138 | 0.188 | 0.00139 | 0.195 | 0.00137 |
| | | | PRS-FH$_{liab}$ | | | |
| No correlation | 0.167 | 0.00104 | 0.166 | 0.00103 | 0.166 | 0.00103 |
| Correlation | 0.245 | 0.00213 | 0.263 | 0.00274 | 0.263 | 0.00272 |
| Differential correlation | 0.248 | 0.00126 | 0.262 | 0.00168 | 0.273 | 0.00159 |

Supplementary Table 25: **Results of simulations of PRS-FH$_{liab}$ methods using pseudo-heritability, versus heritability, in settings with and without environmental correlation.** For each prediction method we report the mean $R_l^2$ across the 10 simulation replicates as well as the standard error of this mean.

| Training sample size | Number of independent training sample sets per fold Non-British European | South Asian / African |
|---|---|---|
| 35K | 1 | N/A |
| 20K | 2 | N/A |
| 10K | 4 | N/A |
| 6K | N/A | 1 |
| 5K | 8 | 2 |
| 2.5K | 16 | 4 |
| 1K | 40 | 10 |
| 0.5K | 80 | 20 |

Supplementary Table 26: **Number and size of down-sampled independent training sample sets for PRS-FH methods.** We report the size, and number, of independent training sample sets constructed for each of the 10-folds when decreasing the number of training samples. Total sample sizes for T2D, Depression, and HTN for Non-British European was 41642, 41842, and 41842, for South Asian was 6881, 7048, and 7048, and for African 6961, 7087, and 7087, respectively.

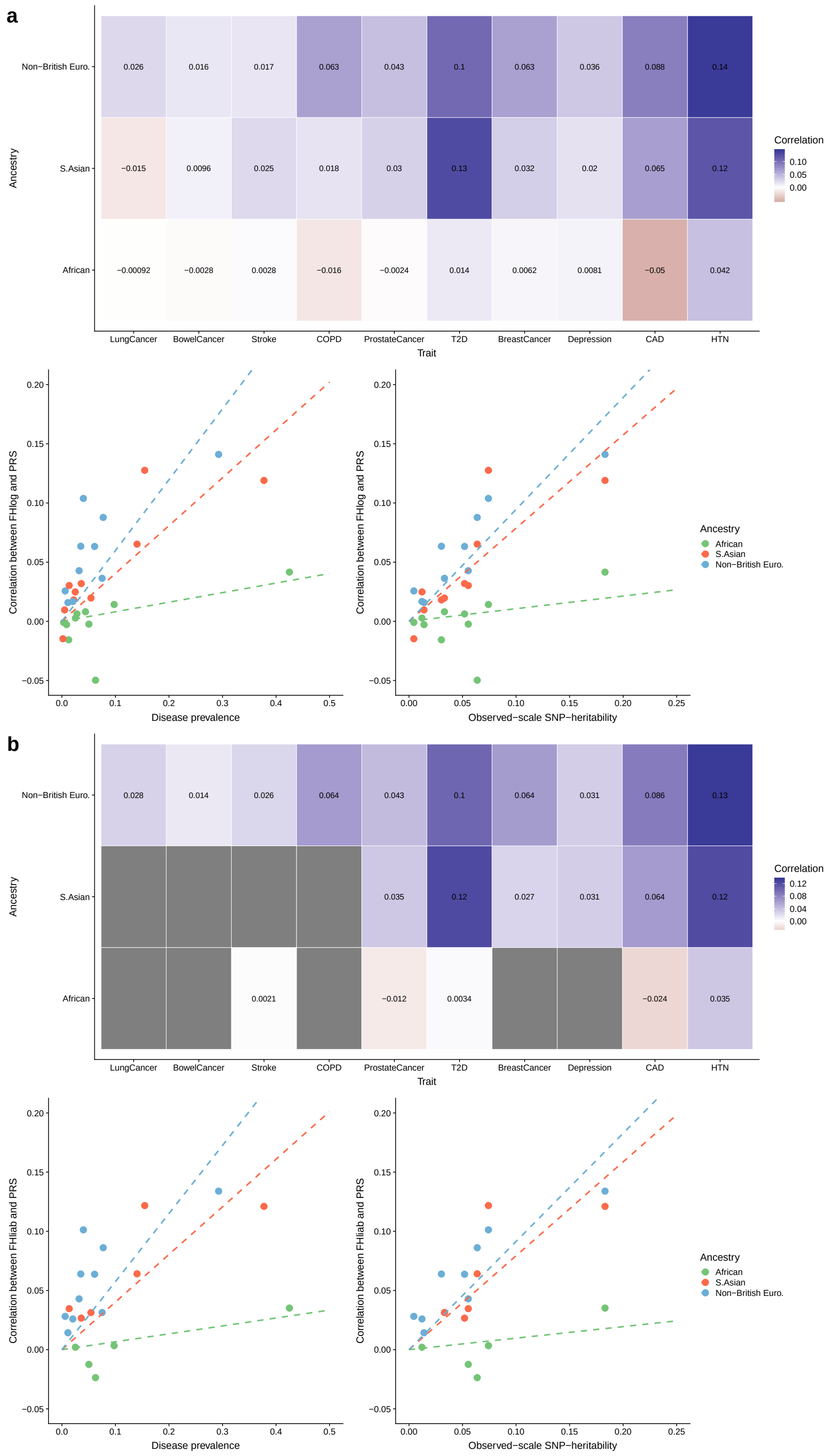


Supplementary Figure 1: **Correlations between PRS vs. FH predictions for 10 diseases from the UK Biobank.** The correlation between the PRS and (a) $FH_{log}$ and (b) $FH_{liab}$ predictors is shown and reported within the heatmaps; a grey box within panel (b) denotes the permutation test of $H_0 : V = 0$ failed to be rejected. Dot plots show the relationship between PRS-FH predictor correlations ((a) $FH_{log}$ and (b) $FH_{liab}$) as a function of disease prevalence as well as observed-scale heritability; lines represent a linear regression fit with an intercept of 0 fitted separately within the three testing sets.

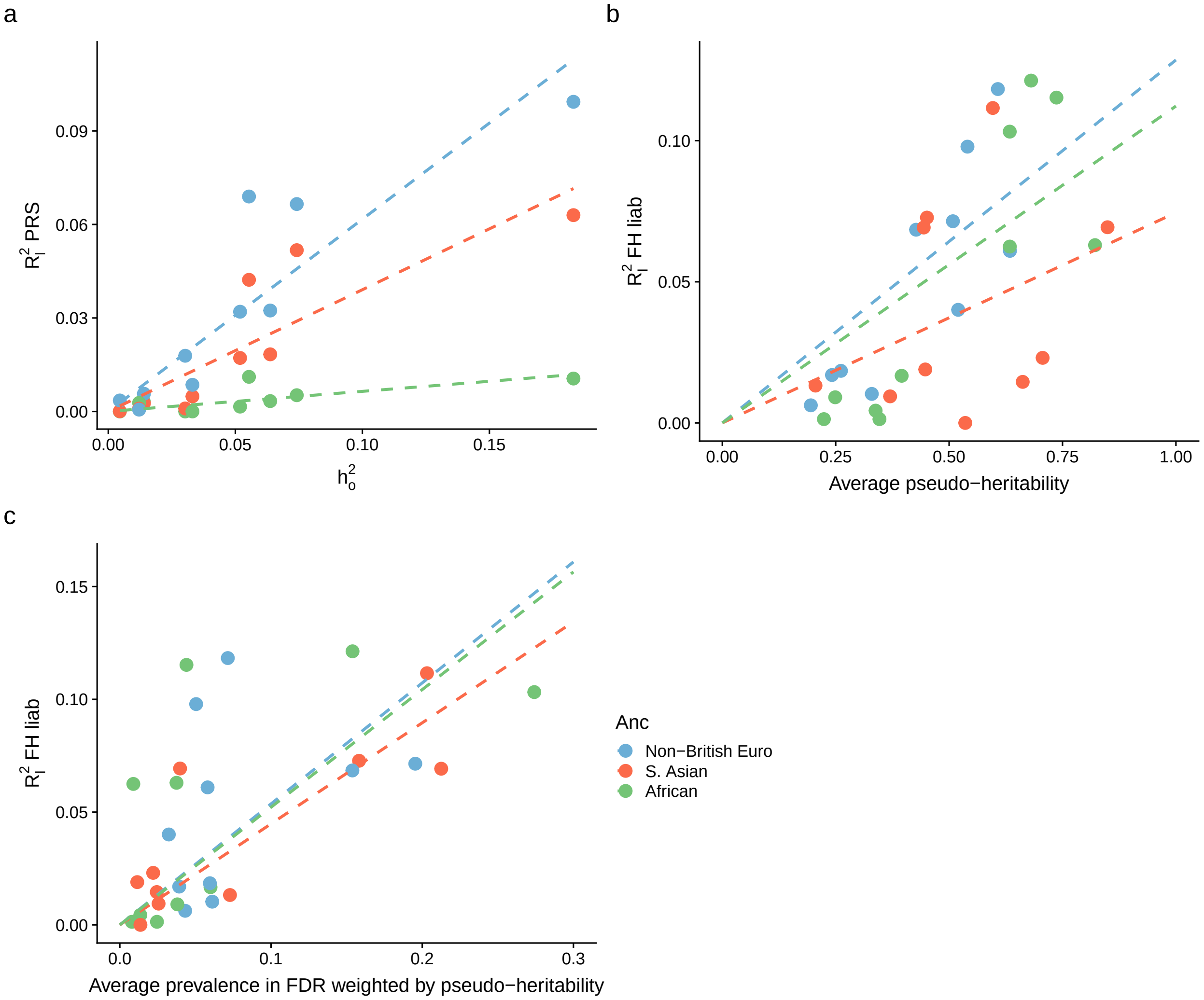


Supplementary Figure 2: **Impact of observed-scale heritability on accuracy of PRS and impact of pseudo-heritability and weighted relative disease prevalence on accuracy of FH$_{liab}$ for 10 diseases from the UK Biobank.** The relationship between (a) the predictive accuracy of PRS and the observed-scale heritability and the relationship between the predictive accuracy of FH$_{liab}$ and (b) the average pseudo-heritability (average covariance between liabilities of target samples and first-degree relatives) or (c) the weighted average disease prevalence in relatives; lines represent a linear regression fit with an intercept of 0 fitted separately within the three testing sets. We show the weighted average disease prevalence in relatives (weighted by pseudo-heritability) to show the prevalence among the relative's which predominantly contribute to the FH$_{liab}$ predictor. Due to similarity between FH$_{log}$ and FH$_{liab}$ when covariates are not included, only FH$_{liab}$ is shown. FDR: first degree relatives.

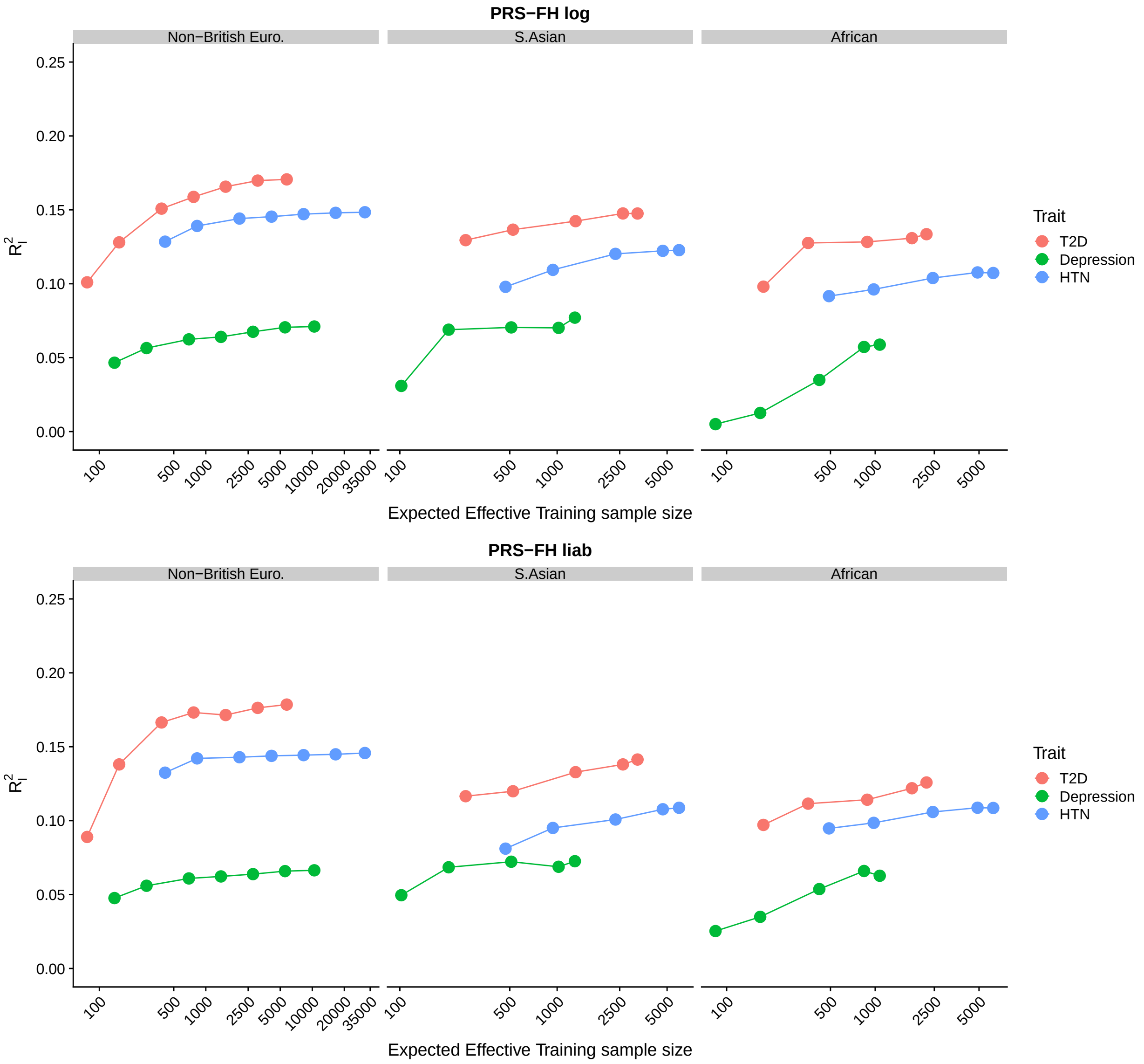


Supplementary Figure 3: **Impact of decreasing the number of training samples from the target population used to fit PRS-FH model parameters for 3 well-powered diseases from the UK Biobank.** We decreased the number of training samples from the target population used to fit PRS-FH model parameters. For each of the 10 folds, the remaining individuals from the other 9 folds are down-sampled to form PRS-FH training sets of various sizes (Supplementary Table 24) and the average prediction $R_l^2$ is then computed across training sizes. The effective sample size ($N_{eff}$) may vary as a function of the number of *cases* sampled within a training sample set; the expected $N_{eff}$ is computed using the overall disease prevalence and the number of training samples.

**a**

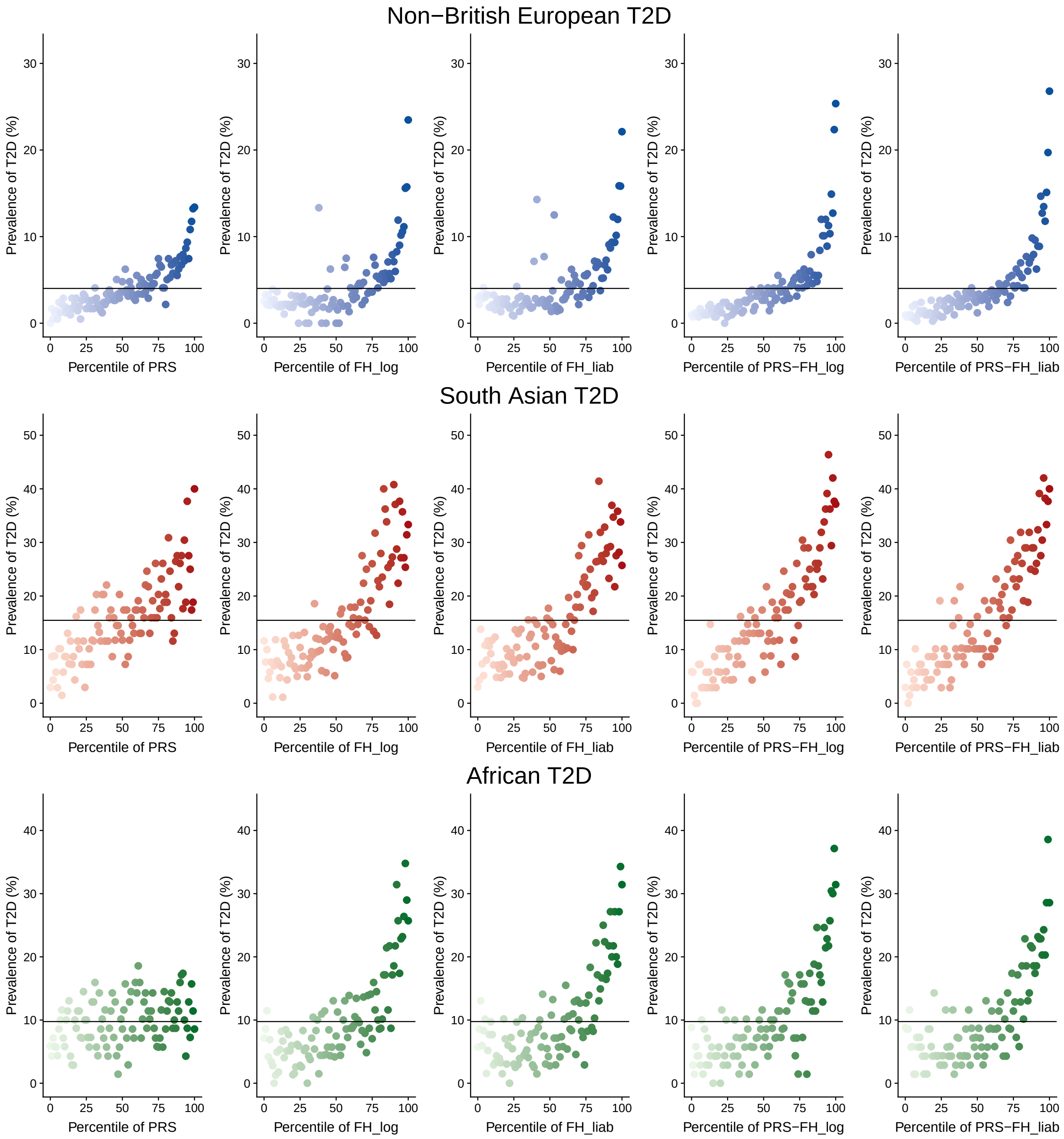

Non−British European T2D
South Asian T2D
African T2D
Prevalence of T2D (%)
Percentile of PRS
Percentile of FH_log
Percentile of FH_liab
Percentile of PRS−FH_log
Percentile of PRS−FH_liab

**b**

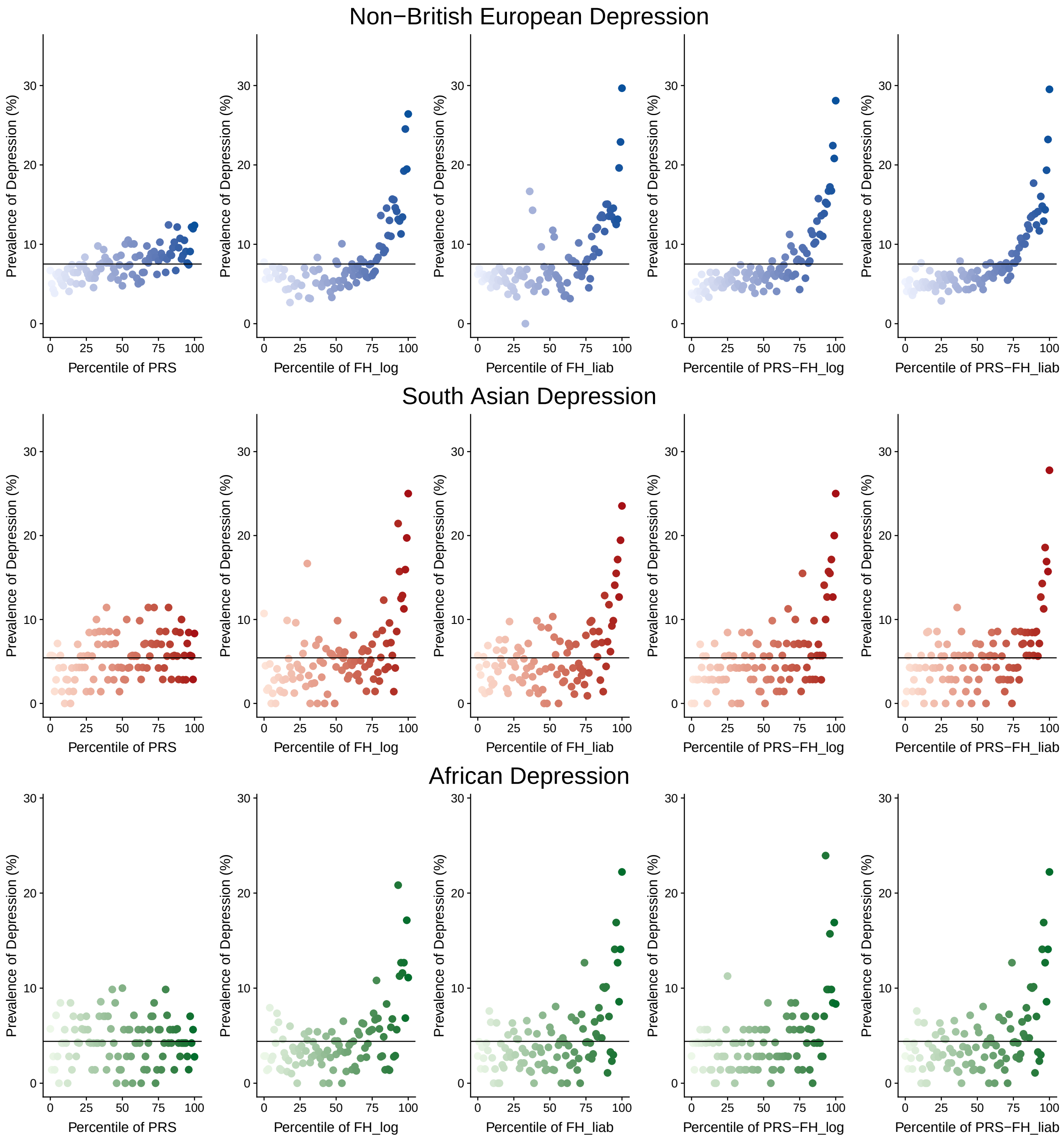
Non−British European Depression
South Asian Depression
African Depression
Prevalence of Depression (%)
Percentile of PRS
Percentile of FH_log
Percentile of FH_liab
Percentile of PRS−FH_log
Percentile of PRS−FH_liab

**c**

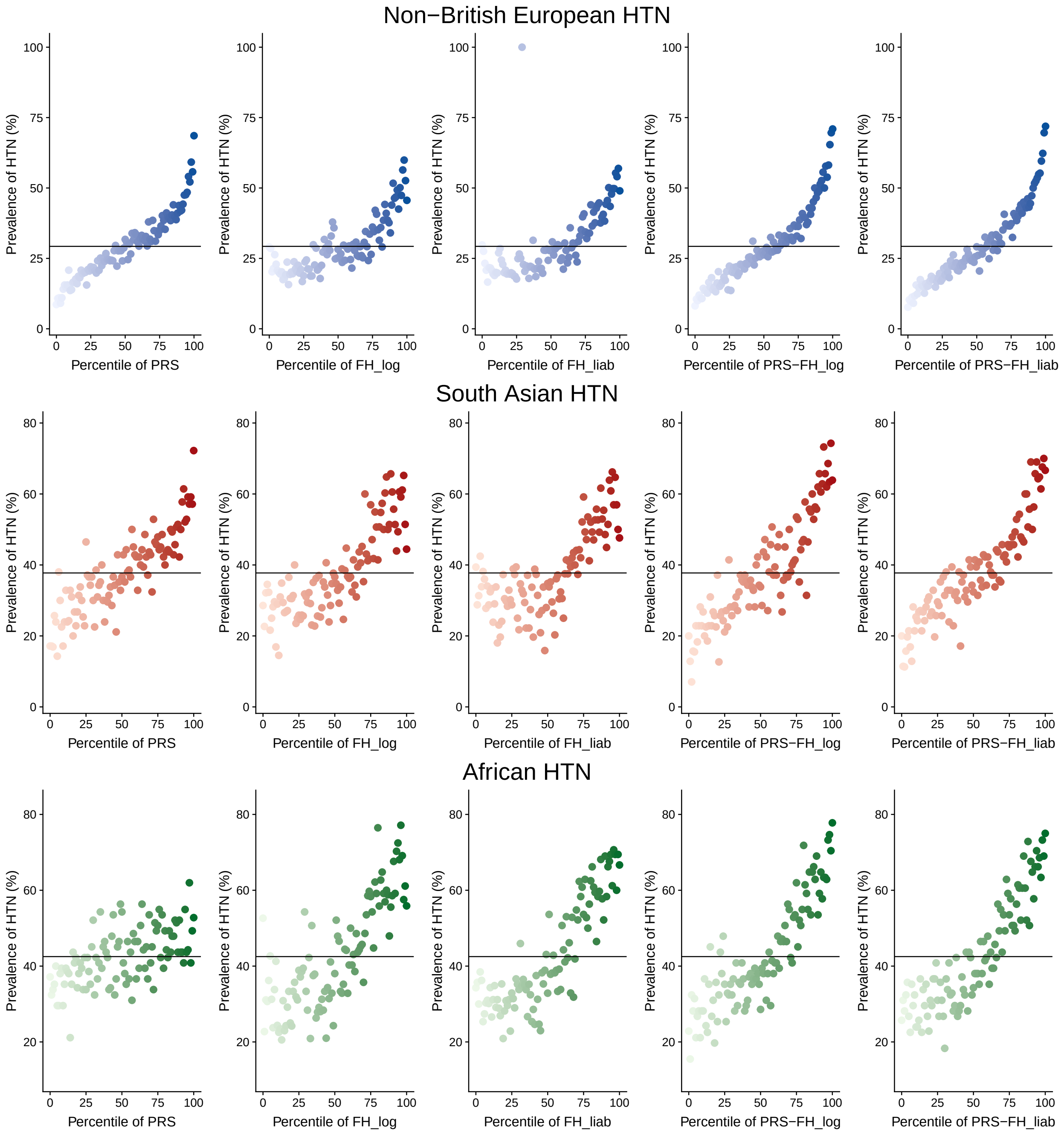


Supplementary Figure 4: **Prevalence of disease in each percentile of predicted disease risk for 3 well-powered diseases from the UK Biobank.** For each percentile of predicted disease risk for 5 main methods, the prevalence of (a) T2D, (b) Depression, and (c) HTN, is shown.
